# Targeting tRNA-Synthetase Interactions towards Novel Therapeutic Discovery Against Eukaryotic Pathogens

**DOI:** 10.1101/711697

**Authors:** Paul Kelly, Fatemeh Hadi-Nezhad, Dennis Liu, Travis J. Lawrence, Roger G. Linington, Michael Ibba, David H. Ardell

**Author notes:** Co-corresponding authors and persons to whom requests should be addressed:* Michael Ibba, PhD, Department of Microbiology, The Ohio State University, 318 West 12^th^ Avenue, Columbus, Ohio 43210, David H. Ardell, PhD, Department of Molecular and Cell Biology, University of California, 5200 North Lake Road, Merced, California 95343.

## Abstract

The development of chemotherapies against eukaryotic pathogens is especially challenging because of both the evolutionary conservation of drug targets between host and parasite, and the evolution of strain-dependent drug resistance. There is a strong need for new nontoxic drugs with broad-spectrum activity against trypanosome parasites such as *Leishmania* and *Trypanosoma.* A relatively untested approach is to target macromolecular interactions in parasites rather than small molecular interactions, under the hypothesis that the features specifying macromolecular interactions diverge more rapidly through coevolution. We computed tRNA Class-Informative Features in humans and eight clades of trypanosomes, identifying parasite-specific informative features (including base-pairs and base mis-pairs) that are broadly conserved over approximately 250 million years of trypanosome evolution. Validating these observations, we demonstrated biochemically that tRNA:aminoacyl-tRNA synthetase interactions are a promising target for anti-trypanosomal drug discovery. From a marine natural products extract library, we identified several fractions with inhibitory activity toward *Leishmania major* alanyl-tRNA synthetase (AlaRS) but no activity against the human homolog. These marine natural products extracts showed cross-reactivity towards *Trypanosoma cruzi* AlaRS indicating the broad-spectrum potential of our network predictions. These findings support a systems biology model in which combination chemotherapies that target multiple tRNA-synthetase interactions should be comparatively less prone to the emergence of resistance than conventional single drug therapies.

**Author Summary:** Trypanosome parasites pose a significant health risk worldwide. Conventional drug development strategies have proven challenging given the high conservation between humans and pathogens, with off-target toxicity being a common problem. Protein synthesis inhibitors have historically been an attractive target for antimicrobial discovery against bacteria, and more recently for eukaryotic pathogens. Here we propose that exploiting pathogen-specific tRNA-synthetase interactions offers the potential for highly targeted drug discovery. To this end, we improved tRNA gene annotations in trypanosome genomes, identified functionally informative trypanosome-specific tRNA features, and showed that these features are highly conserved over approximately 250 million years of trypanosome evolution. Highlighting the species-specific and broad-spectrum potential of our approach, we identified natural product inhibitors against the parasite translational machinery that have no effect on the homologous human enzyme.

## Introduction

Developing therapies against eukaryotic pathogens has proven challenging due to high conservation between the infectious agent drug target and their host counterpart [1]. Of particular concern is the trypanosome parasite *Leishmania* that infects upwards of 2 million individuals every year and accounts for more than 50,000 deaths annually [2]. While current treatments of amphotericin B and miltefosine are commonly prescribed to patients with leishmanial infections, they have undesired off-target cytotoxicity, leading to poor patient compliance and low-dose administration [3], and ultimately contributing to the rise of strain-dependent drug resistance [4,5]. There is a strong need for new nontoxic drugs with broad-spectrum activity against different species of *Leishmania* and other trypanosomes [6,7].

Given their essential role in protein synthesis, aminoacyl-tRNA synthetases (aaRSs) have been an attractive target for antimicrobial therapeutics [8]. AaRSs are essential enzymes found in all domains of life that are responsible for the correct pairing of free amino acids in the cell to their cognate tRNA [9]. AaRSs perform their activity in two steps: first, free amino acid is activated by the enzyme through the hydrolysis of ATP, forming an aminoacyl-adenylate. Second, the amino acid is transferred to its corresponding tRNA before being released into the aminoacyl-tRNA pool [9]. Given the complex pool of free amino acids and uncharged tRNAs in the cell, aaRSs have co-evolved discrete mechanisms to ensure mutually exclusive amino acid activation and cognate tRNA recognition [10]. The sequence/structural determinants (or anti-determinants) that lead to accurate aaRS-tRNA recognition are also known as the tRNA identity elements. The primary tRNA identity elements that aid in cognate aminoacylation have been extensively studied for several decades [11,12]. For example, across all three domains of life, all tRNA^Ala^ isoacceptors contain a conserved G:U base pair in the acceptor stem that is recognized by alanyl-tRNA synthetase (AlaRS), leading to accurate Ala-tRNA^Ala^ synthesis in the cell [13-15].

While some aaRS inhibitors have successfully made it to the clinic, including the IleRS-targeting mupirocin [16], ProRS inhibitor halofuginone [17], and the LeuRS inhibitor tavaborole [18,19], there are likely many potential aaRS drugs still to be identified. Target-based approaches relying on structural data and sequence identity have previously been used to try and predict novel trypanosome aaRS drug targets [20-22] with some recent success [23]. While structure-based approaches have their utility, exploiting tRNA-aaRS interactions has been under-explored for its therapeutic potential. In particular, while interactions with small molecules are expected to be quite conserved across species, the evolutionary diversification of tRNA identity element interactions through coevolution with aaRSs opens the possibility of greater species-specific inhibition.

While major identity elements have been experimentally characterized for many aaRS-tRNA pairs in various model systems, much less is known about how tRNA identity elements evolve and diverge over the Tree of Life. Recent theoretical advances explain how tRNA identity elements can evolve and diverge in a phylogenetically informative way, even while under strong selective constraints [24]. In earlier work, we developed a bioinformatic method to predict tRNA identity elements [25]. Our bioinformatic predictions are called Class-Informative Features (CIFs), based on the statistic of structure-conditioned Shannon Information [26], and visualized through graphs called Function Logos [25]. In later work, we applied two other statistics, Information Difference and Kullback-Leibler Divergence, to facilitate pairwise comparisons of CIFs between two taxa, in two new visualizations called Information Difference (ID) logos and Kullback-Leibler Divergence (KLD) logos, respectively [27]. ID logos visualize gains and losses of CIFs, while KLD logos visualize the functional conversion of CIFs from one functional type of tRNA to another. In the present work, we integrate together all three statistics (structure-conditioned information about function, ID, and KLD) and apply it to the problem of identifying parasite-specific tRNA identity elements. Our approach visualizes functionally informative CIFs in parasites that have either gained or retained functional information relative to humans, altered functional associations, or both, since divergence from their common ancestor with humans.

Our modeling approach integrates genomic tRNA sequence variation across multiple tRNA gene families of different functions, revealing potentially useful information about the specification of substrate identity for all aaRSs simultaneously. The multiplicity of aaRSs in cells provides multiple potential targets for inhibition of essential parasite enzymes, opening the door to improved combination chemotherapies. Advances in systems biology and chemogenomics have fueled interest in combination chemotherapies to benefit from synergistic drug interactions [28-31] and combat the evolution of resistance [32]. Combination chemotherapies are naturally effective, for example, in the pathogenic defenses of arthropods [33] and have yielded exciting antifungal [34] and antihelminthic [35] therapies. Additionally, artemisinin-based combination therapies (ACTs) are the primary treatment plan for *Plasmodium falciparum* malaria infections [36,37].

Here, we report our improved annotation of TriTrypDB genomes and new methodologies for predicting conserved identity elements across biological domains. As proof of principle, we screened for identity element divergence between trypanosomes and humans to search for new therapeutic targets for these eukaryotic pathogens. Validating our computational approaches, we found several natural products that inhibit *Leishmania major* AlaRS activity but have no effect on the homologous human enzyme. The compounds we identified also have inhibitory activity against *Trypanosoma cruzi* AlaRS, showing that our approach holds promise towards identifying new broad-spectrum anti-trypanosomal therapies.

## Methods

### Annotation, Filtering and Alignment of tRNA Genes in TriTrypDB Genomes

We downloaded data for 46 genomes from TriTrybDB version 41 released December 5^th^, 2018. We ran tRNAscan-SE v.2.0 [38] and Aragorn v.1.2.38 [39] using option “-i116” (implying a maximum intron length in search targets of 116 base-pairs) on this data. We unified gene records from the two finders if they overlapped by at least one base-pair, verifying that the orientations were consistent and end-displacements were less than or equal to 4 bp. To independently identify initiator tRNA genes, we computed edit distances [40] of CAT-anticodon-containing genes and clustered them using Ward.D2 method [41], examining each cluster for initiator-distinguishing features described in [42].

To further investigate these gene records, we examined their genetic clustering in TriTrypDB genomes as defined by co-occurrence within a distance of 1000 bp on either strand. We computed similarities of tRNA gene-clusters by clustering them according to pairwise Jaccard Index, treating gene clusters as sets of functions. We finalized our annotation gene-set by retaining genes that had an Aragorn score above 106 bits or a tRNAscan-SE2 score above 49 bits, and reannotating sequences as described in the Results. All statistical analyses and sequence processing was done in R v.3.6 [43].

### Prediction of Divergent tRNA Class-Informative Features (CIFs) in Humans and Parasites

To compare CIFs between TriTrypDB genomes and humans and to have sufficient data to estimate CIFs, we defined eight phylogenetic clades for 40 of the 46 trypanosome genomes as shown in Table 1. These clades were based on a composite of phylogenetic results in the literature [44-47] and CIFs were subsequently analyzed independently by clade. We removed two incomplete genomes from analysis, *T. rangeli* SC58 and *T. cruzi* CLBrenner that had fewer than half the number of tRNA genes identified in any other genome. We filtered the gene annotation gene set, removing selenocysteine genes, pseudogenes, truncated genes, and genes of ambiguous function, leaving 3488 high-confidence functionally annotated gene records from TriTrypDB for analysis. To this set we added 431 high-confidence human tRNA gene records downloaded from GtRNADB [48] on May 15, 2019 (in the file “hg38-tRNAs.fas”). Two human selenocysteine tRNA genes were removed and the remainder combined with TriTrypDB data to yield a grand total of 3919 tRNA genes for CIF analysis. More statistics on the CIF estimate gene set by clade are shown in Table 1.

**Table 1.** Clades and Genomes analyzed, with statistics on CIF Estimation Gene Sets

| Clade | Genome Assemblies | Genes | Genes/<br>Genome | %G | %C | %T |
| --- | --- | --- | --- | --- | --- | --- |
| **Humans** | Hg38 | 431 | 431 | 31.9 | 25.1 | 22.4 |
| ***L. major***<br>**Clade (n = 8)** | 1. *L. major* Friedlin, 2. *L. major* LV39c5, 3. *L. major* SD75, 4. *L. tropica* L590, 5. *L. aethiopica* L147, 6. *L. gerbilli* LEM452, 7. *L. turanica* LEM423, 8. *L. arabica* LEM1108 | 664 | 83 | 31.3 | 25.4 | 22.7 |
| ***L. infantum***<br>**Clade (n = 3)** | 1. *L. donovani* BHU1220, 2. *L. donovani* BPK282A1, 3. *L. infantum* JPCM5 | 250 | 83.33 | 31.3 | 25.4 | 22.6 |
| ***L. Mexicana***<br>**Clade (n = 2)** | 1. *L. amazonensis* MHOMBR71973M2269, 2. *L. mexicana* MHOMGT2001U1103 | 148 | 74 | 31.3 | 25.4 | 22.6 |
| **Viannia** | 1. *L. braziliensis* MHOMBR75M2904 2. *L. braziliensis* MHOMBR75M2903 | 327 | 81.75 | 31.3 | 25.4 | 22.7 |
| **Subclade (n = 4)** | 3. *L. panamensis* MHOMPA94PSC1 4. *L. panamensis* MHOMCOL81L13 |  |  |  |  |  |
| ***L. enriettii***<br>**Clade (n = 3)** | 1. *Leishmania* sp MARLEM2494, 2. *Leishmania enriettii* LEM3045 | 160 | 53.33 | 31.2 | 25.3 | 22.7 |
| ***Leptospira/Crithidia***<br>**Clade (n = 3)** | 1. *Crithidia fasciculata* CfCl, 2. *Leptomonas seymouri* ATCC30220, 3. *Leptomonas pyrrhocoris* H10 | 300 | 100 | 31.2 | 25.4 | 22.7 |
| **American *Trypanosoma* (n = 11)** | 1. *T. grayi* ANR4, 2. *T. cruzi* CLBrennerEsmeraldo-like, 3. *T. cruzi* CLBrennerNon-Esmeraldo-like, 4. *T. cruzi* Dm28c, 5. *T. cruzi* Dm28c, 6. *T. cruzi* Esmeraldo 7. *T. cruzi* JRcl4, 8. *T. cruzi* marinkelleiB7, 9. *T. cruzi* SylvioX10-1, 10. *T. cruzi* SylvioX10-1-2012 11. *T. cruzi* Tulacl2 | 773 | 70.27 | 31.5 | 25.6 | 22.3 |
| **African *Trypanosoma* (n = 6)** | 1. *T. bruceigambiense* DAL972, 2. *T. brucei* Lister427, 3. *T. brucei* TREU927, 4. *T. evansi* STIB805, 5. *T. congolense* IL3000, 6. *T. vivax* Y486 | 324 | 54 | 31.8 | 25.8 | 22.3 |

We aligned the CIF annotation gene set of 3919 genes using COVEA v.2.4.4 [49] to the eukaryotic tRNA covariance model supplied with tRNAscan-SE v.1 [50]. The output alignment was manually edited in SEAVIEW [51] to correct the misalignment of 595 human and trypanosome tRNA genes (almost exclusively of type Leu and Ser) at Sprinzl coordinates 45 and 47. Sequences were processed with the FAST toolkit [52]. For each cluster independently, we computed function logos [25], Information Difference logos and Kullback-Leibler Divergence logos [27] with a newly written Python 3 program tSFM (tRNA Structure-Function Mapper) v0.9.14 available on github (https://github.com/tlawrence3/tsfm), which we describe briefly here, and more fully in a forthcoming publication. tSFM provides a command-line user interface for estimating function, ID, and KLD logos using our published methods. tSFM additionally calculates tRNA CIFs for features at structurally paired sites in addition to single sites, which are defined as (*n_i_* × *n_i_*) × *BP*, where *n_i_* ∈ {*A*, *C*, *G*, *U*, −} is the set of possible states at an alignment position and *BP* is the set of pairs of Sprinzl Coordinates pairs involved in basepairs. We ran tSFM with options “-e Miller -x 5” corresponding to computing entropies exactly for samples of size five sequences or fewer, and the Miller-Madow estimator otherwise.

Briefly, we computed the gain-of-information of a CIF in a particular functional class and trypanosome clade as its information difference in bits, with that clade as foreground and humans as background, multiplied by the normalized ratio of posterior-to-prior odds of the CIF in that functional class in trypanosomes and humans, corresponding to letter heights in ID logos, and measured in bits. We computed change-of-function of a CIF in a particular functional class and trypanosome clade as its Kullback-Liebler Divergence in bits, with that clade as foreground and humans as background, multiplied by the normalized ratio of posterior-to-prior odds of the CIF in that functional class, corresponding to letter heights in KLD logos and measured in bits. To avoid division by zero when calculating KLD, we added pseudocounts to either the background or the foreground posterior distributions when one or more of the 21 functional classes was not observed. When calculating the normalized ratio of posterior-to-prior odds for a specific functional class, we only added pseudocounts to the background posterior distribution. Furthermore, to avoid inaccuracies, we defined the KLD of feature to be zero when its frequency in the background is less than or equal to five.

We wrote a custom script in R 3.6 to visualize CIFs within each cluster for each functional class of tRNA in a structural context, and color the parasite CIFs according to whether those CIFs have gained information or changed function relative to human since divergence from their common ancestor. All data and scripts are provided as supplementary data.

### AaRS Cloning and Protein Purification

*Leishmania major* (*Lm*) AlaRS and *Lm* ThrRS-encoding genes were codon optimized, synthesized, and sub-cloned into pUC57 (GenScript). Engineered flanking NdeI and SmaI restriction sites were used to clone the aaRS genes into pTYB2, creating in-frame C-terminal intein fusions. The resulting expression vectors were transformed into the *E. coli* expression strain BL21 (DE3). The gene encoding *Tc* AlaRS was codon optimized, synthesized, and directly cloned into NdeI and XhoI cut sites in the pET21b expression vector (GenScript). The resulting plasmid expressed *Tc* AlaRS under T7 control and was in-frame with a C-terminal 6x-His tag. The pET21b-*Tc* AlaRS vector was transformed into the *E. coli* expression strain XJb (DE3) (Zymo Research).

Both *Lm* AlaRS and ThrRS were purified by growing cells to an OD600 of ∼0.5 and cooling on ice for 30 minutes. Protein induction was initiated by the addition of IPTG to a final concentration of 500 µM and cells continued to grow at 16°C for 16 hours. Cells were harvested by centrifugation and lysed by sonication in Buffer A (25 mM HEPES pH 7.2, 500 mM NaCl, 3 mM DTT) with cOmplete mini protease inhibitor (Sigma) added. Clarified lysate was added to a chitin resin column (NEB) and washed with Buffer A. The intein tag was cleaved by the incubation of Buffer B (25 mM HEPES pH 7.2, 100 mM NaCl, and 100 mM DTT) on the resin bed overnight at 4°C. Protein was dialyzed in two stages in Buffer C (25 mM HEPES pH 7.2, 30 mM NaCl, 6 mM BME, and 10% - 50% glycerol).

*Trypanasoma cruzi* (*Tc*) AlaRS-expressing cells were grown to an OD600 ∼0.3 and then cooled to 18°C and induced with 500 µM IPTG. Cells were grown for an additional 16 hours at 18°C before harvesting by centrifugation. Cell pellets were re-suspended in lysis buffer [Buffer I (500 mM Tris-HCl pH 8.0, 300 mM NaCl, and 10 mM imidazole) with cOmplete mini protease inhibitor (Sigma)], sonicated, clarified, and cell lysate passed over a TALON metal affinity column (Takara). After washing the column with Buffer I, protein was eluted with Buffer II (Buffer I with 250 mM imidazole). Protein was dialyzed in two stages to remove imidazole and to store the enzyme in 50% glycerol.

Human AlaRS was expressed in *E. coli* Rosetta (DE3) (Novagen) from pET21a which encodes the human AlaRS gene in-frame with a C-terminal 6x-His tag (expression plasmid provided by Karin Musier-Forsyth, Ohio State University). Cell were grown to an OD600 of ∼0.5 and cooled on ice for 30’ before inducing expression with 500 µM IPTG. Upon induction, cells grew for an additional 16 hours at 20°C before harvesting. Human AlaRS was purified as described above with the addition of 5 mM β-mercaptoethanol to both Buffer I and Buffer II. All enzyme concentrations were determined by active site titration [53,54] using [^14^C]-alanine (Perkin Elmer) and [^14^C]-threonine (American Radiochemicals).

### Preparation of *in vitro* Transcribed tRNA

*Lm* tRNA^Ala^ (chr11. trna1-AlaCGC), *Lm* tRNA^Thr^ (chr23. trna6-ThrTGT), and *Tc* tRNA^Ala^ (TctRNA-Ala.03) DNA sequences were cloned into EcoRI and XbaI restriction sites in pUC18 by slow cooling complementary synthetic DNA oligos and ligation as previously described [55]. PCR was used to amplify 50 µg DNA template from the pUC18-tRNA plasmids to be used for T7 runoff transcription. *In vitro* transcription was performed with 40 mM Tris-HCl pH 8, 2 mM spermidine, 22 mM MgCl_2_, 5 mM DTT, 50 µg/mL BSA, 4 mM NTPs, 20 mM 5’GMP, 20 U Protector RNase Inhibitor, 2 U pyrophosphatase, DNA template, and T7 RNAP at 42°C for 16 hours. Transcription products were purified on a Diethylaminoethyl (DEAE) Sephacel (GE Healthcare) column in 20 mM Tris-HCl pH 8.0, 5 mM MgCl_2_, and 250 mM NaCl. tRNA was eluted from the resin with 1 M NaCl. The RNA was precipitated overnight at −20°C in 1/10^th^ volume sodium acetate and 3x volume ethanol and re-suspended in RNase-free H_2_O.

### Marine Natural Product Library

The marine natural products screening library comprises 5,304 fractions from organic extracts of marine-derived Actinobacterial fermentations (1 litre culture, following our standard protocol [56]). Crude extracts were fractionated in to six sub-fractions on Seppak C_18_ cartridges using a stepwise elution profile (20, 40, 60, 80, 100% MeOH/ H_2_O, 100% EtOAc). The resulting fractions were solubilized in DMSO (1 mL per fraction), 4 μL aliquots diluted 1:5 in DMSO, and arrayed in 384 well format (17 x 384 well plates). Following primary screening against *L. donovani*, 120 active fractions were arrayed as serial dilutions (8 x 2-fold dilutions; 50 - 0.4 μM) in 96 well format for aaRS screening.

### Screen for Aminoacylation Inhibitors

Serial dilutions from the marine natural product (MNP) library were screened using the following protocol. Aminoacylation reactions were performed at room temperature using 10 mM DTT, 8 mM ATP, 5 µM tRNA, 60-80 µM [^14^C]-Ala or [^14^C]-Thr, 100-500 nM aaRS, and DMSO or MNP samples. After incubating the reaction for either 15 or 20 minutes, 1 µL of the reaction was spotted on 5% pre-soaked TCA 3 MM Whatman filter paper. The precipitated tRNA-bound filter paper was washed 3x with 5% TCA, washed once with ethanol, and dried. The dried filter paper was exposed overnight on a phosphor imager screen and imaged the following day. Qualitatively, the phosphor image screen was examined for a change in signal intensity relative to the DMSO control; a decrease in phosphor image intensity indicates partial or full inhibition of the reaction in the presence of the inhibitor. While active concentrations were unknown for each of the MNP mixes, the serial dilution helped prevent false-positive identification. All lead candidates from the preliminary screen were confirmed using similar reaction conditions; the reactions were monitored over a time course and placed at 37°C. Samples were quantified using a scintillation counter.

### Pyrophosphate Exchange

Amino acid activation was monitored using ATP/PP_i_ exchange as previously described [57]. Reactions were performed at 37°C in 100 mM HEPES pH 7.2, 30 mM KCl, 10 mM MgCl_2_, 2 mM NaF, 2 mM ATP, 2 mM [^32^P]-PP_i_ (Perkin Elmer), 90 µM alanine, 160 nM AlaRS, and DMSO or aaRS inhibitor. At increasing time points, aliquots of the reaction mixture were quenched in a charcoal solution containing 1% activated charcoal, 5.6% HClO_4_, and 75 mM PP_i_. Quenched reactions were vacuum filtered on to 3MM Whatman filter discs, washed three times with 5 mL of water and once with 5 mL of ethanol. After drying the filter discs, charcoal-bound radiolabeled ATP was quantified on a scintillation counter. Relative endpoint amino acid activation was determined by comparing the inhibitor-treated enzymes to their respective DMSO control samples.

## Results

### Custom Annotation of tRNA Genes and Gene Clusters in TriTrypDB Genomes

We obtained 4381 unified gene records from the output of two tRNA gene-finders, Aragorn and tRNAscan-SE v.2.0, to TriTrypDB v.41. Of these, 3597 were found by both gene-finders, 750 were found by Aragorn only, and 34 were found by tRNAscan-SE 2.0 only. We identified the same 76 genes as initiator tRNA genes, using either tRNAscan-SE 2.0’s profile-based predictions or our own edit-distance-based clustering approach, by finding the unique set of genes carrying conserved initiator tRNA features as described in [42]. There were 45 functionally ambiguous but high scoring genes, including 2 with identity unassigned by both gene-finders, 6 marked as pseudogenes or truncated by tRNAscan-SE 2.0, 4 containing sequence ambiguities, and 33 with conflicting structural and anticodon annotations, including 10 intron-containing genes predicted as tRNA^Tyr^ genes by tRNAscan-SE and tRNA^Asn^ genes by Aragorn, all from genomes in the American *Trypanosoma* clade. We annotated these as tRNA^Tyr^ genes, which are known to contain introns in that clade [58], as this helped complete the sets of functional types for 8 genomes in that clade (evidence not shown; one genome for *T. Cruzi* dm28c remained incomplete, missing a gene for tRNA^Phe^).

To further refine the final annotation gene set, we identified tRNA gene clusters in TriTrypDB genomes using a maximum intergenic distance criterion of 1000 bp on either strand. Doubling this distance criterion did not substantially increase cluster number or size. After filtering 4381 gene records by their gene-finder scores as described below, 3616 high-confidence gene records remained, of which 77% occur in clusters of size two or greater (Fig. 1). The largest tRNA gene clusters were of size ten, accounting for 9% of total genes. After clustering tRNA gene clusters with similar gene content and strandedness, we could identify putatively orthologous tRNA gene clusters conserved within each of the *Leishmania* and *Trypanosoma* genera, and sometimes both genera, with substantial evidence of evolution in gene organization through duplication, divergence, inversion and other changes (Supplementary Tables 1-3).

**Fig 1.**
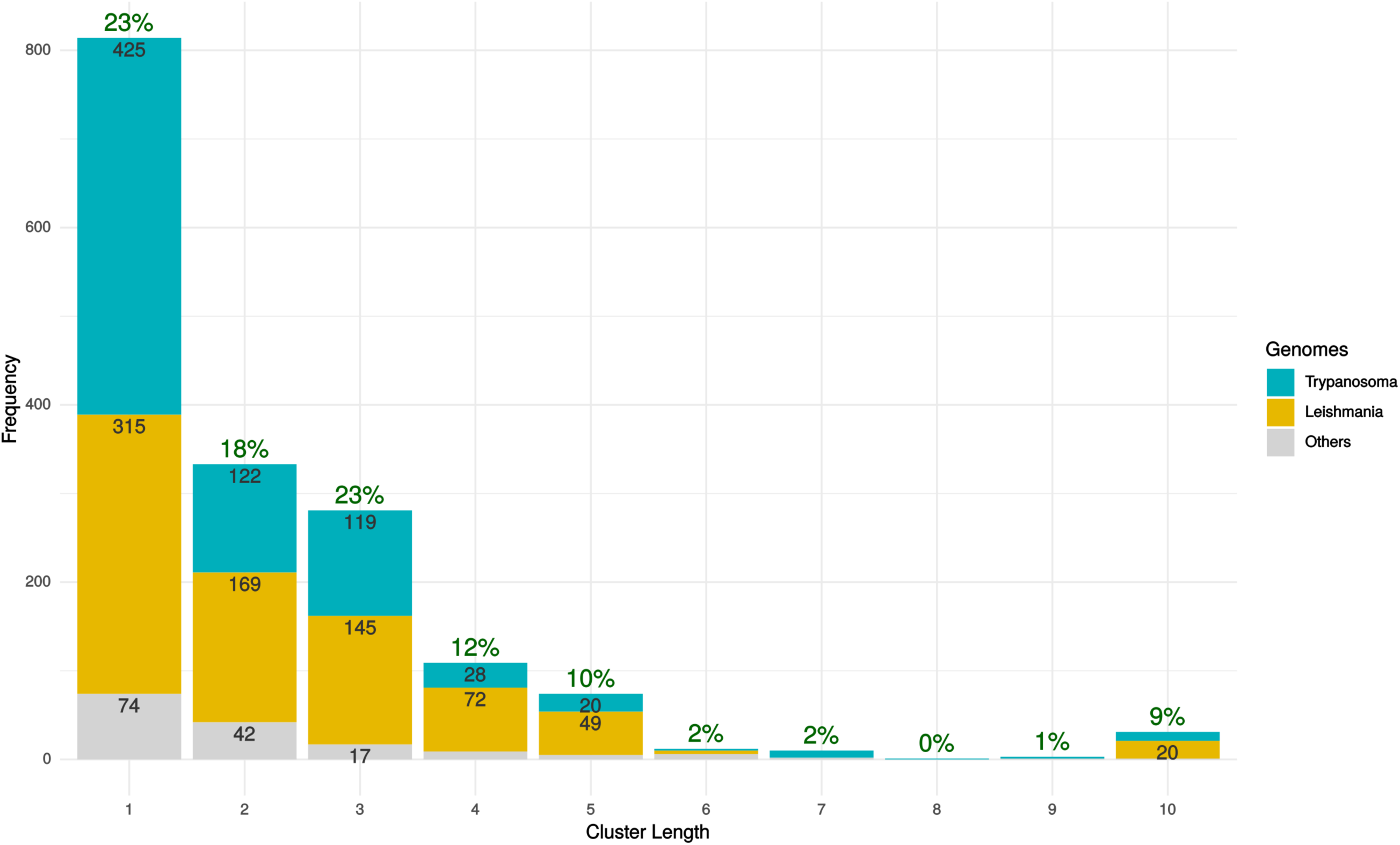
tRNA gene cluster size distribution for *Trypanosoma*, *Leishmania*, and other TryTryp genomes. Green labels at tops of stacks show percentages of total tRNA genes in clusters of a given length. Numbers within each bar show frequencies of gene clusters of that length.

Using the similarities of tRNA gene clusters across genomes, we found putative homologs for some of the 45 functionally ambiguous but high-scoring genes marked as pseudogenes or truncated by tRNAscan-SE 2.0 as well as some genes detected only by Aragorn. With these results in mind, we plotted the densities of gene-finder scores according to whether they were found by both or only one gene-finder, showing clear evidence of a small fraction of Aragorn-only genes with high scores, making up about 1% of our total finalized gene set (Fig. 2 and Supplementary Table 4).

**Fig 2.**
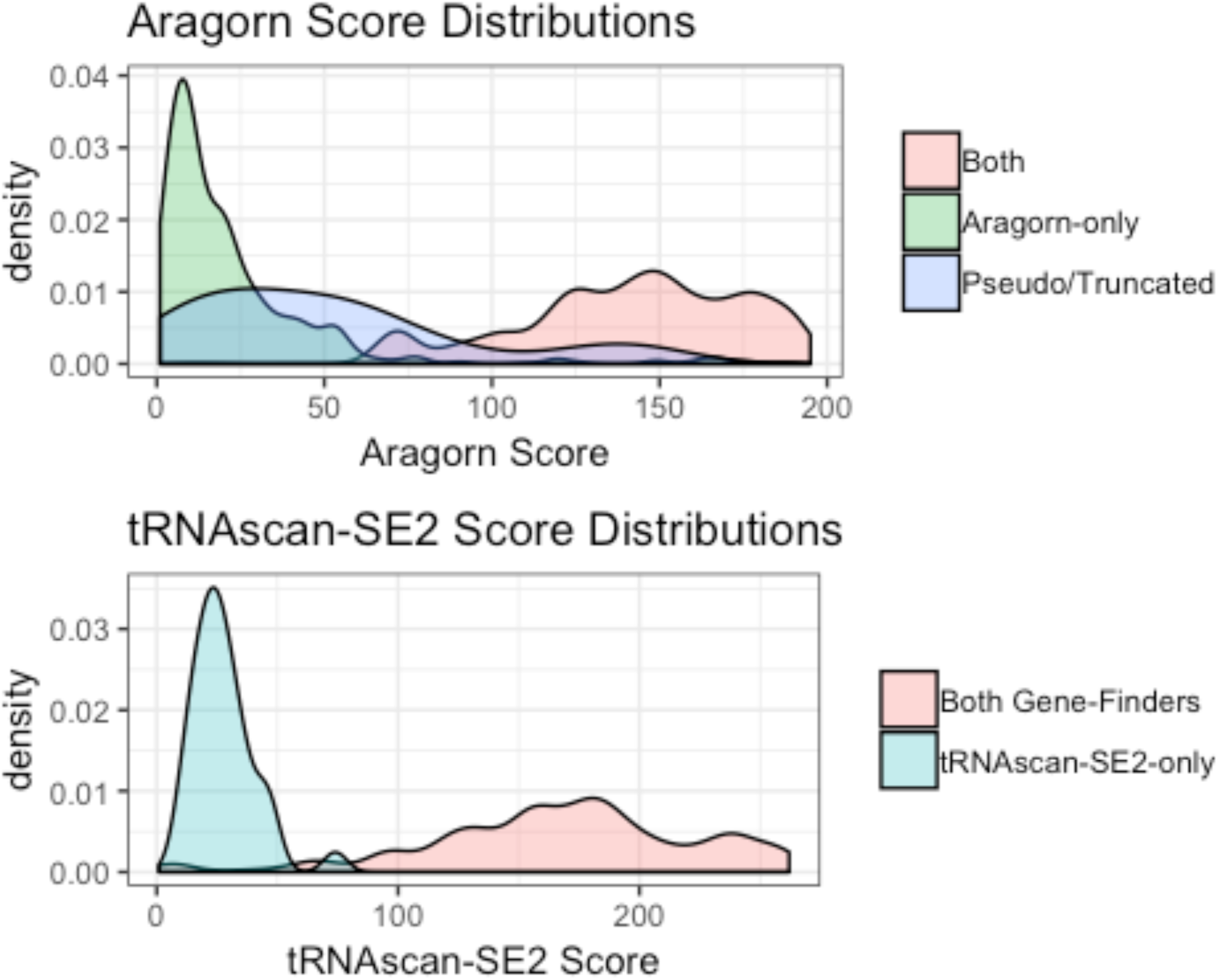
Density Plots of Gene-Finder Scores According to Source of Detection.

Based on this evidence, we retained 3616 genes that had an Aragorn bit-score of at least 107 or a tRNAscan-SE 2.0 bit-score of at least 50, including 35 genes found by Aragorn only and 1 gene found by tRNAscan-SE 2.0 only. These score cutoffs separated Aragorn-only genes within conserved gene clusters from singletons, which had lower scores (evidence not shown). The median number of genes per genome was 83. Supplementary Table 4 shows statistics on the final gene set broken down by genome. Two incomplete genome assemblies, belonging to *T. cruzi* Brener and *T. rangeli* SC58, were annotated with only six or 18 genes, respectively, and were excluded from further analysis.

### Divergent Class-Informative Features between Humans and Try-TrypDB Genomes

We developed a bioinformatic workflow that combines information from tRNA function logos estimated from a parasite clade and Information Difference (ID) logos [25] and Kullback-Leibler Divergence (KLD) logos between the parasite clade and humans [27]. The workflow quantitates tRNA features that are functionally informative in the parasite clade and have either gained or retained functional information or altered functional association since divergence of the parasite clade and humans from their common ancestor. We found many examples of highly informative trypanosome CIFs that have been gained, retained or changed function since divergence from their common ancestor with humans, and most of these divergent CIFs have been strongly conserved in trypanosomes over 231–283 million years of evolutionary divergence between *Leishmania* and *Trypanosoma* [59], for example among alanine tRNAs (Fig. 3) and threonine tRNAs (Fig. 4). Structural bubble-plot visualizations at single-site resolution of these CIF divergence measures are provided for all functional classes in supplementary materials, showing that some classes have diverged much more than others. Even though they are calculated at single-site resolution, CIF divergences are correlated across structurally paired sites. Inspection of function logos across taxa confirms the conservation of parasite-specific CIFs and reveals A and U features underlying the signals shown in Figs 3 and 4, including some sharing of divergent features between tRNA^Ala^ and tRNA^Thr^ functional classes, for example at Sprinzl coordinate 39 (Fig. 5). Inspection of the data shows that a U:A informative base-pair diverged to an adjacent paired site during tRNA^Thr^ gene divergence between trypanosomes and humans, from 31:39 to 30:40 (Fig. 5). This can be seen more easily by inspecting function logos for paired sites directly. Figures 6-8 show base-pair function logos for Humans, the *L. major* clade and the American *Trypanosoma* respectively, showing that both Class Informative Base-Pairs (CIBPs) and Class-Informative Mis-Pairs (CIMPs) can be relatively conserved, and that recurring hot-spots of CIF evolution appear in the data, yielding insight to mechanisms of CIF evolution. Full function logo results for all clades are provided in supplementary data.

**Fig. 3.**
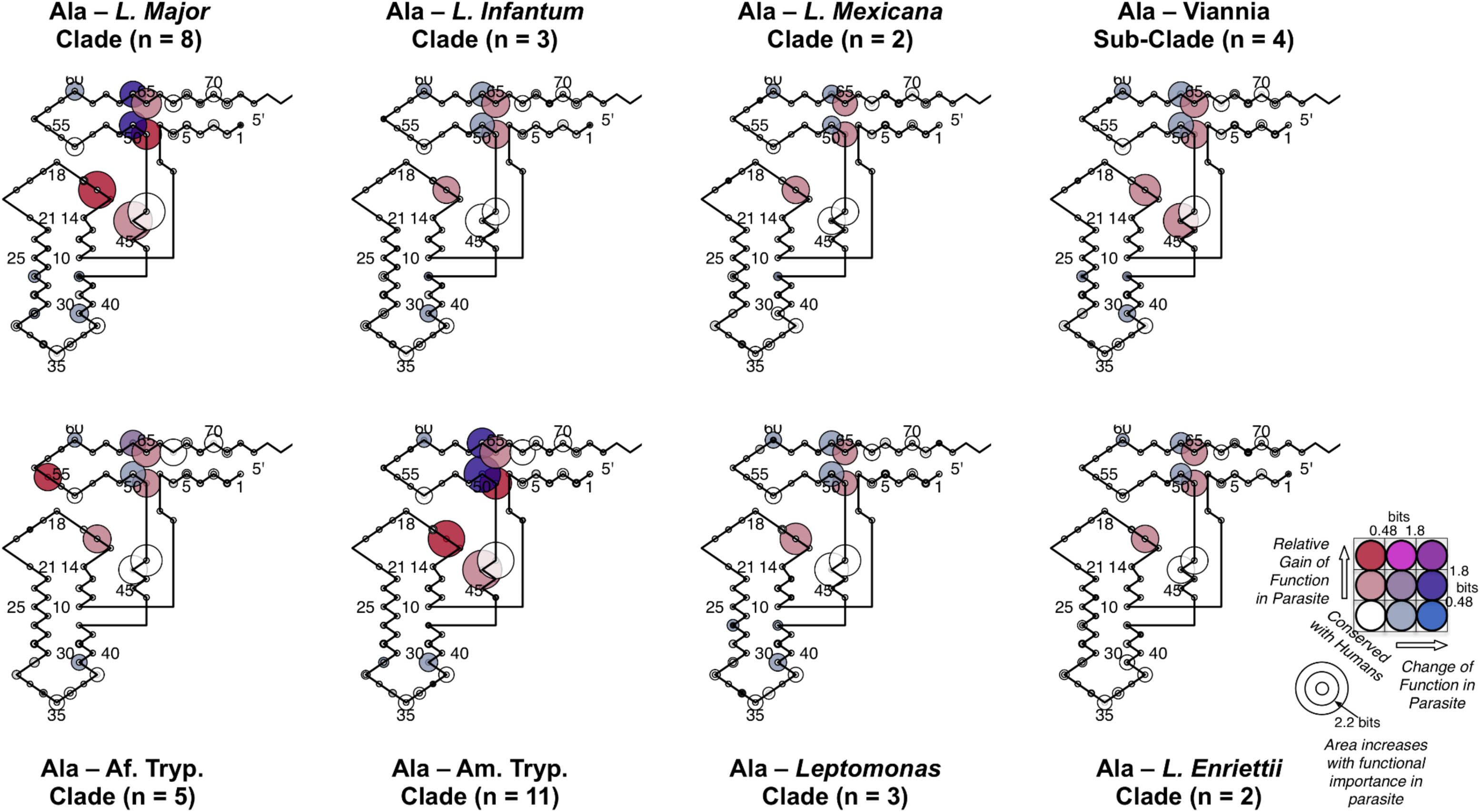
Conserved divergence of parasite tRNA^Ala^ CIFs across eight phylogenetic clades of *Leishmania* and *Trypanosoma*. Evolutionary divergence of trypanosomes relative to *Leishmania major* increases clockwise from *Leishmania major*.

**Fig. 4.**
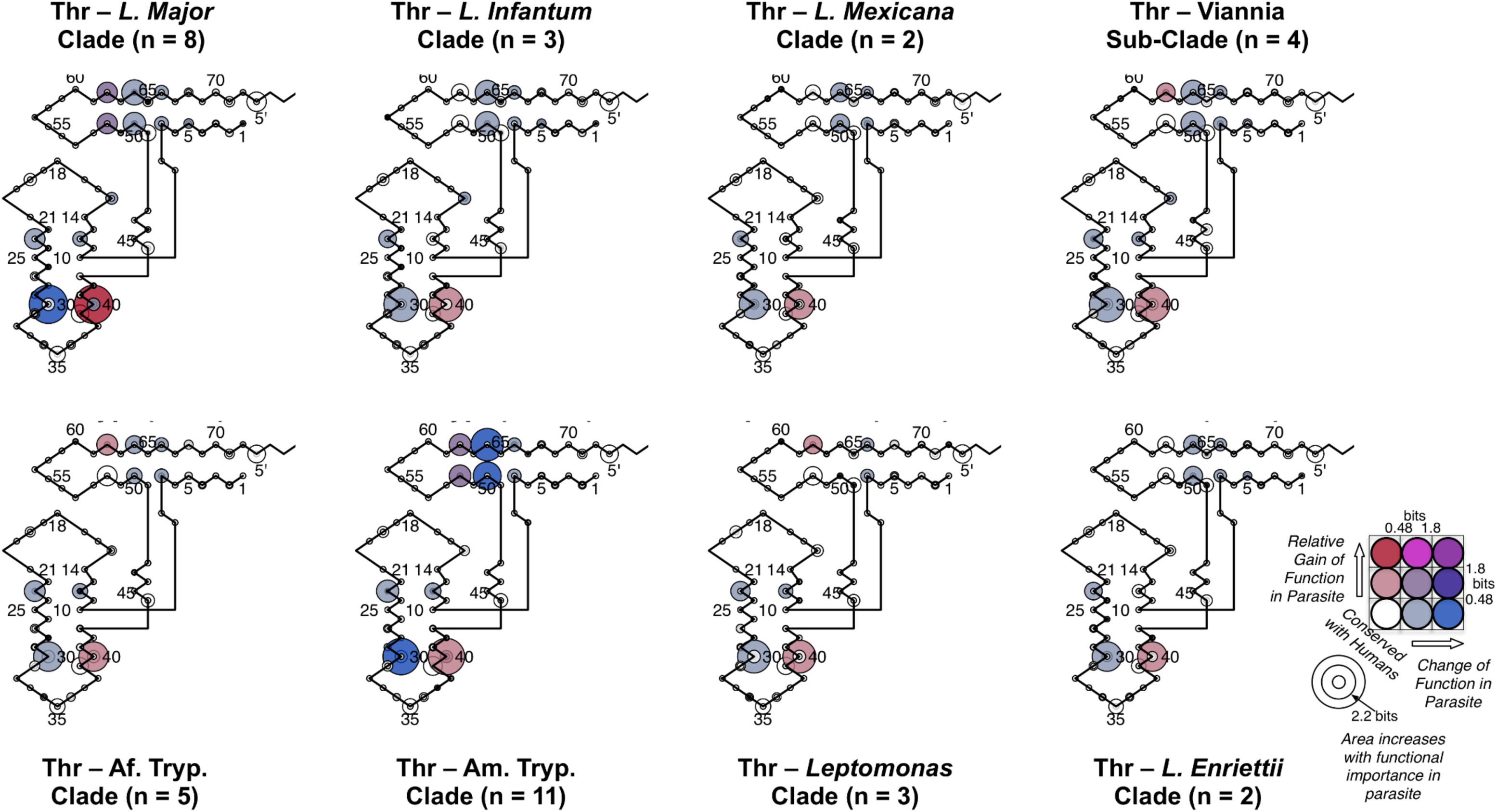
Conserved divergence of parasite tRNA^Thr^ CIFs across eight phylogenetic clades of *Leishmania* and *Trypanosoma*. Evolutionary divergence of trypanosomes relative to *Leishmania major* increases clockwise from *Leishmania major*.

**Fig. 5.**
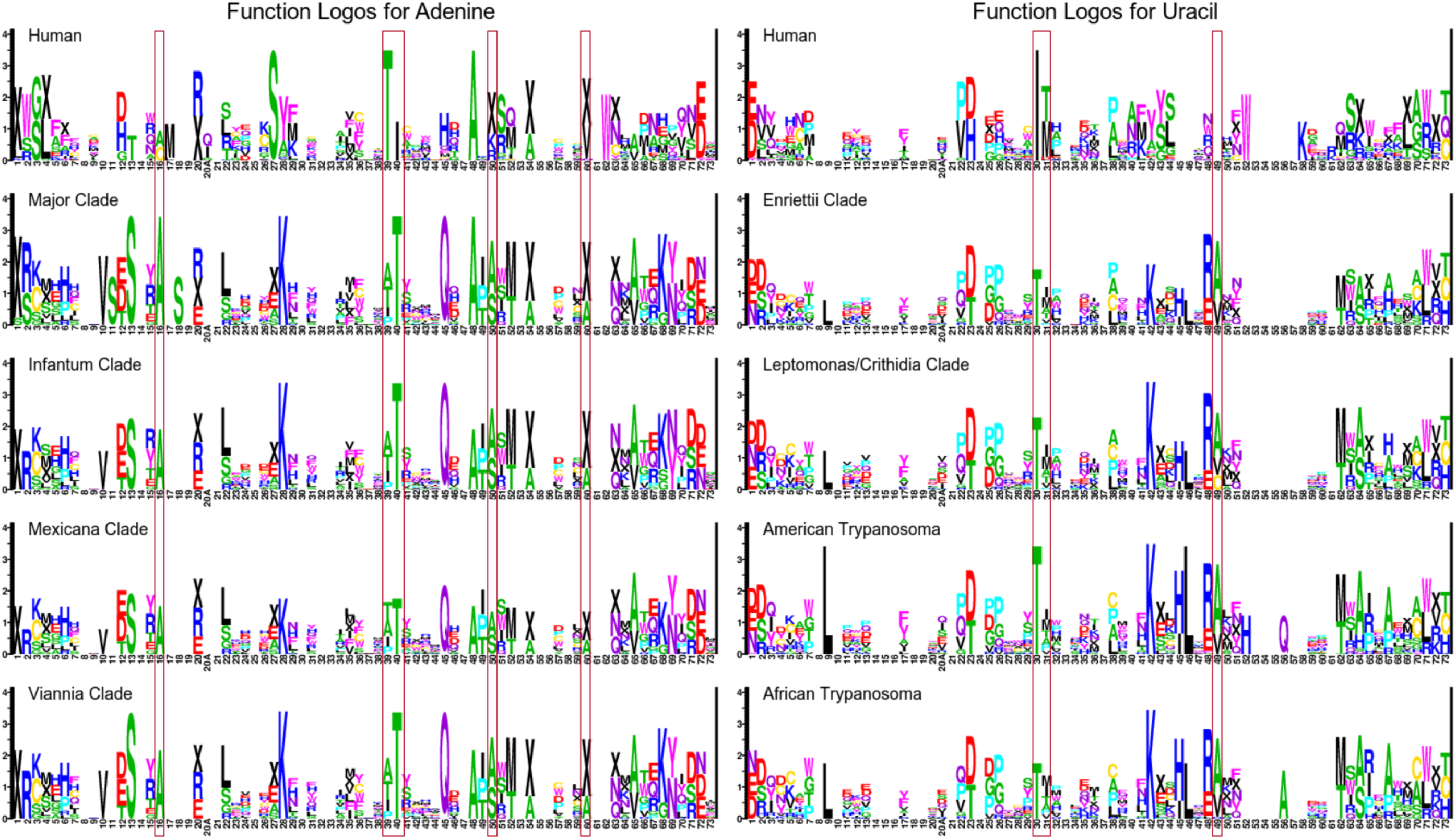
Adenine and Uracil function logos for humans and eight phylogenetic clades of *Leishmania* and *Trypanosoma*. A selection of sites with CIF divergence between humans and trypanosome clades for tRNA^Ala^ and tRNA^Thr^ are indicated.

**Fig. 6.**
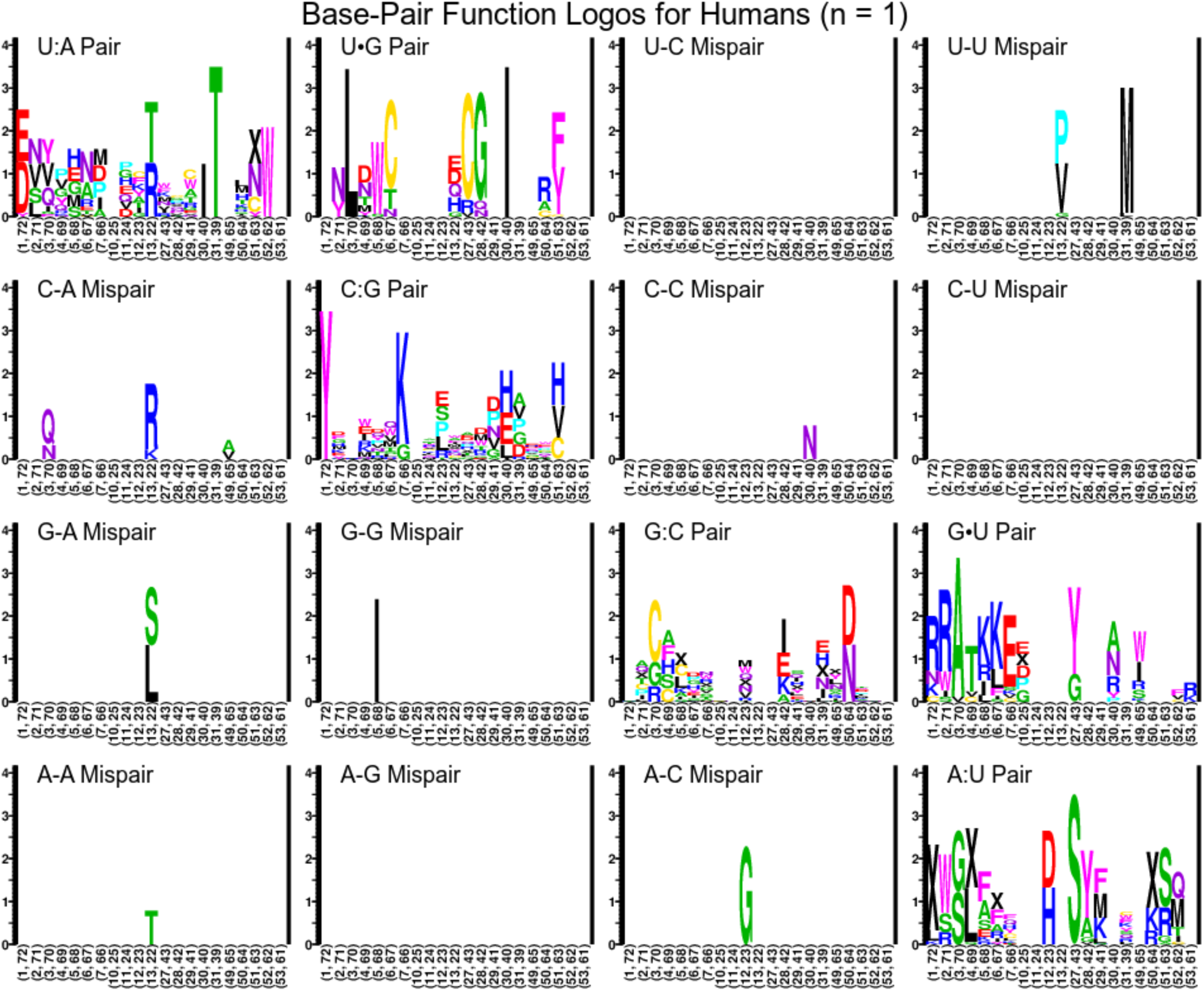
Function logos for tRNA Class-Informative Base-Pairs and Class-Informative Mis-Pairs in humans.

**Fig. 7.**
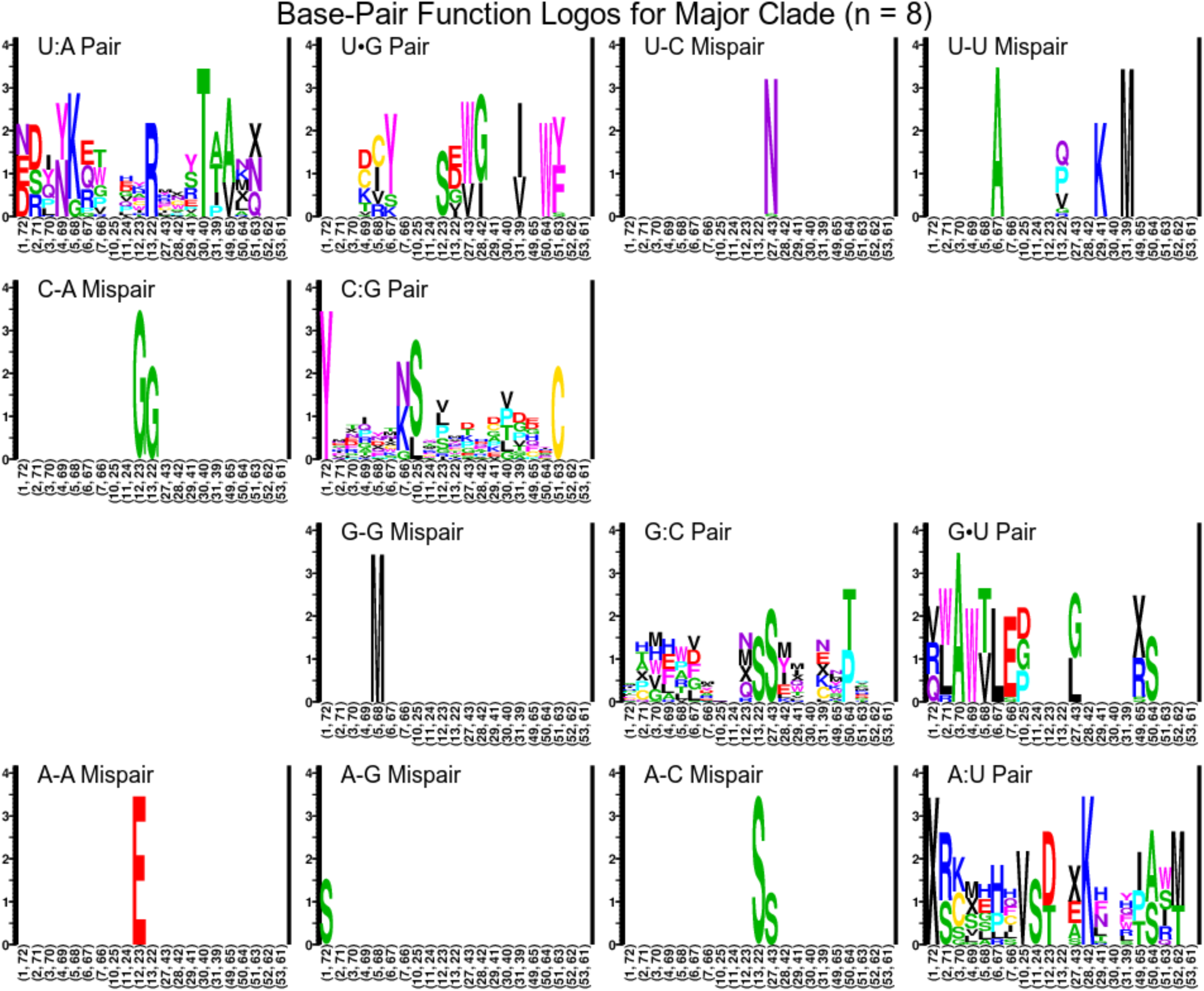
Function logos for tRNA Class-Informative Base-Pairs and Class-Informative Mis-Pairs in the *L. major* clade.

**Fig. 8.**
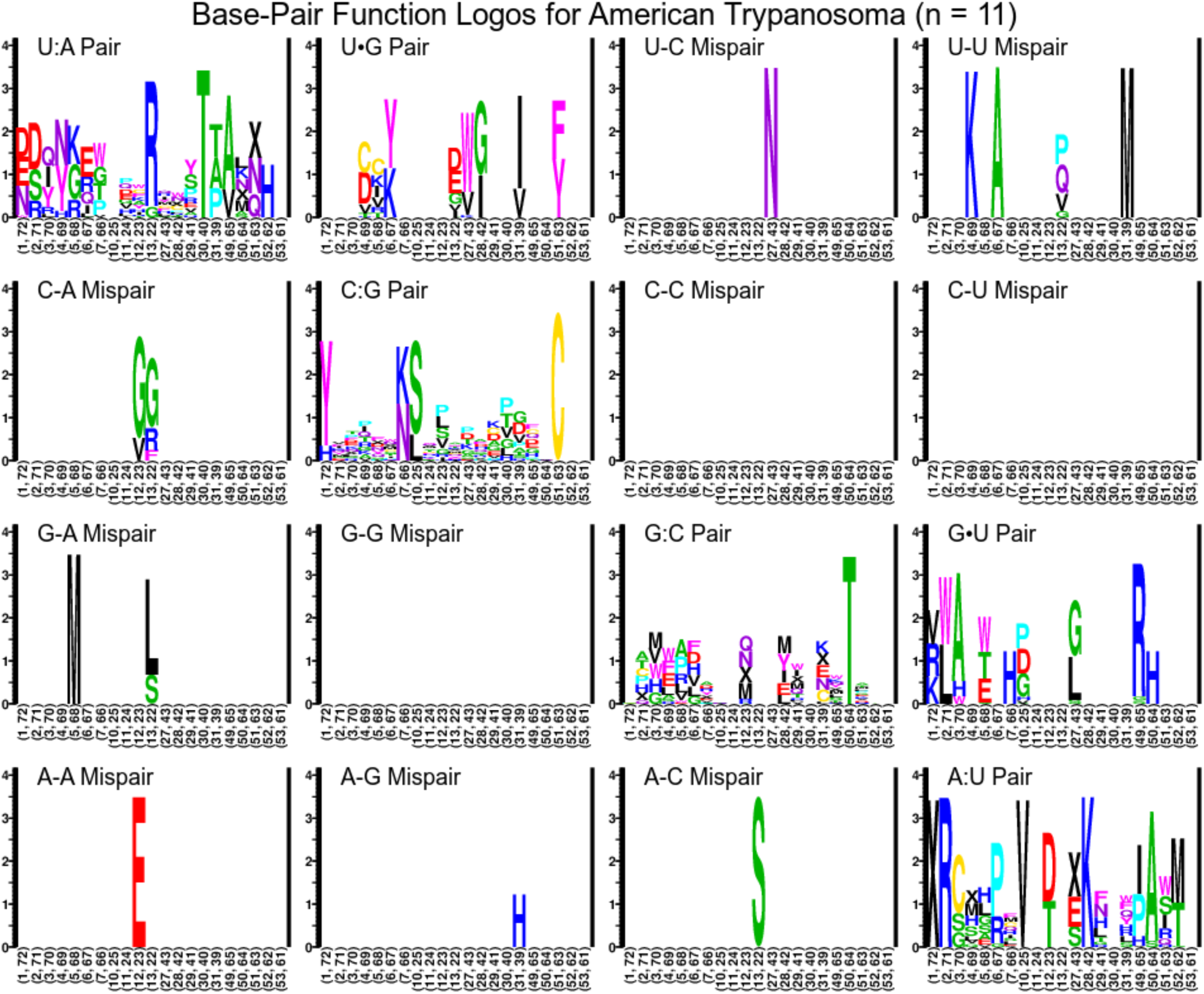
Function logos for tRNA Class-Informative Base-Pairs and Class-Informative Mis-Pairs in the American *Trypanosoma* clade.

### AaRS Screen Identified *Leishmania major* AlaRS Inhibitors

Using the pre-validated MNP library, we developed a medium-throughput aminoacylation screen to identify aaRS inhibitors *in vitro* (Fig. 9A and 9B). From the one hundred and twenty complex inhibitory mixes tested in the MNP library, we qualitatively identified four potential *Lm* AlaRS inhibitors as determined by a decrease in the overall tRNA-aminoacylation signal. These four candidates were then re-screened using time-dependent quantitative approaches and we concluded that three of the four mixes, 1881C, 2059D, and 2096B were altering aminoacylation, with inhibitory activities ranging between 80% and 99% (Fig. 9C).

**Fig. 9.**
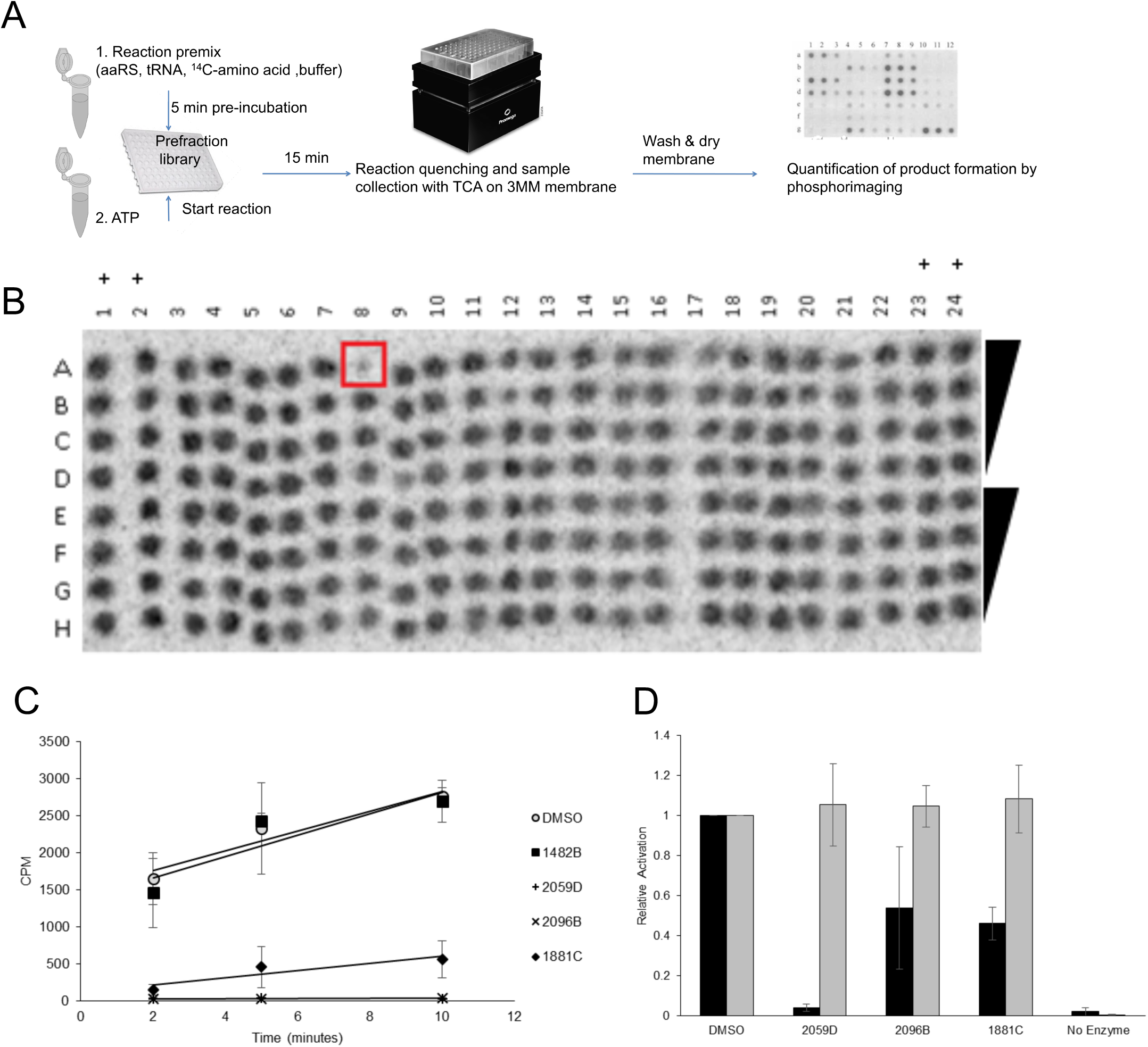
Identification of *Leishmania major* AlaRS inhibitors. A) Workflow to identify aminoacylation inhibitors (details described in Methods). B) Representative image of the MNP chemical screen. The spot boxed in red is an example of a predicted inhibitor depicted by the decrease in signal intensity. DMSO positive control (+). C) Three of the four identified inhibitors prevented the accumulation of Ala-tRNA^Ala^ formation, 1428B was a false-positive result from our preliminary screen. D) The three identified inhibitors perturbed *L. major* AlaRS activation (black) but had no effect on human AlaRS (gray). The relative amino acid activation is plotted relative to the DMSO control. Error bars indicate the standard deviation of three replicates.

Since the aminoacylation screen discerns total net changes to the aaRS activity, we attempted to identify which step of the aaRS catalyzed reaction is being affected by the MNPs. To observe any tRNA-independent effects on aaRS function, we used pyrophosphate exchange to monitor ATP-dependent amino acid activation. From this experiment, we were able to conclude that our inhibitors were perturbing amino acid activation, with lead compounds ranging in inhibitory activity between 45% and 95%. The differences in MNP activity between amino acid activation and tRNA-dependent aminoacylation highlight the multiple aaRS activities that can be targeted in our network predictions. (Fig. 9D). To validate the predictive tool for identifying anti-trypanosomal drugs, we counter-screened the newly identified *Lm.* AlaRS inhibitors against the human AlaRS enzyme. Treatment of the human AlaRS enzyme with the MNP inhibitors had no effect on amino acid activation (Fig. 9D). Combined with our original screening data, these results show the utility of our computational and biochemical workflow to identify new novel therapeutics that have minimal cross-reactivity with the human homolog of the parasite drug target.

### Natural Product Library Inhibitors of *Leishmania major* ThrRS

As our network predictions identified CIF divergence among many functional classes of tRNAs (as shown in the Supplementary materials), we also wanted to determine if our MNP library would find inhibitors against non-AlaRS aaRS. The most concentrated mixes from our MNP library were re-screened against *Lm* Threonyl-tRNA synthetase (ThrRS) and tRNA^Thr^ aminoacylation (Fig. 10A). The preliminary screen led to the identification of eight extracts with inhibitory activity. Those compounds were re-analyzed using quantitative aminoacylation reactions and the results show that that two of the candidates did not inhibit aminoacylation, two inhibited the reaction at ∼50%, and four had greater than 75% inhibition (Fig. 10B). In addition, two of the four most active inhibitors (2059D and 2096B) also had activity against *Lm* AlaRS (Fig. 10C). The cross-reactivity of these inhibitors may be a consequence of the extensively conserved aaRS architecture found between AlaRS and ThrRS [60,61].

**Fig. 10.**
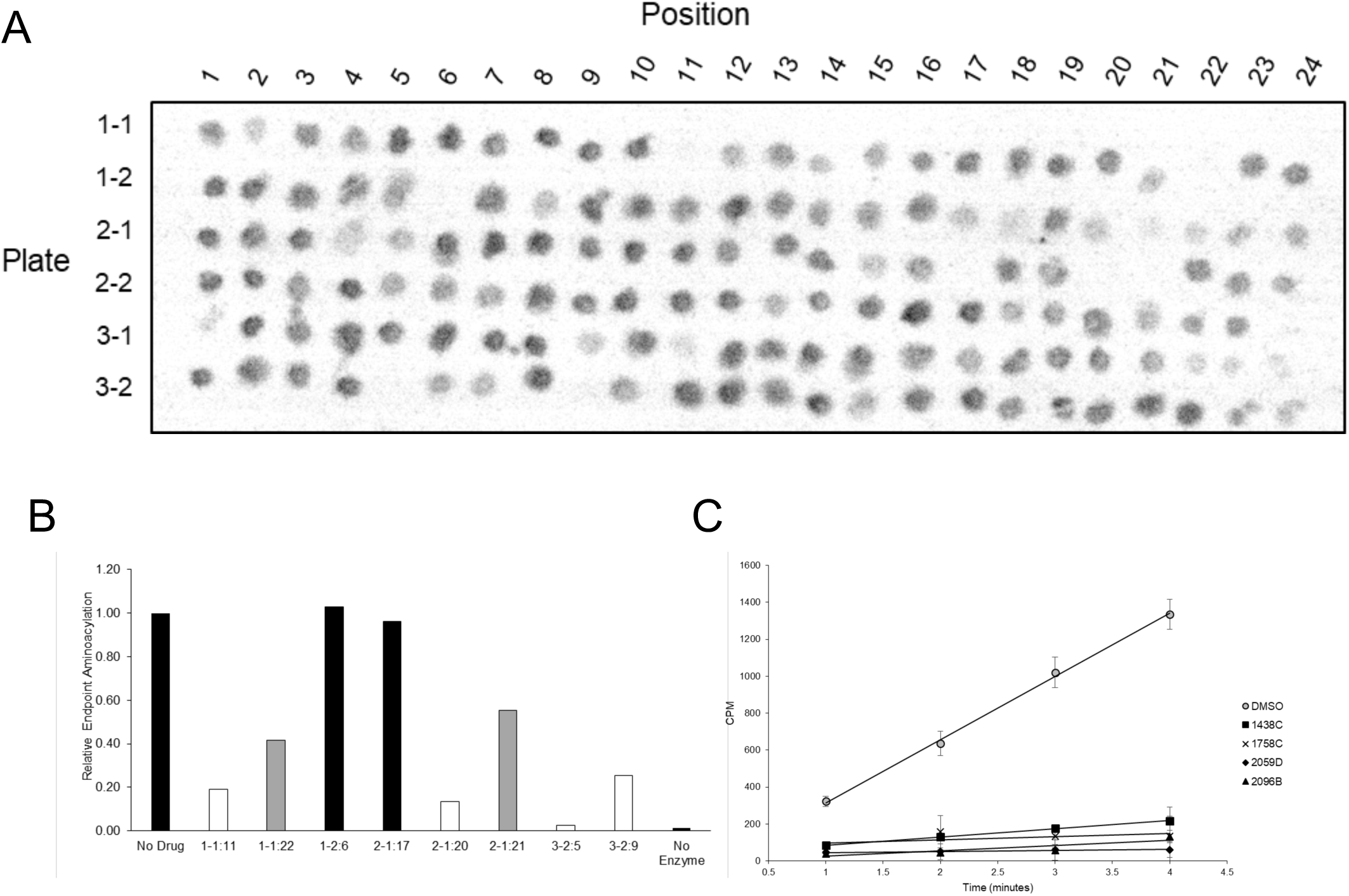
Identification of *Leishmania major* ThrRS inhibitors. A) The MNP library was re-screened at the highest concentrations to qualitatively identify *Lm* ThrRS aminoacylation inhibitors. Plate IDs reference the position within the original library and not library IDs. B) Eight inhibitors were qualitatively identified from the preliminary screen. Two of the candidates did not inhibit aminoacylation (black), two inhibited at ∼50% activity (gray), and four inhibited at greater than 25% (white).C) All four active inhibitors continued to perturb aminoacylation over a time course experiment. Error bars indicate the standard deviation of three replicates.

### Predictive Network Interactions Identified Broad-Spectrum Anti-trypanosomal Targets

The tRNA-aaRS network analyses suggested that parasite-specific tRNA^Ala^ identity elements were highly conserved between the *Leishmania* and *Trypanosoma* genera (Figs. 3,4,5,11A). To test this hypothesis, we purified *Tc* AlaRS and screened our three active *Lm* AlaRS inhibitors in an aminoacylation inhibition assay using *Tc* AlaRS and tRNA^Ala^. Supporting our network prediction, all three *Lm* inhibitors also had activity against the *Tc* enzyme, with activities ranging between 40% and 95% total inhibition (Fig. 11B). While these activities were slightly reduced compared to their effect of the *Lm* AlaRS enzyme (Fig. 9C), these results highlight the additional potential utility of our computational methodologies as a means of identifying broad-spectrum antimicrobials for closely-related clades.

**Fig. 11.**
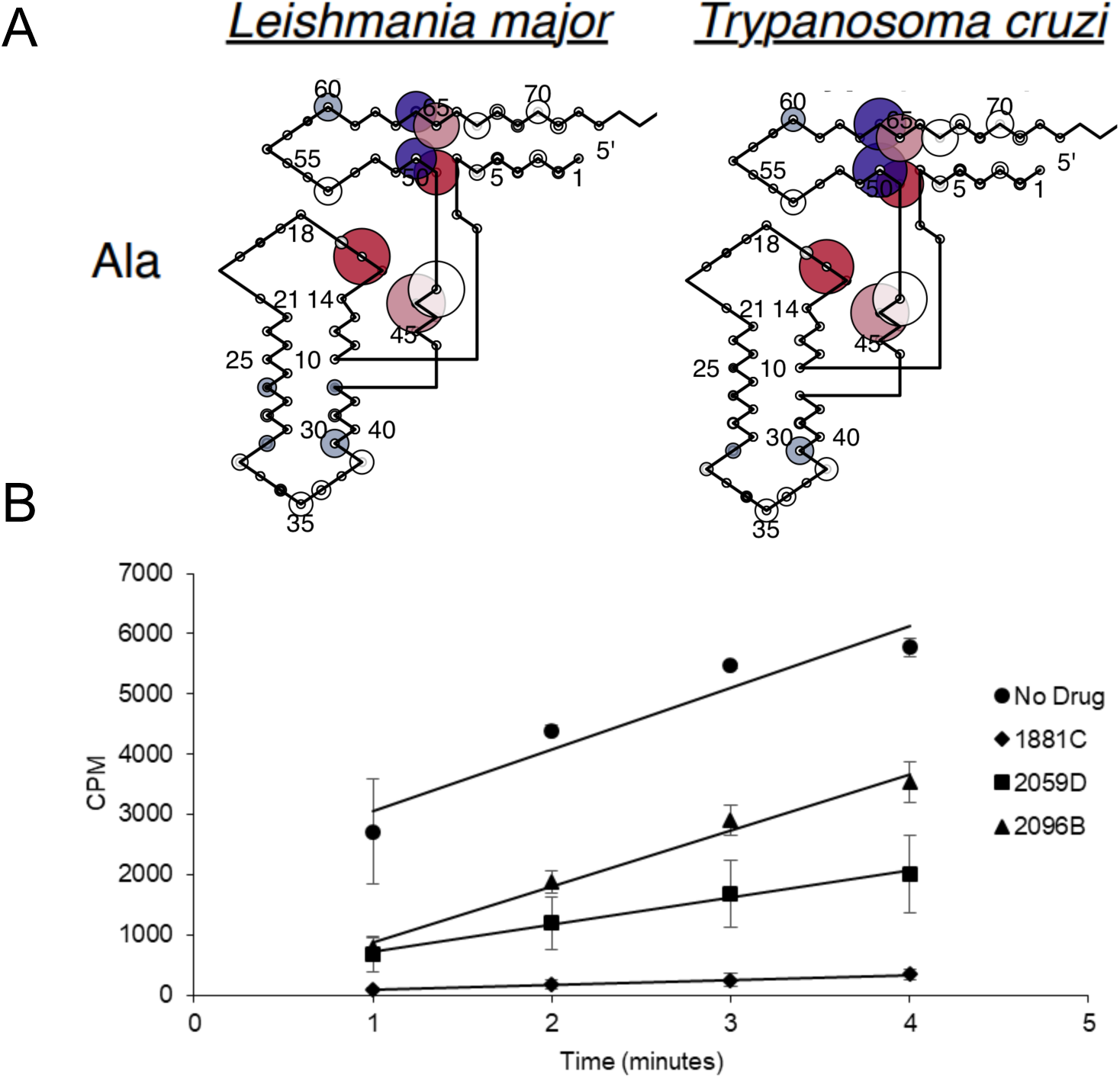
*Leishmania major* and *Trypanosoma cruzi* AlaRS have conserved tRNA identity elements. A) CIF Divergence Models for tRNA^Ala^ in Leishmania *major* and *Trypanosoma cruzi* B) The three identified *Lm* AlaRS inhibitors also have activity against the *Tc* AlaRS enzyme. Error bars indicate the standard deviation of three replicates.

### Separation of Active Components from Natural Products Extracts

From the initial set of 120 extracts with activity against *L. donovani* parasites, four extracts showed corresponding activity in the initial aaRS assay. Of these, three (1881C, 2059D, and 2096B) were prioritized for chemical follow up, based on potent, dose dependent biological activity. Initially, each sample was separated into 10 sub-fractions using HPLC (Phenomenex Synergy C18, 5μ, 4.6 x 250 mm). Screening of these fractions identified one fraction (2096B F10) with potent activity. To generate additional material, the producing organism was cultured on large scale (1 L, GNZ medium with 20 g XAD-7 resin), filtered, and the resin/cell slurry extracted with organic solvents (2:1 CH_2_Cl_2_/ MeOH, 400 mL). The crude extract was fractionated using an automated Combiflash chromatography system (C_18_ cartridge; 20, 40, 60, 80, 100% MeOH/ H_2_O, 100% EtOAc) and the resulting fractions subjected to biological screening (Text S1). Two fractions (C and D) showed strong activity and were subjected to subsequent separation to give 10 sub-fractions each (Fig S38-41). Of these, fraction 2096D F10 showed the strongest reproducible activity (Fig. S42). However, subsequent fractionation steps yielded sub-fractions with very low quantities of material. Review of these sub-fractions by UPLC-ESI-qTOF mass spectrometry did not identify any individual mass signatures consistent with a candidate bioactive molecule. Similarly, lack of material precluded the identification of diagnostic signals in the NMR spectra for these subfractions. Provisional information from these analyses, including NMR and MS signatures from earlier fractions and the non-polar nature of the active fractions, suggest that the active component is likely a bioactive lipid, although the precise nature of the structure of this metabolite remains unknown.

## Discussion

### Systems-biology driven identification of trypanosome-specific drug targets

tRNA CIFs apply an information criterion, rather than a conservation criterion, to bioinformatically predict tRNA identity elements. By applying this information criterion independently over different parasite clades, we found that tRNA CIFs were highly conserved over 250 million years of trypanosome evolution. A biological interpretation of this result is that the information contained in tRNA CIFs is functional in the specification of the identities of substrates and non-substrates to aaRSs.

Maintaining efficient and accurate translation is predicated on catalytically productive interactions between aaRSs and free tRNAs in the cell. While the major identity elements for a given aaRS-tRNA pair are generally conserved, here we have identified divergent features within tRNAs that apparently contribute to divergent RNA-protein interaction in trypanosomes. Much of the focus in this work was on the phylogenetic divergence of identity elements among alanine tRNAs. This class of tRNAs strongly support the utility of our computational analyses as the tRNA^Ala^ identity elements have been one of the most well characterized to date [13,14]. Interestingly, it was recently shown that the conserved G3:U70 base pair is recognized by AlaRS using three distinct mechanisms across all domains of life [15]. This observation highlights that even highly conserved identity elements may be recognized and discriminated against by distinct biophysical aaRS interactions, which may therefore be stronger potentially specific drug targets than previously anticipated.

### Inhibition of aminoacyl-tRNA synthetases

A goal of this work was to identify divergent tRNA identity elements in trypanosome parasites. We predicted that parasite-specific tRNA-aaRS interactions would be identified, sufficiently divergent from homologous human machinery to be strong candidates for drug discovery. Interestingly, our network divergence analysis led to the discovery of tRNA-independent, amino acid activation inhibitors that were specific to trypanosomes. We interpret this as consistent with our goal, because tRNAs and aaRSs must coevolve to accommodate changes to structure and mechanism that evolve on either side of their interactions. Presumably, divergence in the structural mechanism of amino acid activation in trypanosome AlaRSs has also changed how they interact with their tRNA substrates. By integrating information from many tRNA functional classes, we gain leverage to interpret divergence in structure and function of the much more structurally complex ensemble of aaRSs as a system. Our tRNA-based network approach identifies potential aaRS targets that may not have been initially predicted when analyzing aaRS functional sequences in isolation.

### Combination chemotherapeutic inhibition of multiple aminoacyl-tRNA synthetases may be relatively resistance-proof

Two of the compounds we described were effective inhibitors of both AlaRS and ThrRS in parasites. Although monotherapeutic inhibitors of aaRSs are highly effective [62], combination therapies involving multiple aaRSs have not been studied. Because aminoacylation pathways are integrated in parallel into protein synthesis [63], their multiple inhibition is not expected to be synergistic, but rather “Bateson-Gaddum non-interaction masked” [64]. Unlike some synergistic combination therapies that can facilitate the evolution of drug-resistance [65,66], aaRS inhibitor combination therapies should be comparatively resistance-proof [32]. Further work is needed to realize this potential benefit.

## Conclusion

Trypanosome parasites pose a significant health risk worldwide. While current therapies exist, they are often also accompanied by off-target cytotoxicity and can lead to the rise of antimicrobial resistance. Here we have demonstrated that targeting tRNA-synthetase interactions have been an underexplored avenue for drug discovery. Using a combination of predictive computational tRNA network analyses and biochemical validation, we showed that aminoacyl-tRNA synthetases are a promising target for broad-spectrum anti-trypanosomal discovery with no significant consequence to the human counterpart target.

## Acknowledgements

The authors would like to thank Dr. Tammy Bullwinkle for assistance in generating expression constructs for this work. This work was supported by the National Institutes of Health grant AI127582 to MI, DA and RL.

## Supporting information

**S1-S21 Figures** Bubbleplots of CIF divergence from humans in eight clades of trypanosomes by tRNA functional class.

**S22-S29 Figures** Single-site function logos by type of feature for humans and trypanosomes.

**S30-S37 Figures** Paired-site function logos for humans and each trypanosome clade.

**S38 Figure** UPLC-qTOF base peak chromatogram of **(A)** RL12-182-HVF-D Sep-Pak Fraction C Subfraction 10-9 and **(B)** extracted ion chromatogram of peak 316.28 *m/z* in active region of trace.

**S39 Figure** UPLC-qTOF mass spectrum for peak at 316.28 *m/z*.

**S40 Figure** Schematic diagram of the large-scale fermentation and extraction of RL12-182-HVF-D.

**S41 Figure** HPLC-UV trace of the final preparative isolation step for the RL12-182-HVF-D active compound. Red box denotes active peak.

**S42 Figure** Marine natural product extract 2096D F10 inhibits *Leishmania major* AlaRS aminoacylation.

**S1 Table.** Putatively orthologous tRNA gene clusters of length at least 3 with frequencies in at least 2 *Leishmania* genomes. Clusters with the same group ID are considered similar.

**S2 Table.** Putatively orthologous tRNA gene clusters of length at least 3 with frequencies in at least 2 *Trypanosoma* genomes. Clusters with the same group ID are considered similar.

**S3 Table.** Putatively orthologous tRNA gene clusters of length at least 3 conserved across Leishmania (L) and Trypanosoma clade (T). Clusters ASD/DSA, LXP, and QLI are the only clusters conserved in both clades. This table shows groups of similar clusters within both clades.

**S4 Table.** Statistics on Final Annotation Gene Set.

**S1 Code and Data.** Code and data to reproduce computational results.

**S1 Text.** Supplementary methods for bacterial fermentation, natural product extraction, and active compound identification using HPLC-UV-MS and UPLC-ESI-qTOF-MS.

---

**Supplementary Figure 1.**
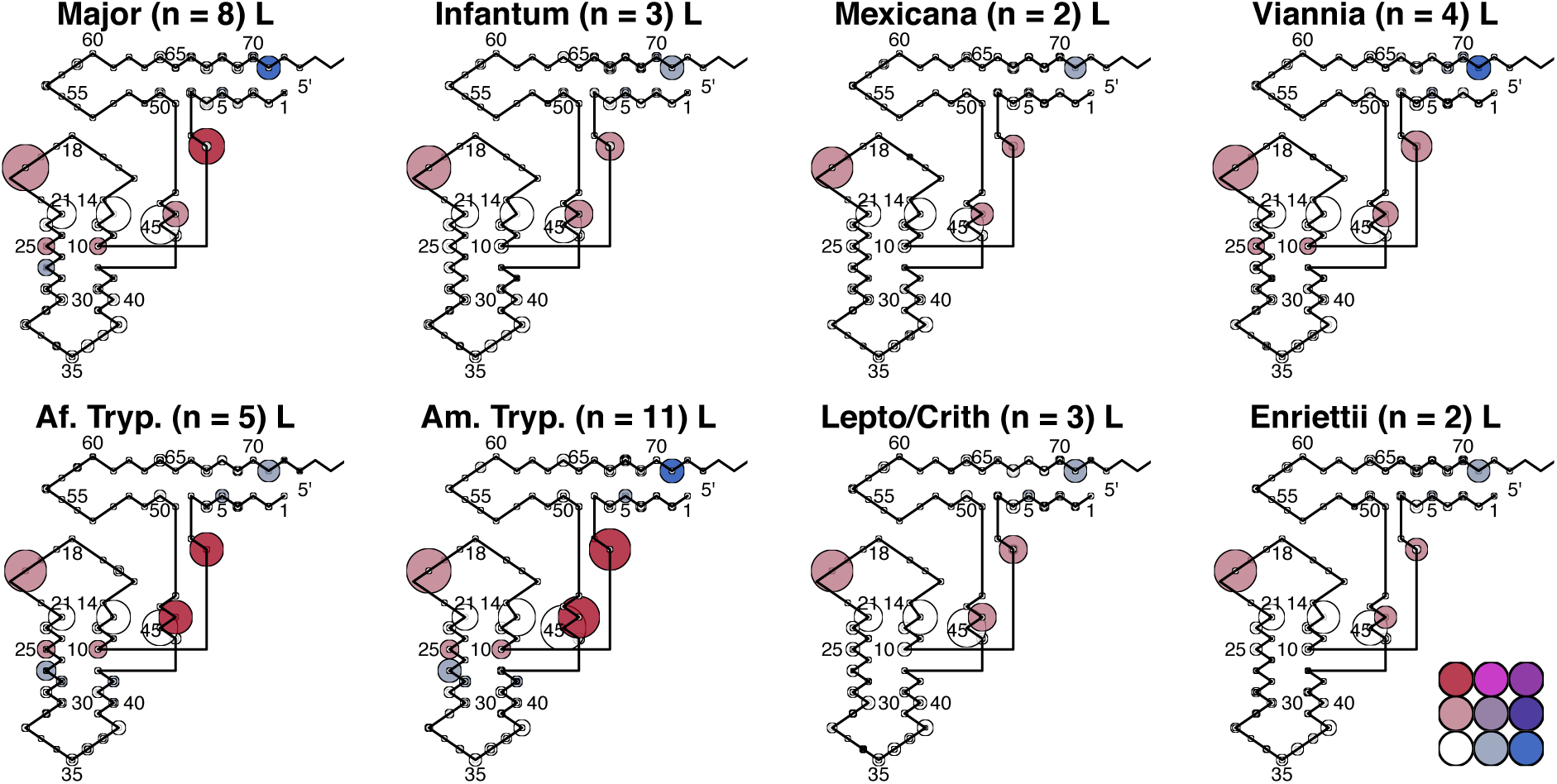
Leu Bubbleplots

**Supplementary Figure 2.**
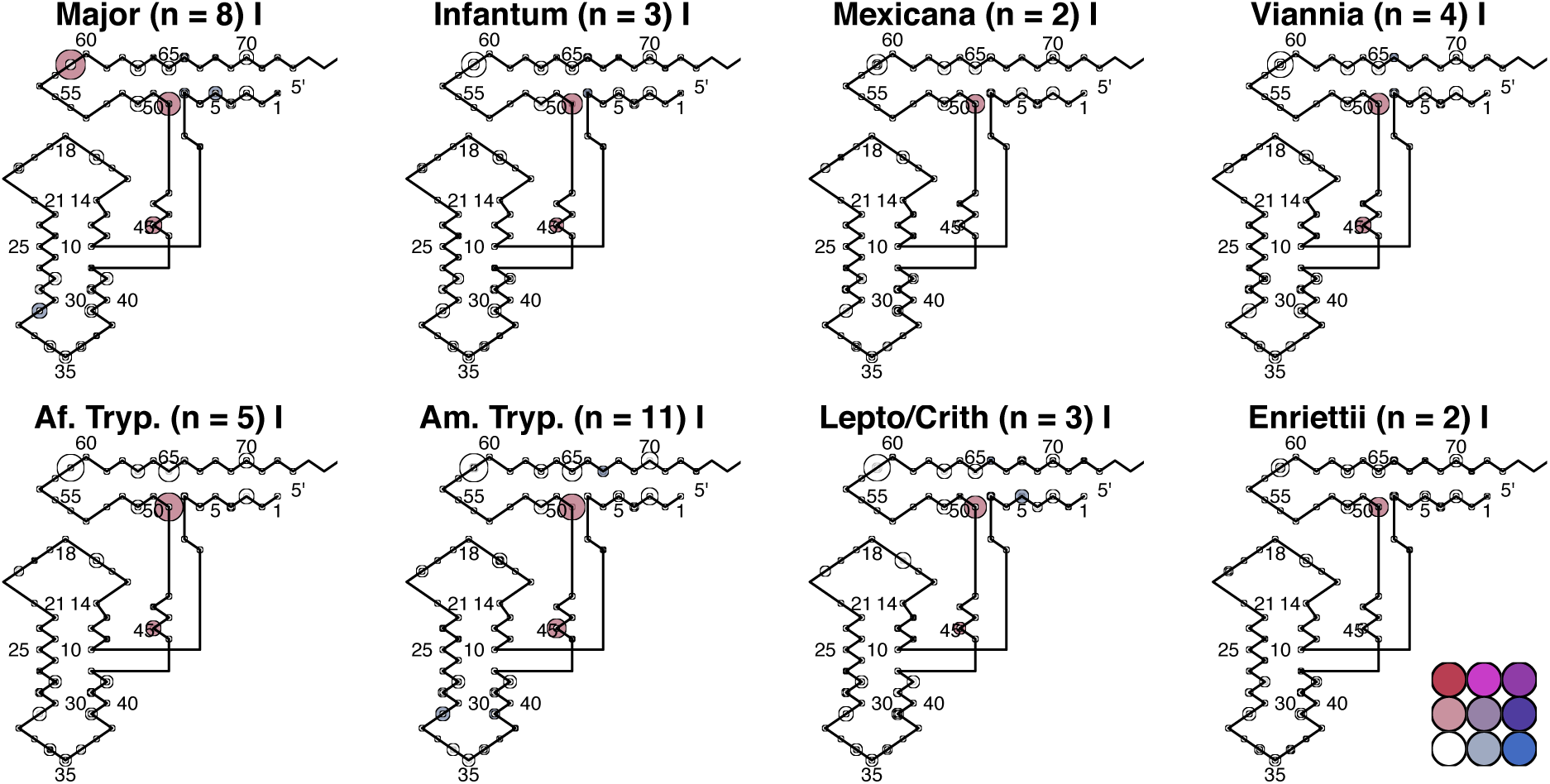
Ile Bubbleplots

**Supplementary Figure 3.**
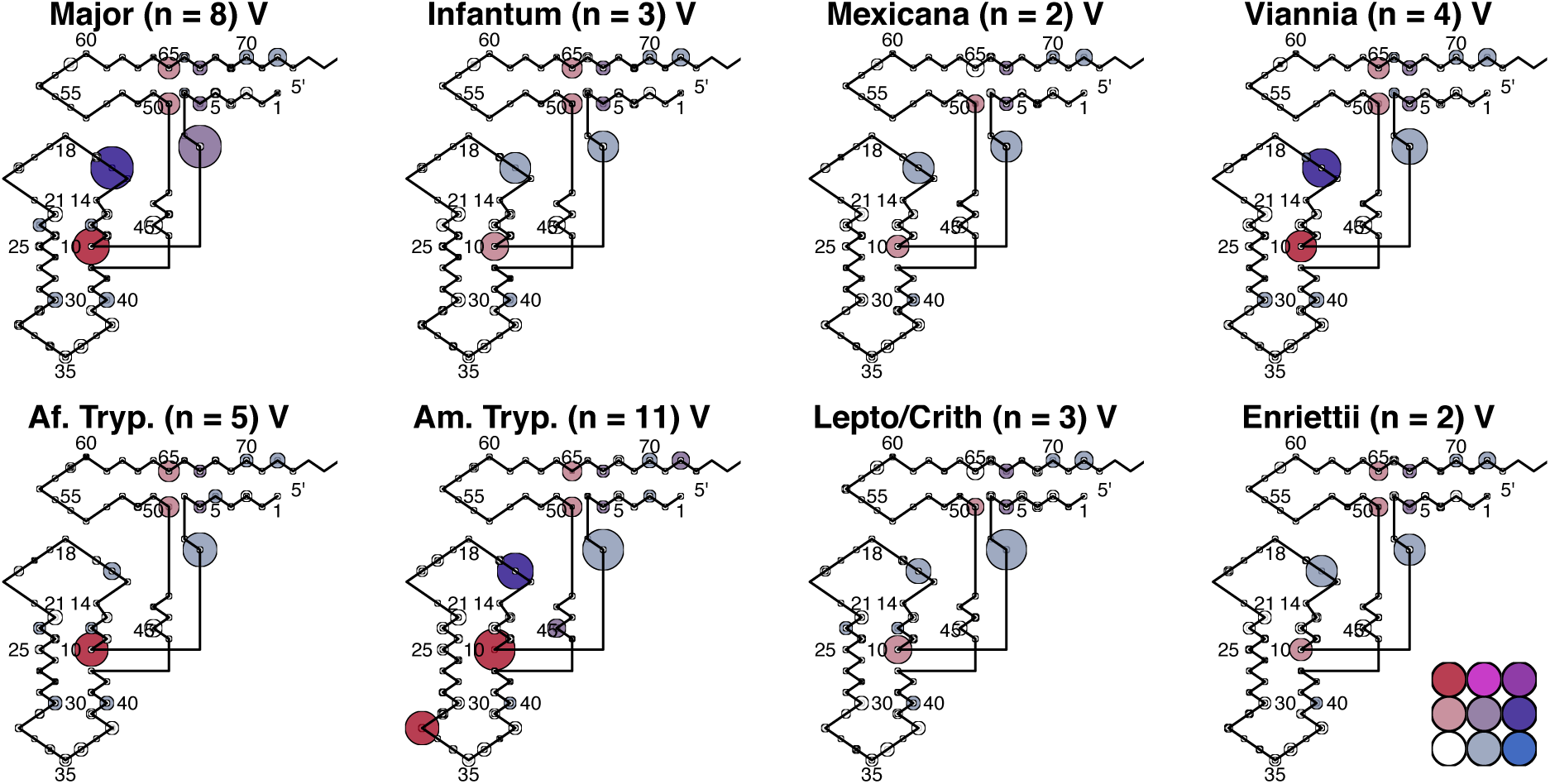
Val Bubbleplots

**Supplementary Figure 4.**
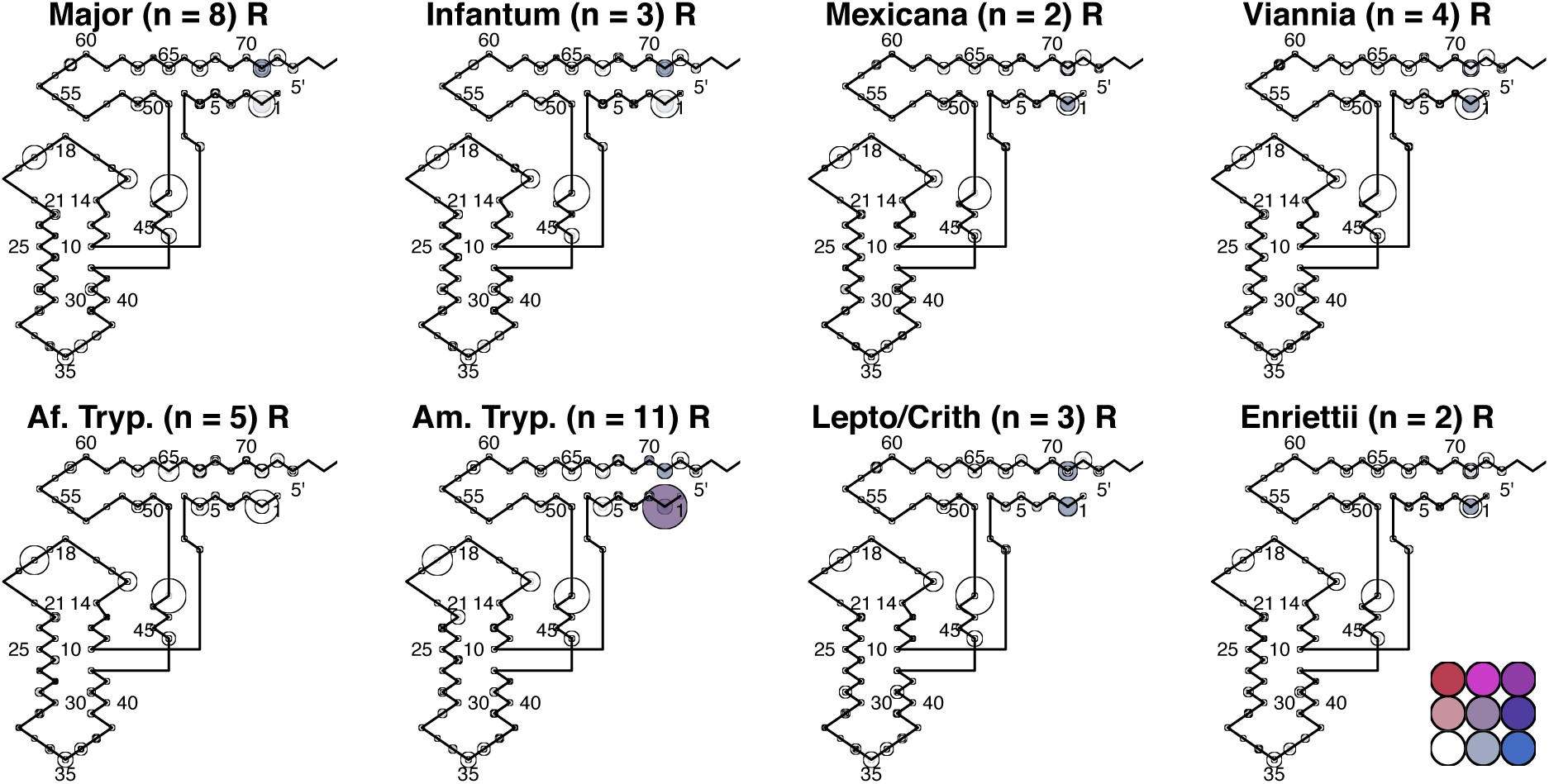
Arg Bubbleplots

**Supplementary Figure 5.**
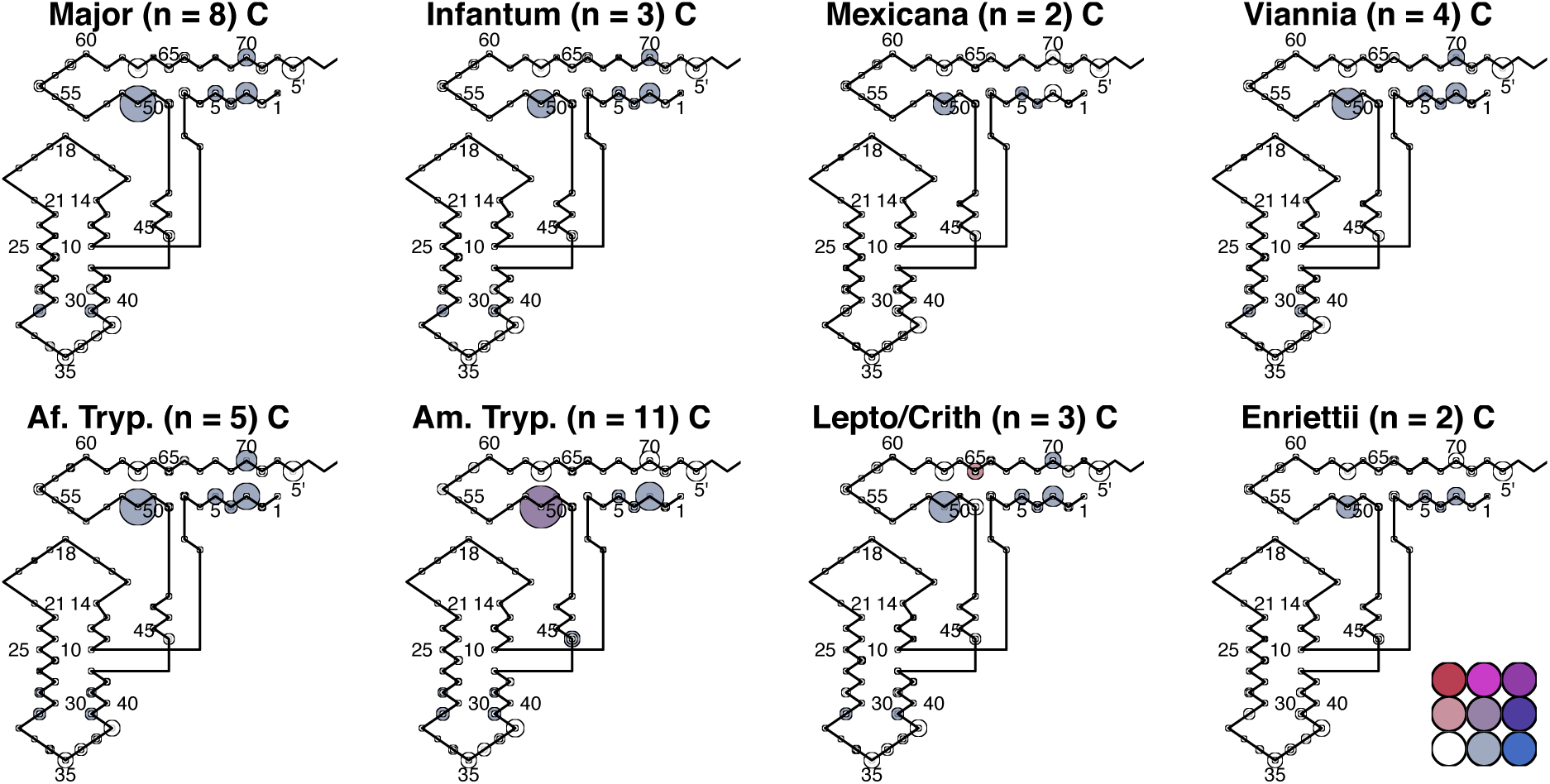
Cys Bubbleplots

**Supplementary Figure 6.**
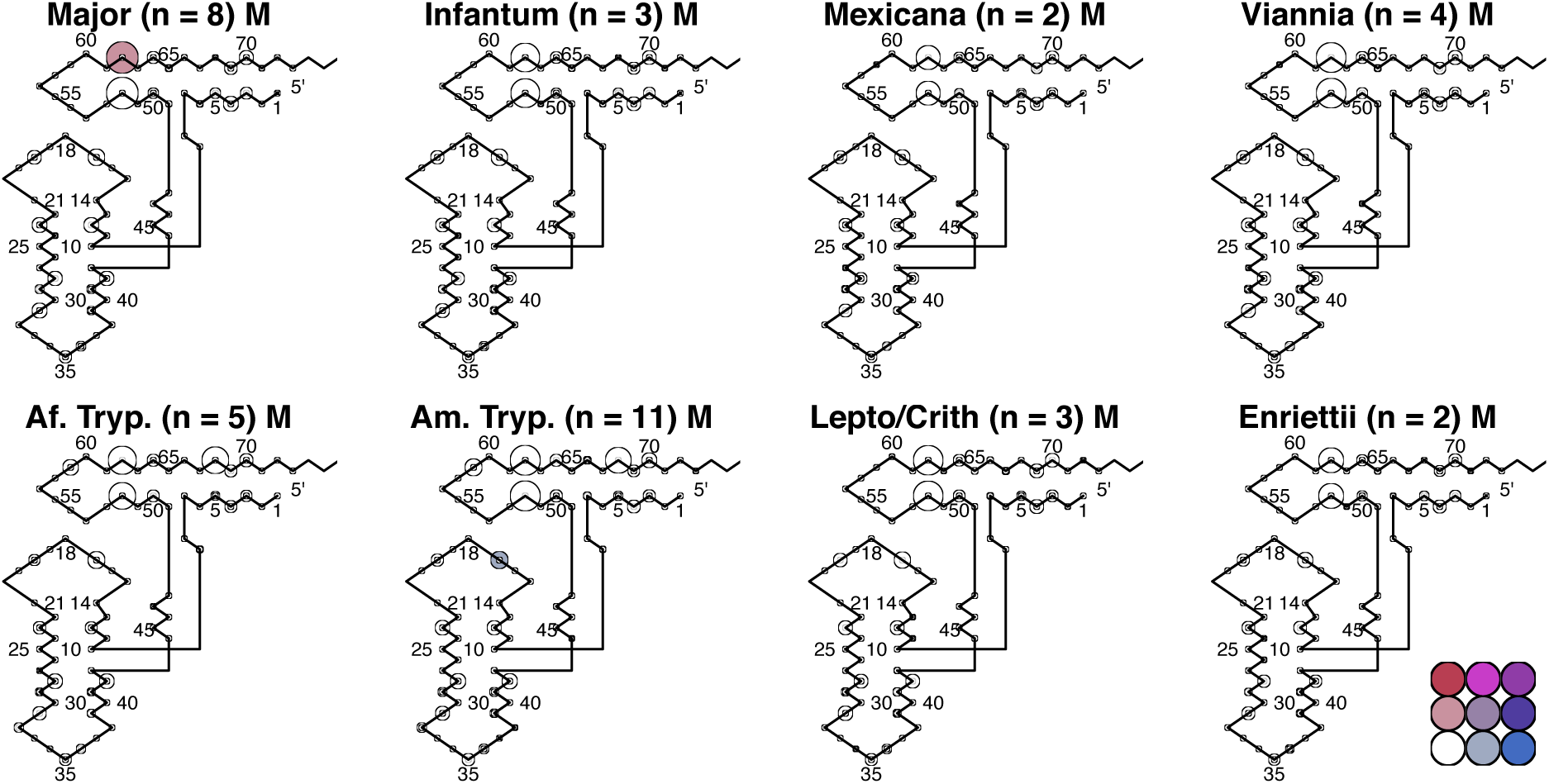
Met Bubbleplots

**Supplementary Figure 7.**
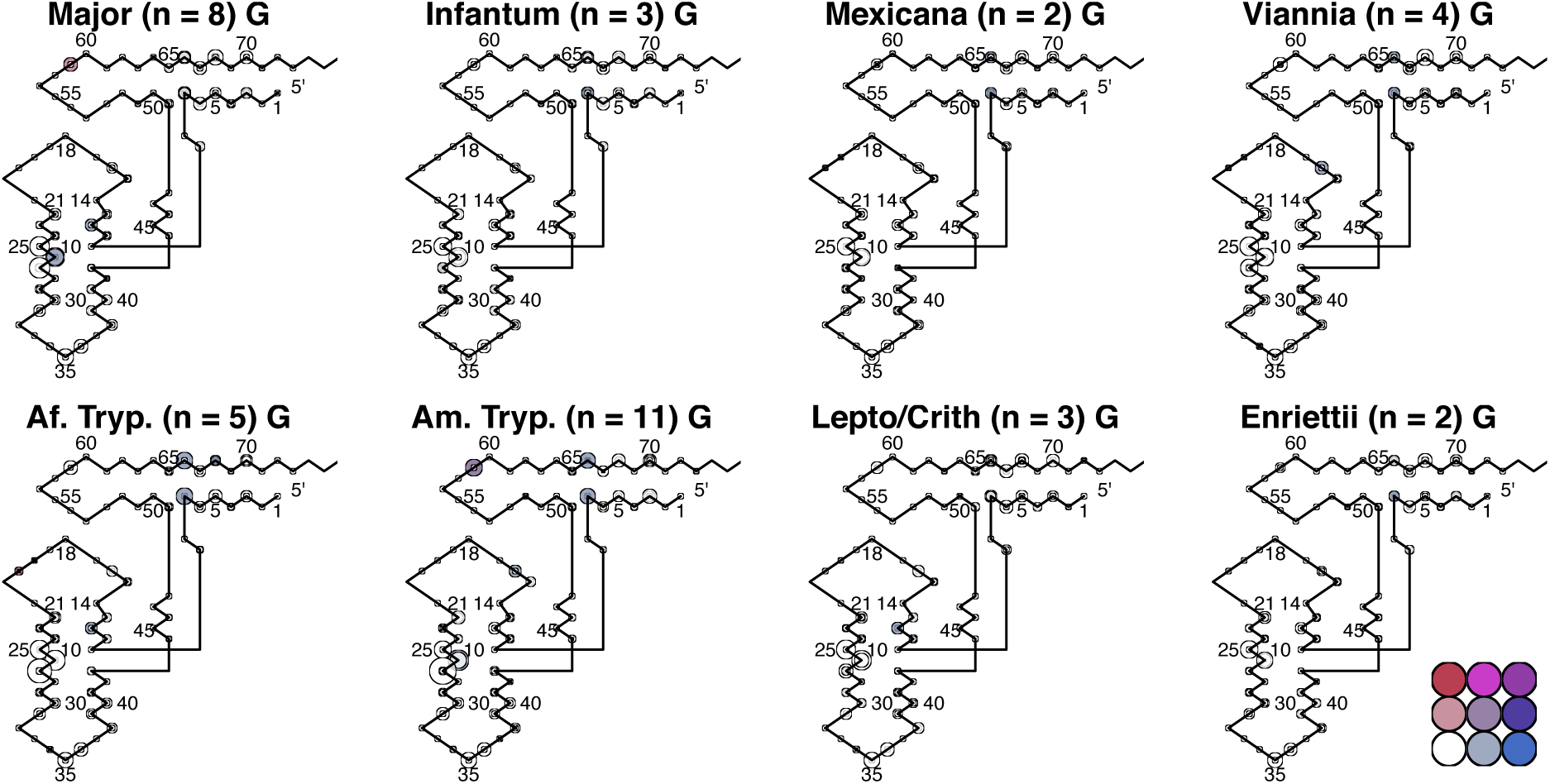
Glu Bubbleplots

**Supplementary Figure 8.**
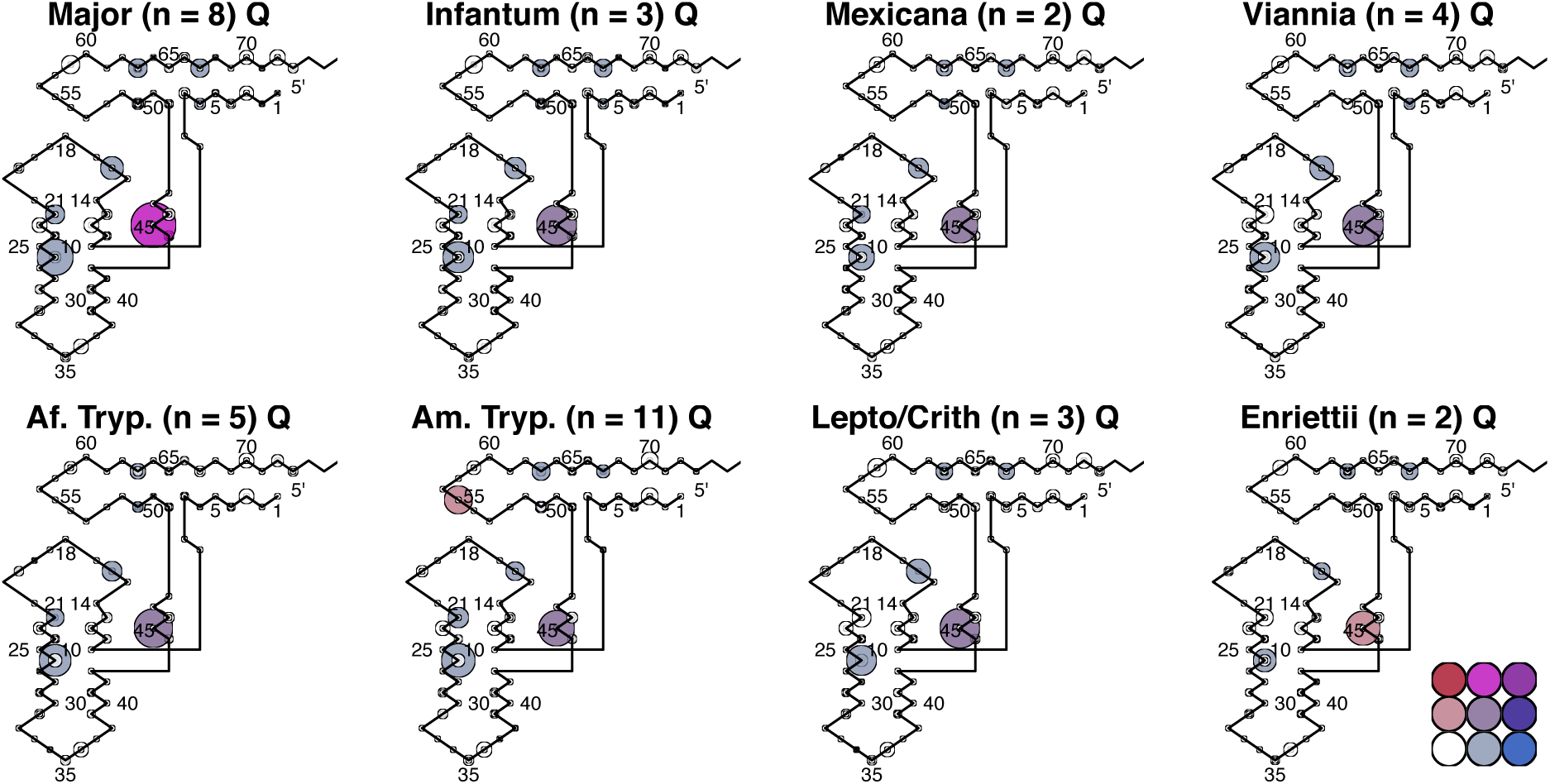
Gln Bubbleplots

**Supplementary Figure 9.**
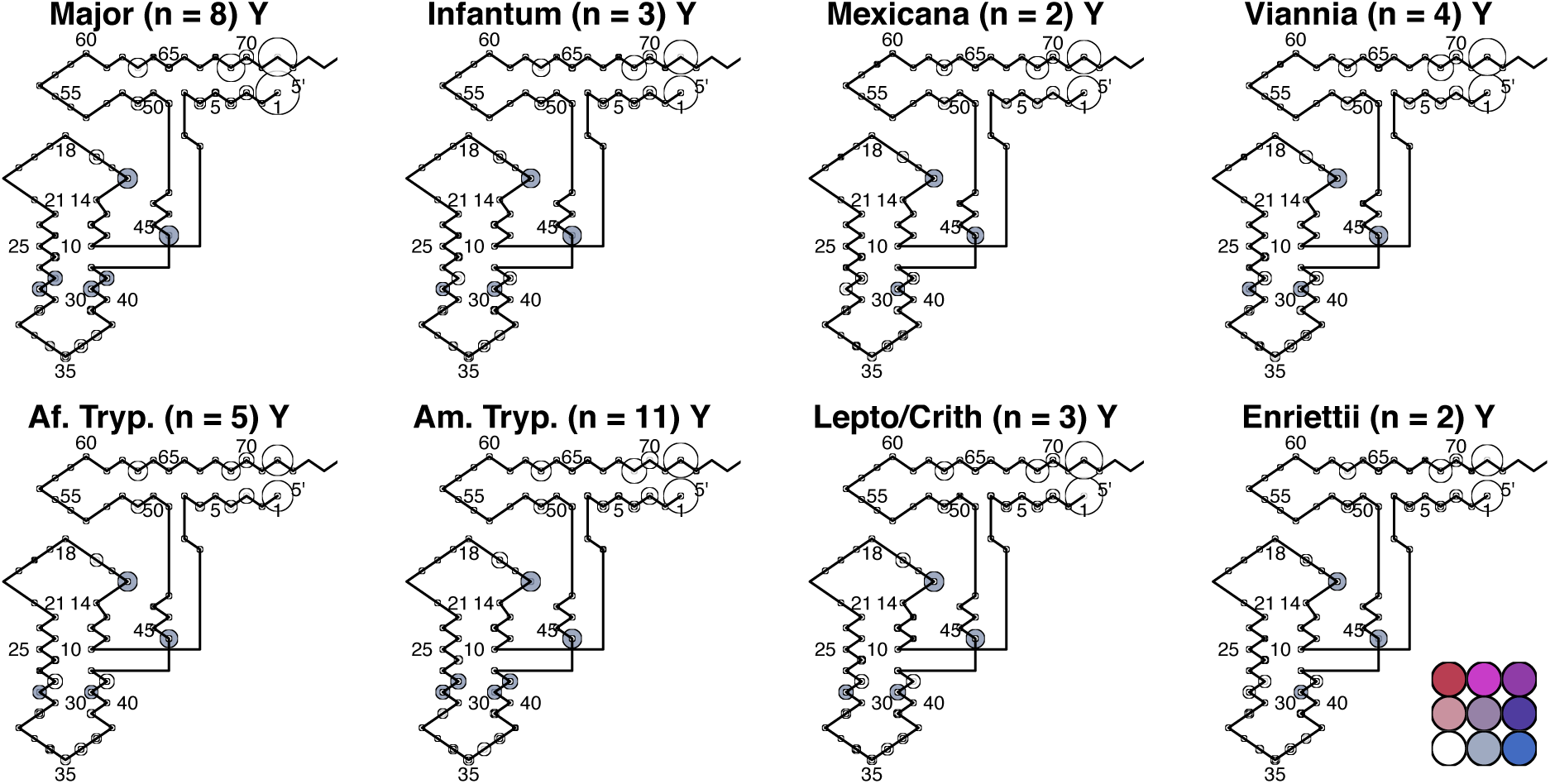
Tyr Bubbleplots

**Supplementary Figure 10.**
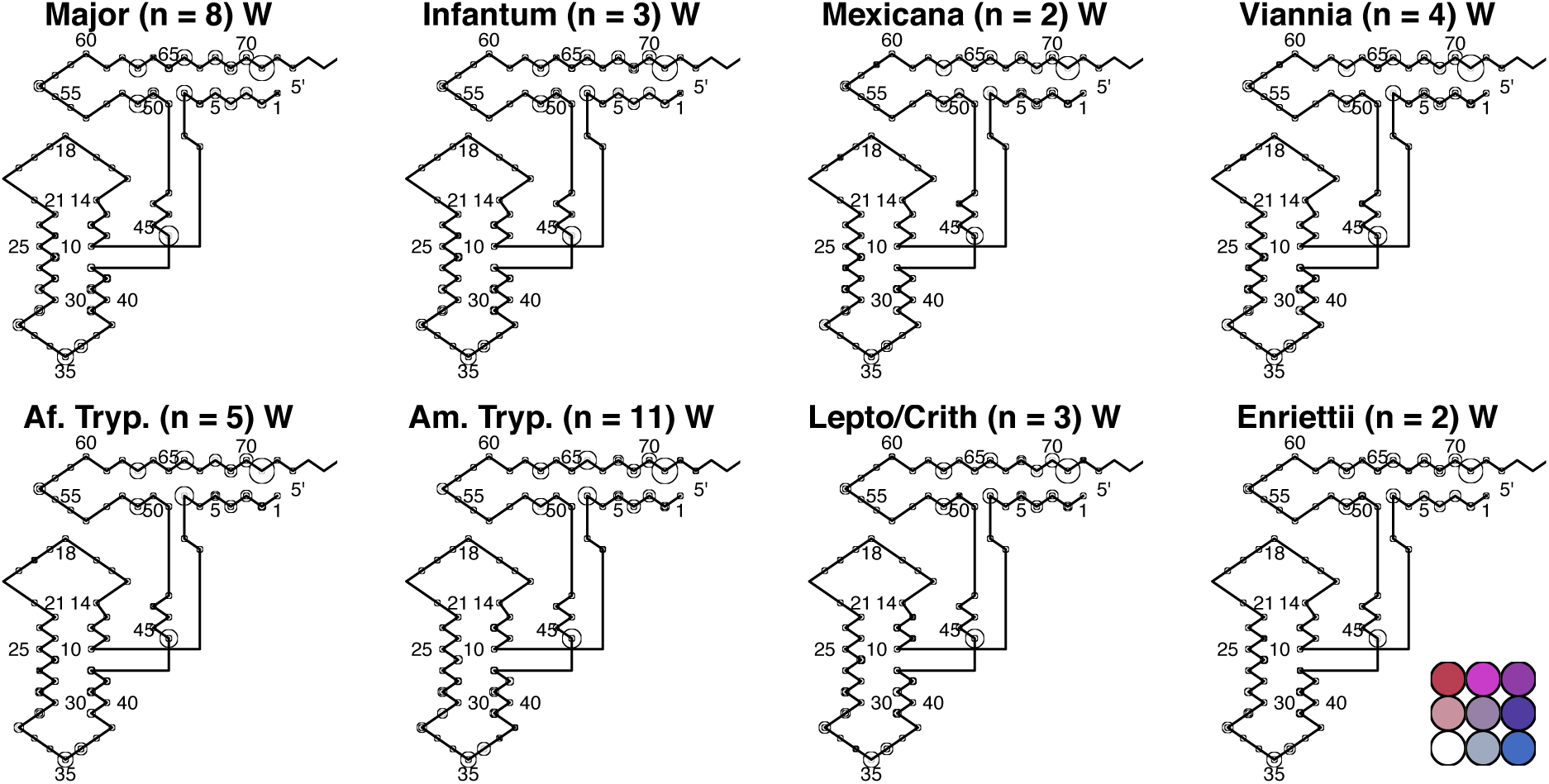
Trp Bubbleplots

**Supplementary Figure 11.**
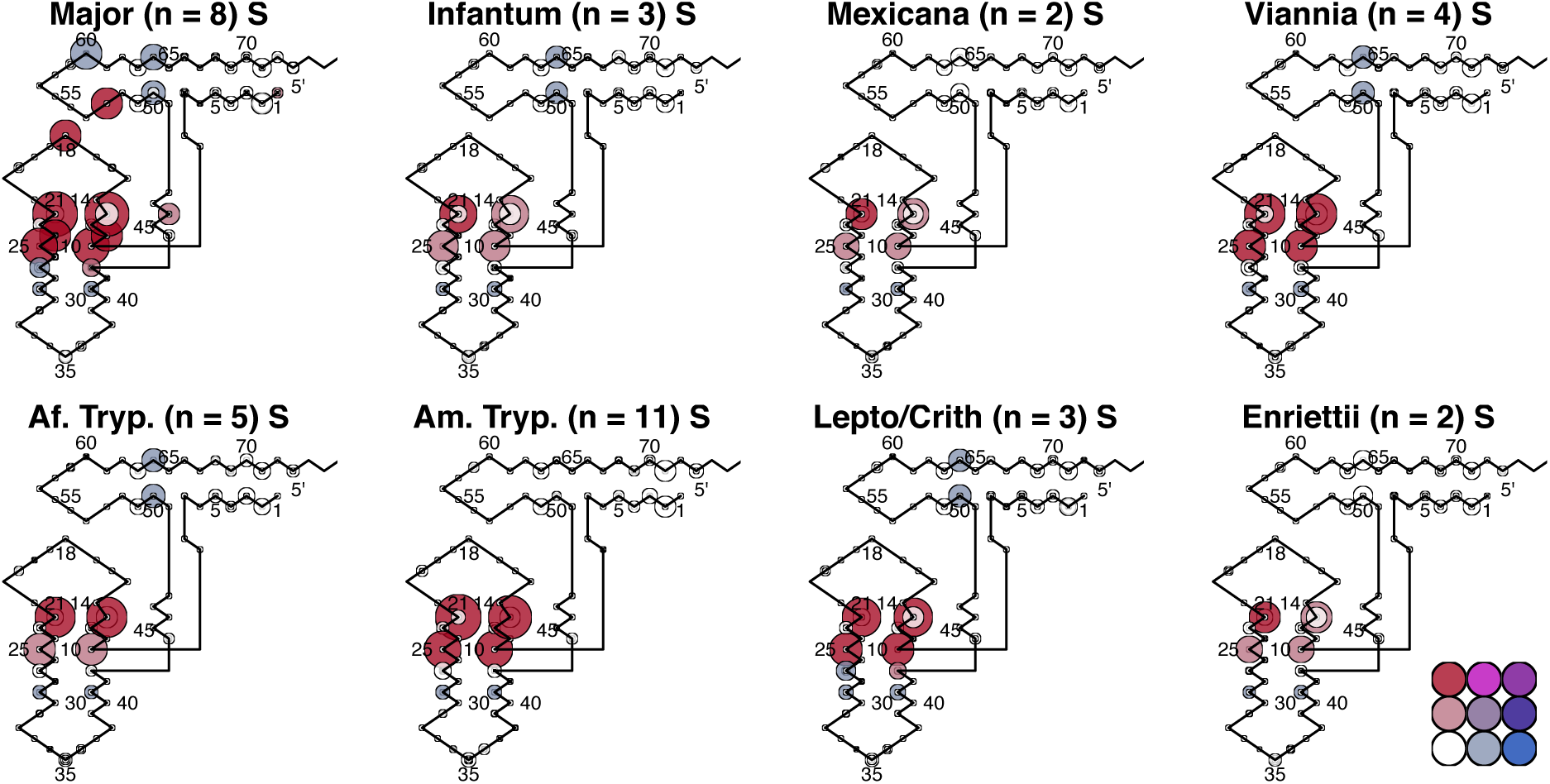
Ser Bubbleplots

**Supplementary Figure 12.**
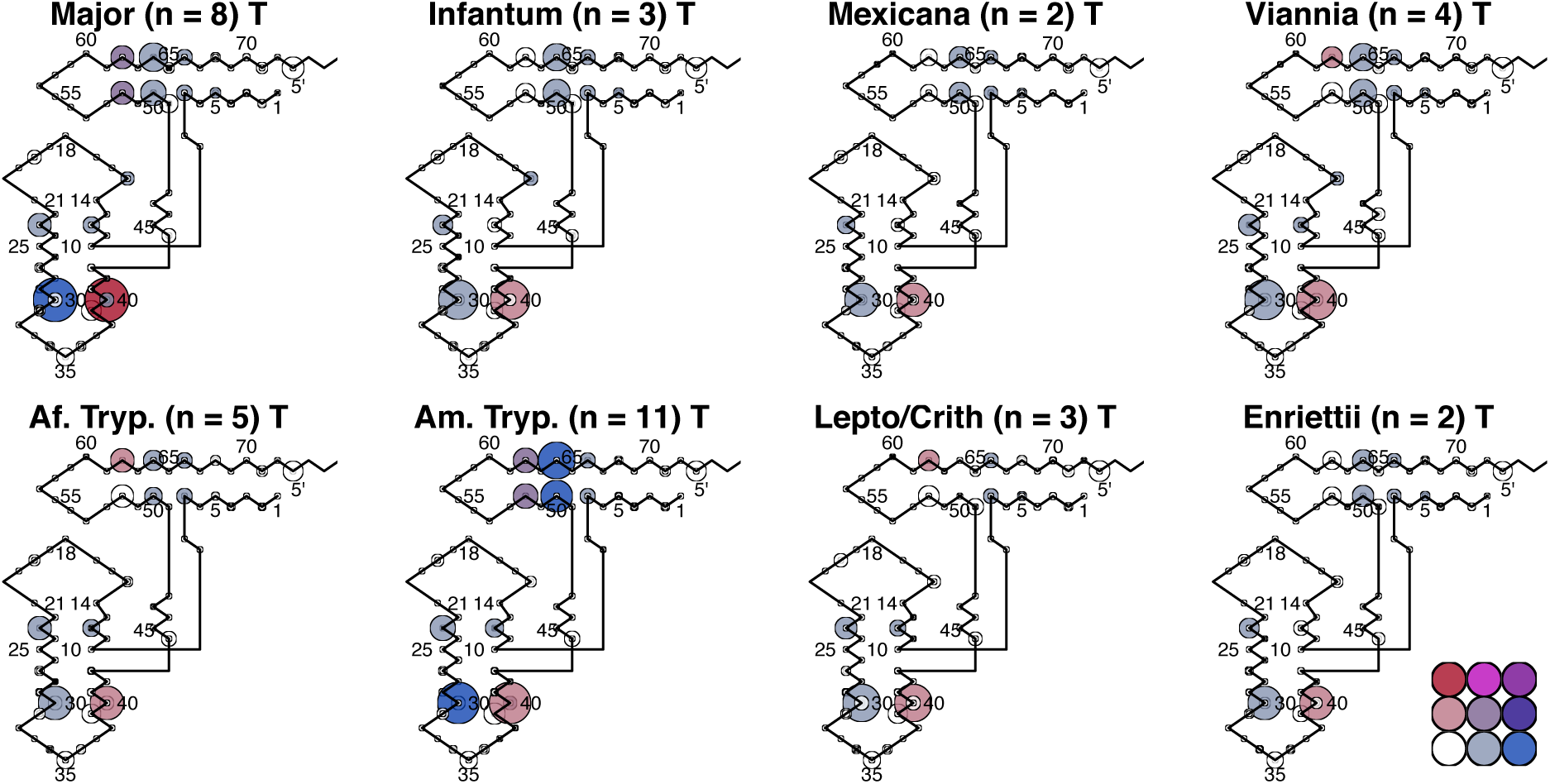
Thr Bubbleplots

**Supplementary Figure 13.**
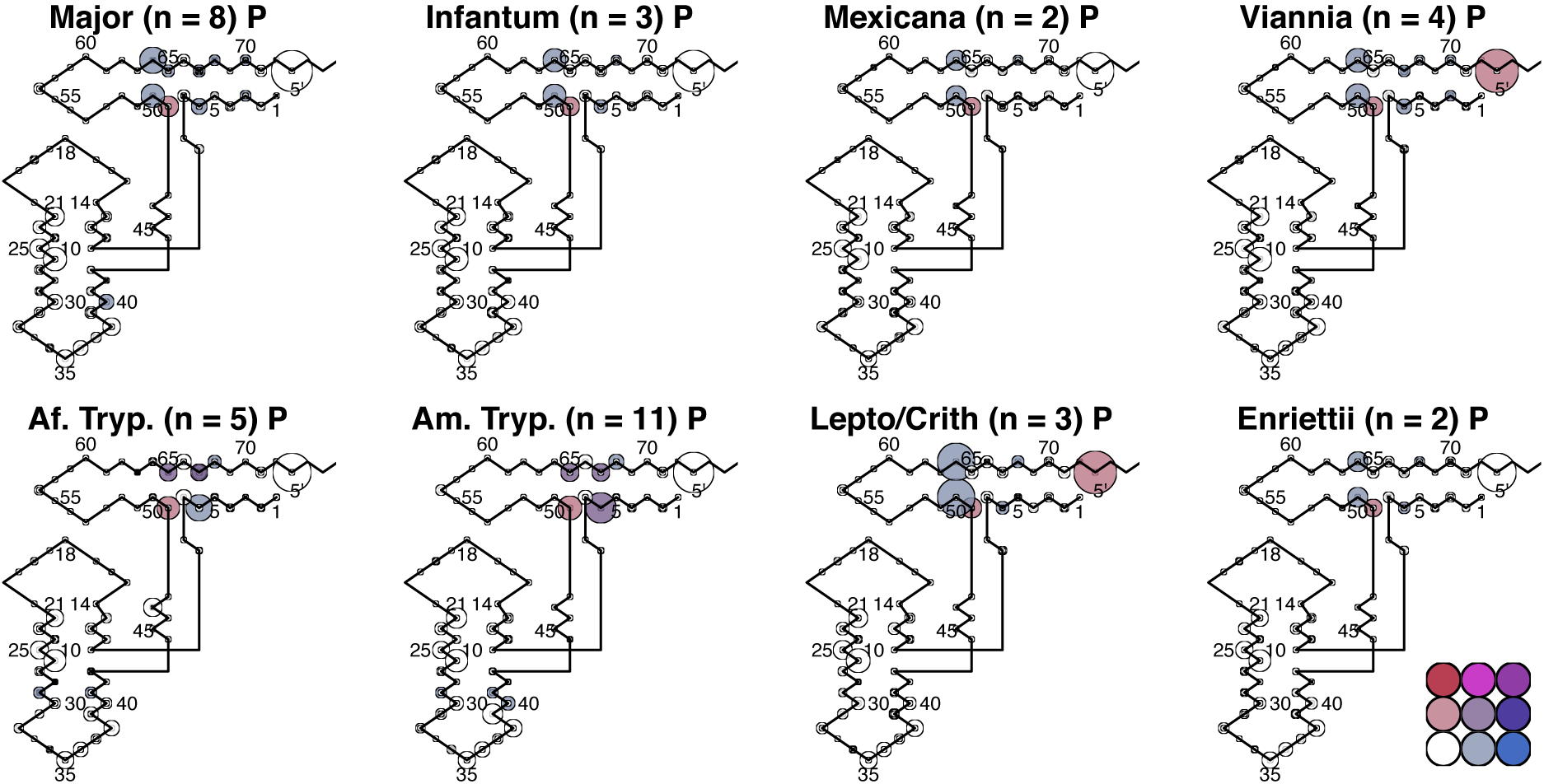
Pro Bubbleplots

**Supplementary Figure 14.**
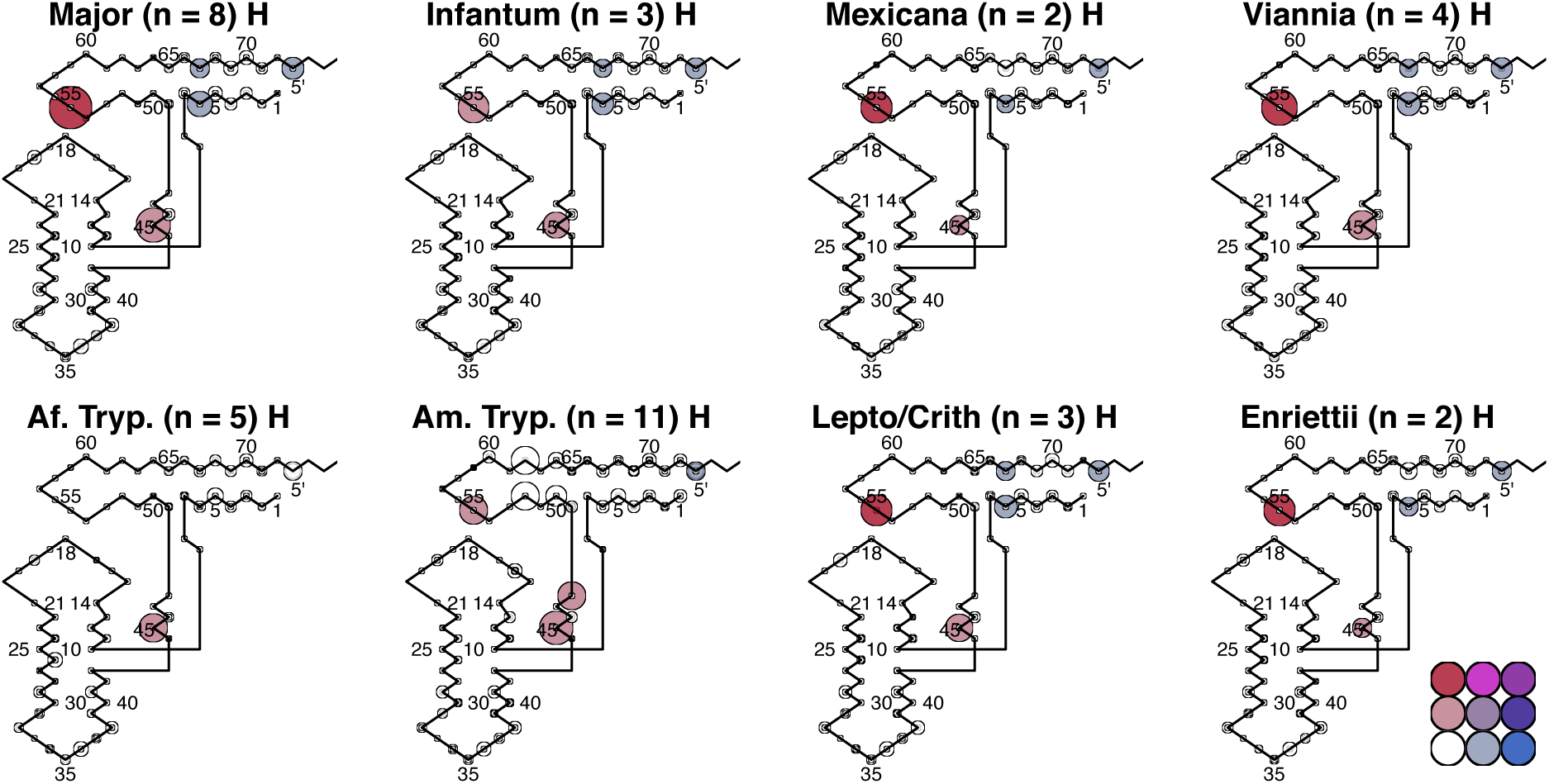
His Bubbleplots

**Supplementary Figure 15.**
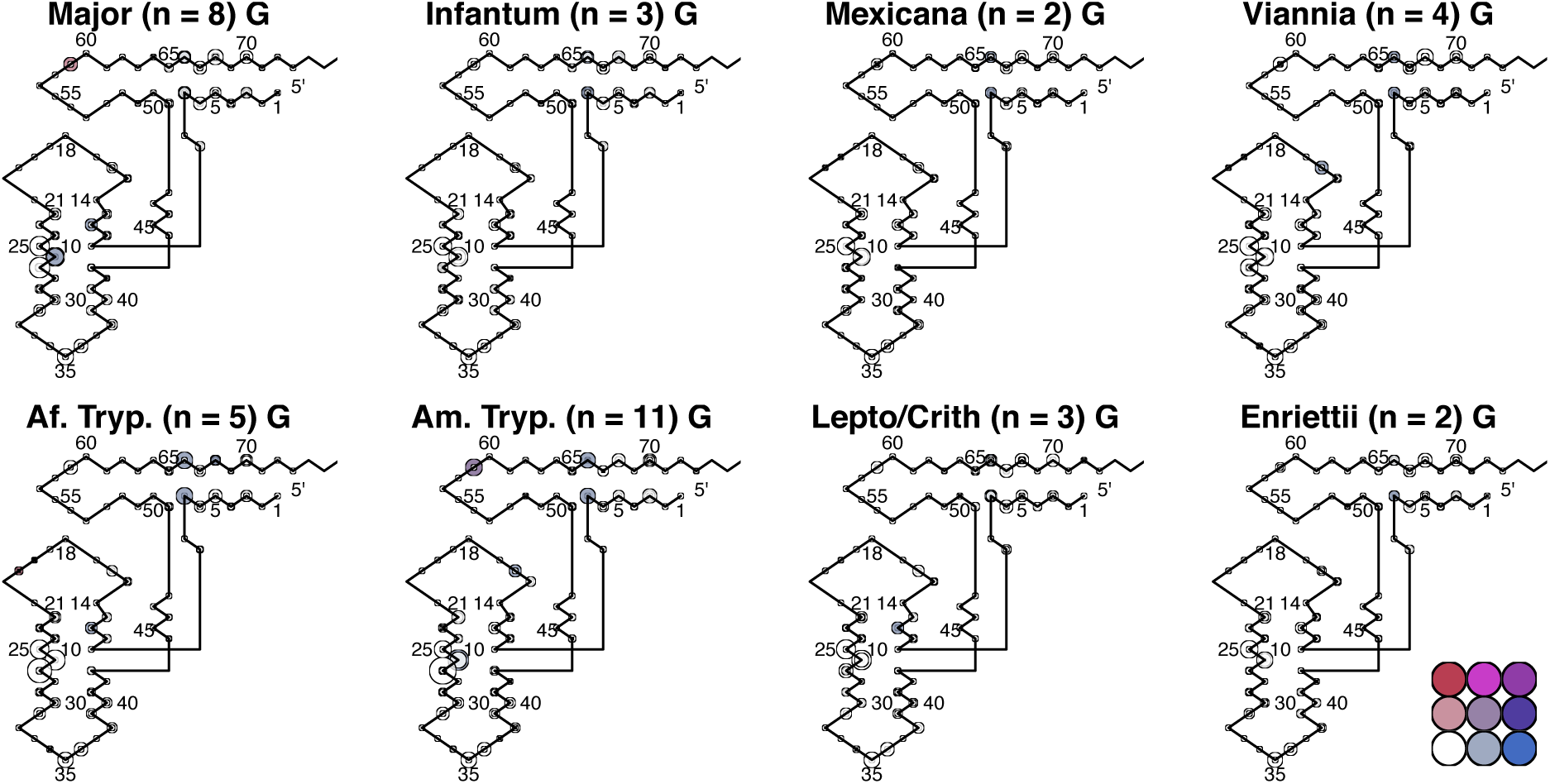
Gly Bubbleplots

**Supplementary Figure 16.**
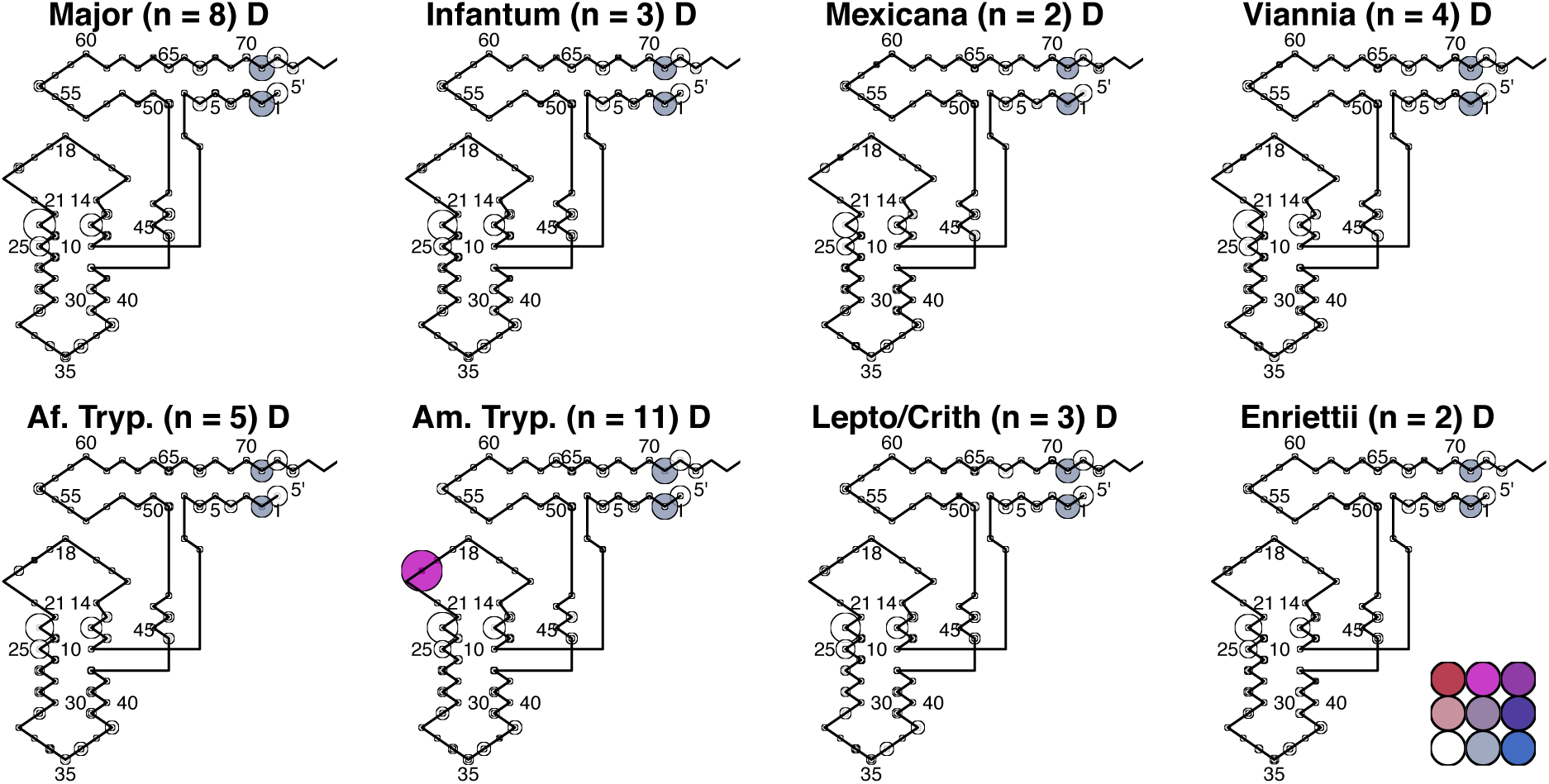
Asp Bubbleplots

**Supplementary Figure 17.**
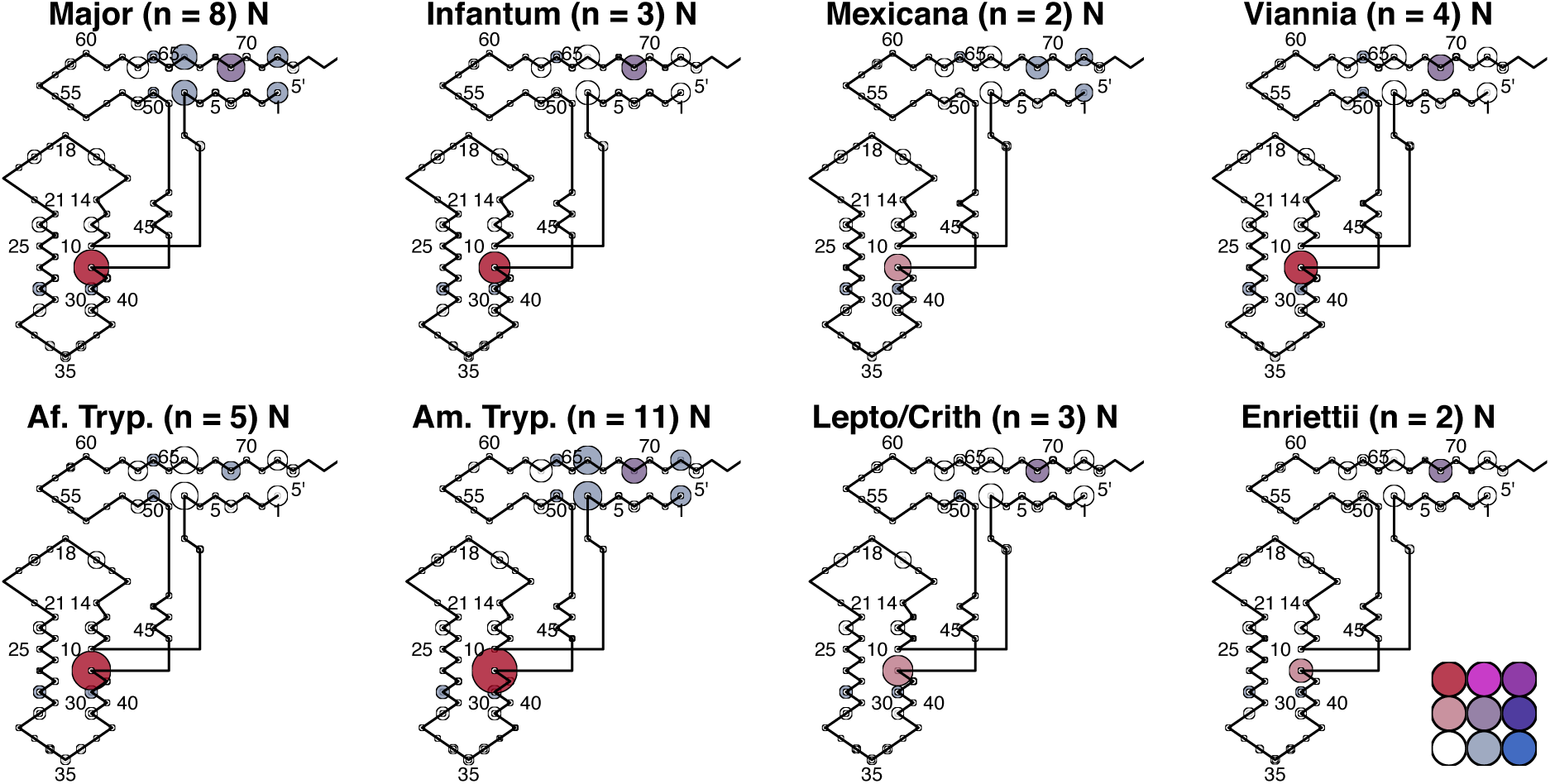
Asn bubbleplots

**Supplementary Figure 18.**
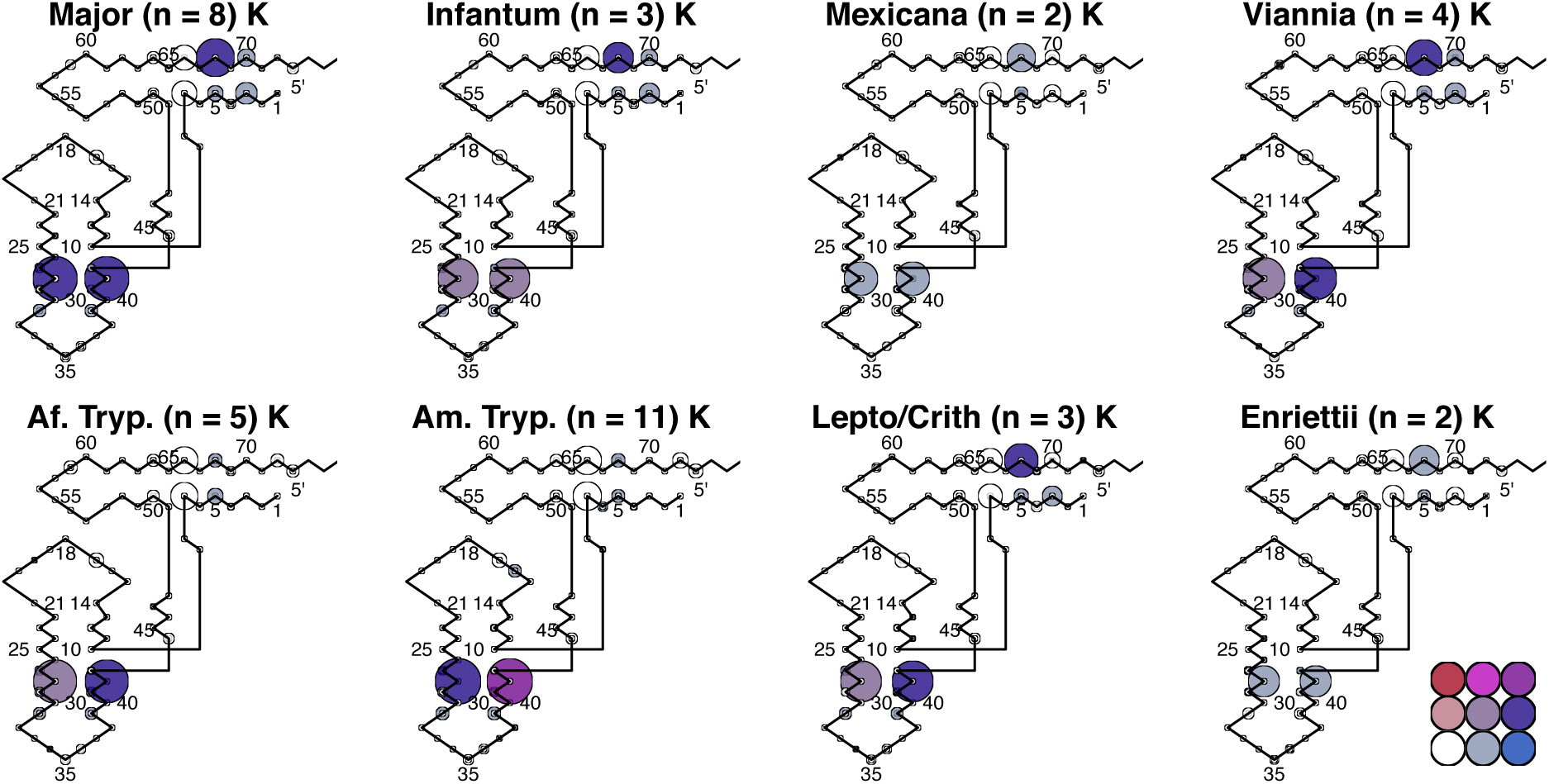
Lys Bubbleplots

**Supplementary Figure 19.**
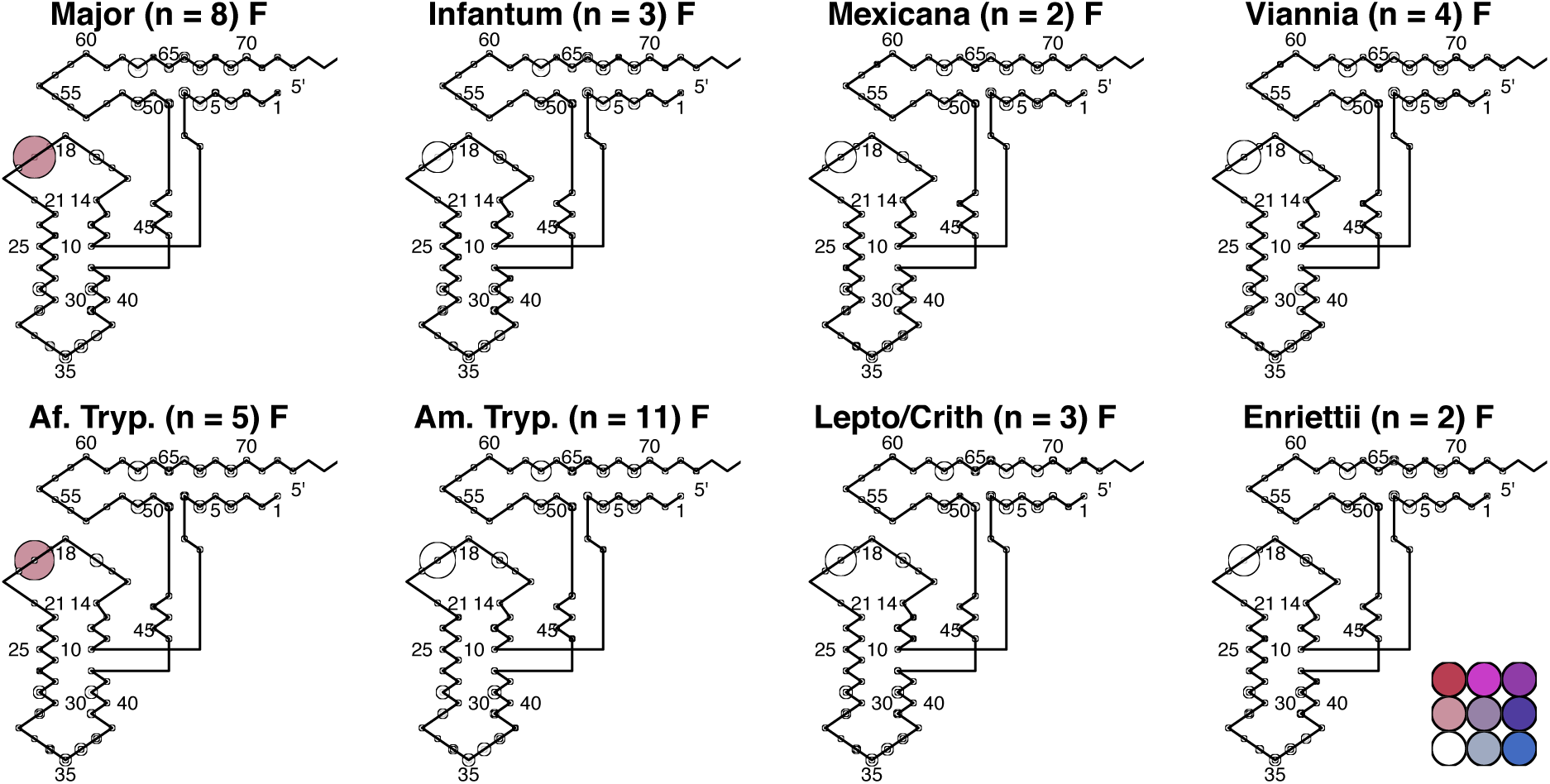
Phe Bubbleplots

**Supplementary Figure 20.**
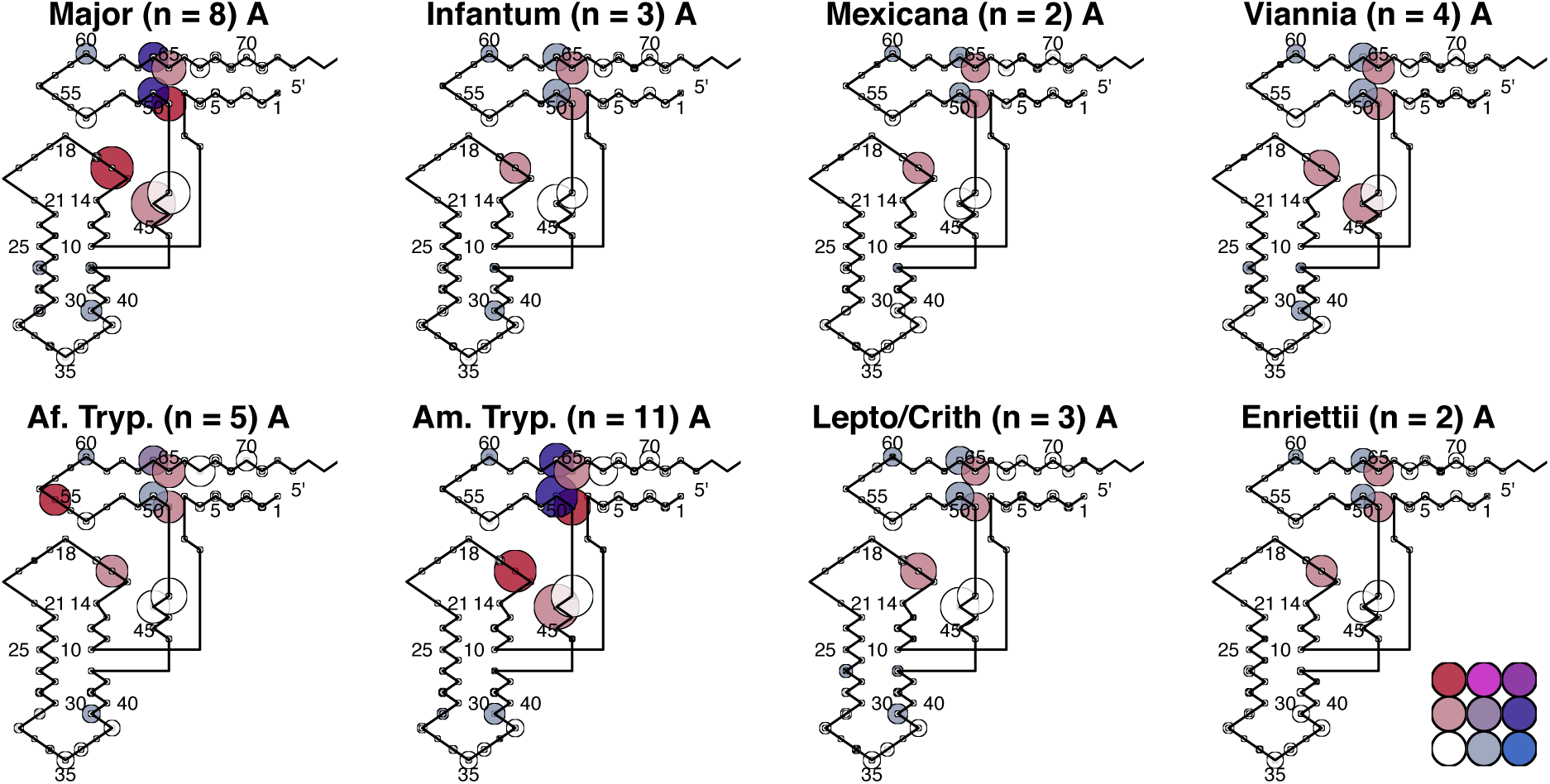
Ala Bubbleplots

**Supplementary Figure 21.**
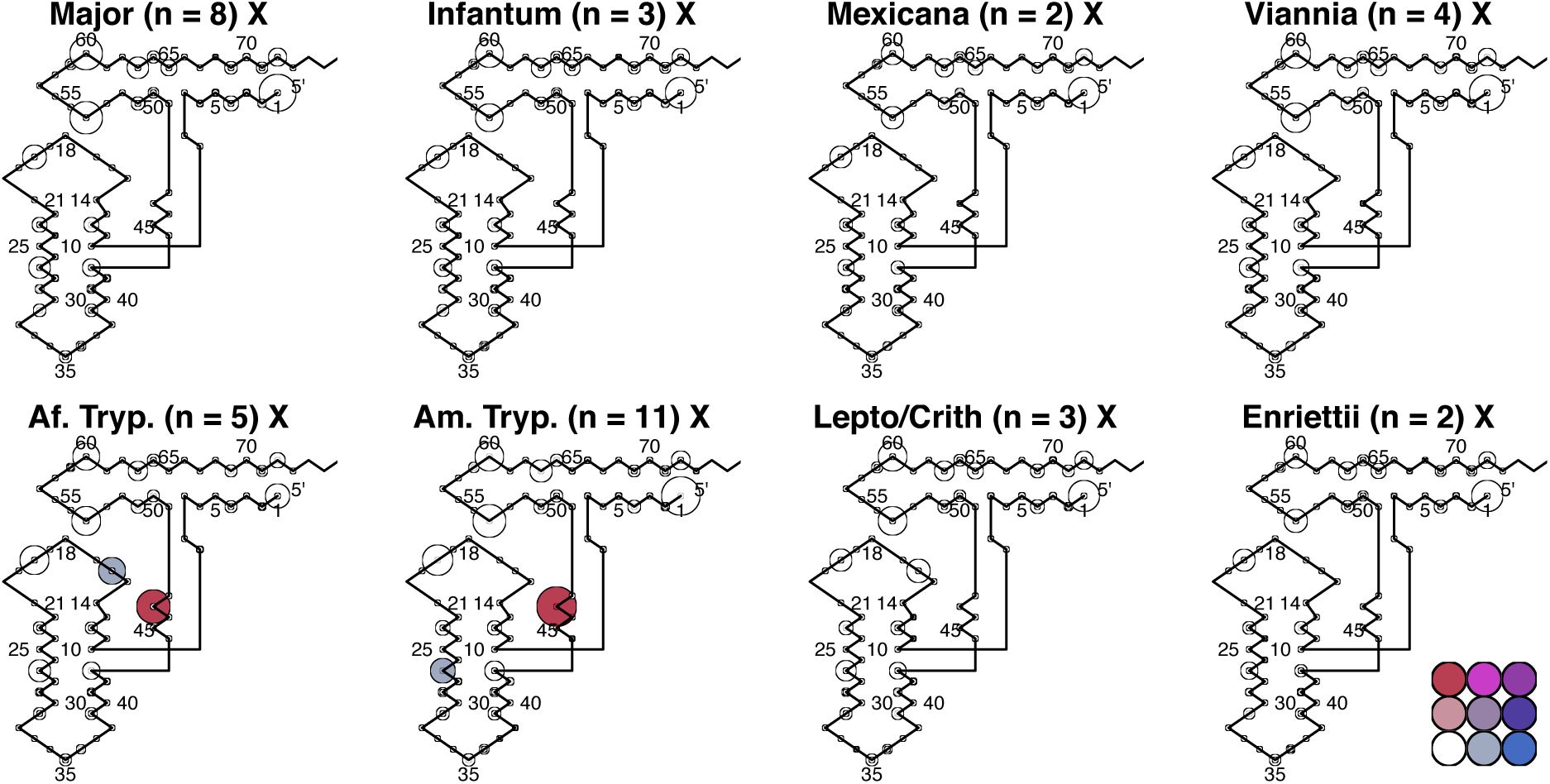
iMet Bubbleplots

**Supplementary Figure 22.**
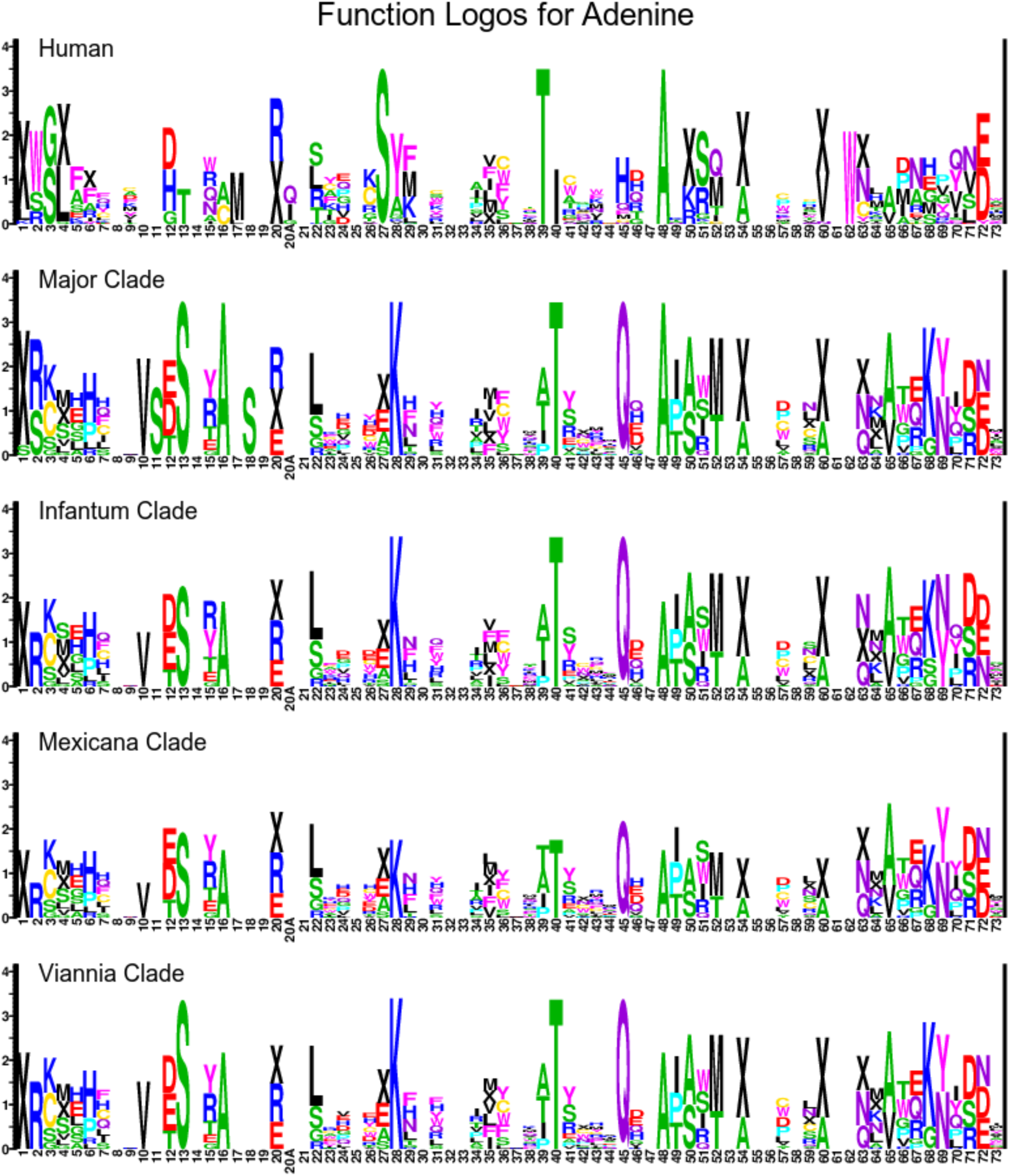
Adenine Function Logos, part I

**Supplementary Figure 23.**
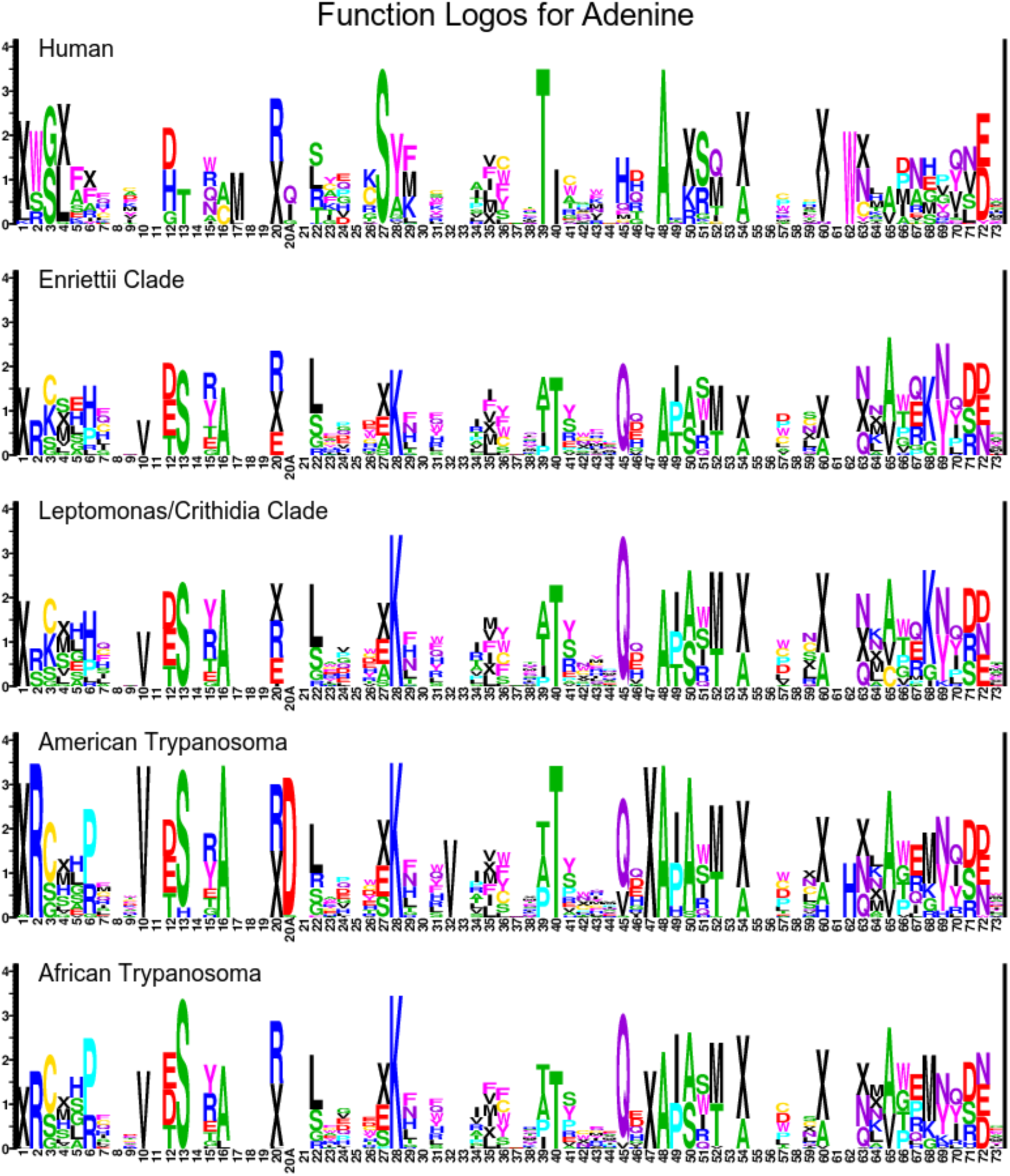
Adenine Function Logos, part II

**Supplementary Figure 24.**
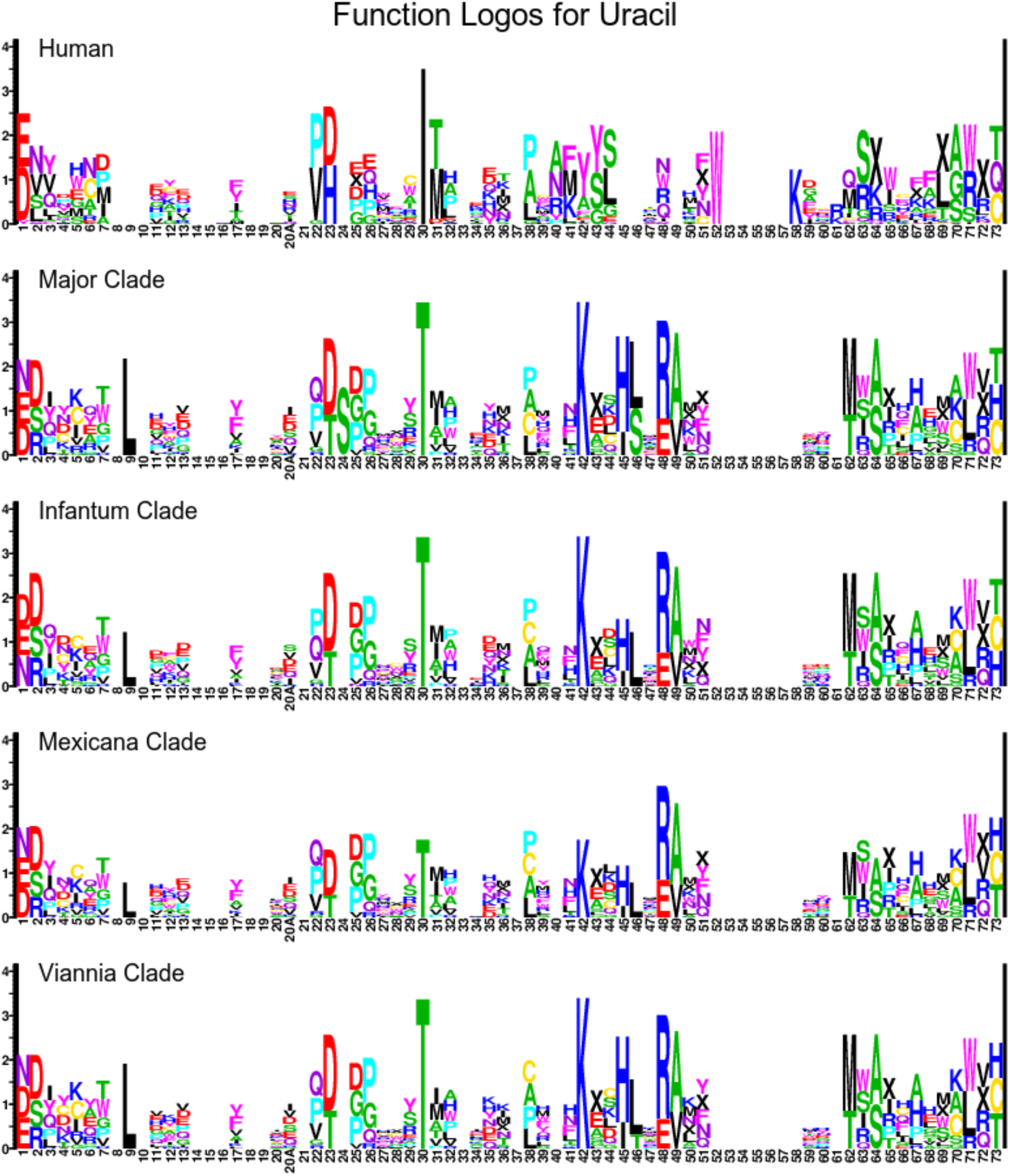
Uracil Function Logos, part I

**Supplementary Figure 25.**
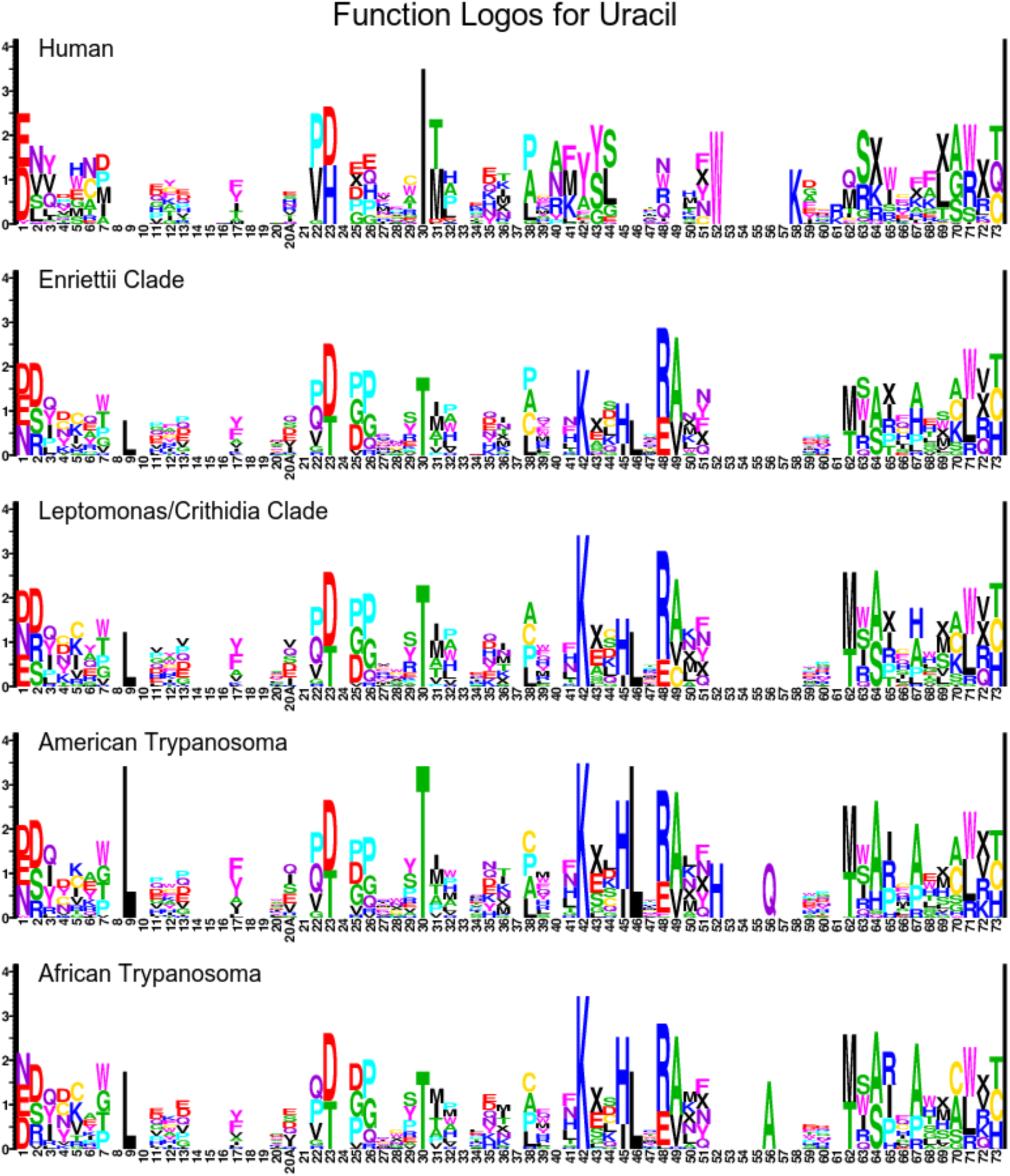
Uracil Function Logos, part II

**Supplementary Figure 26.**
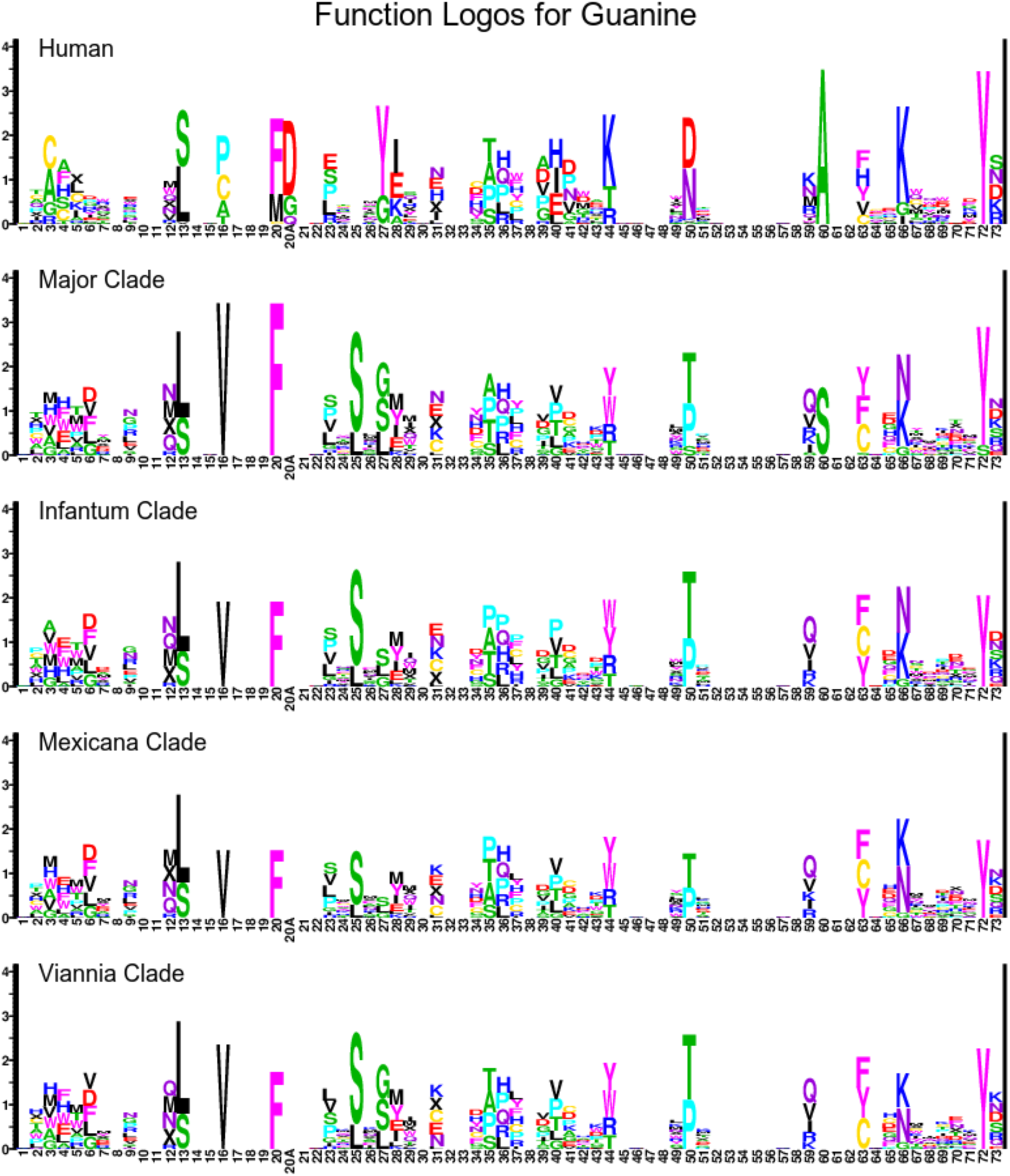
Guanine Function Logos, part I

**Supplementary Figure 27.**
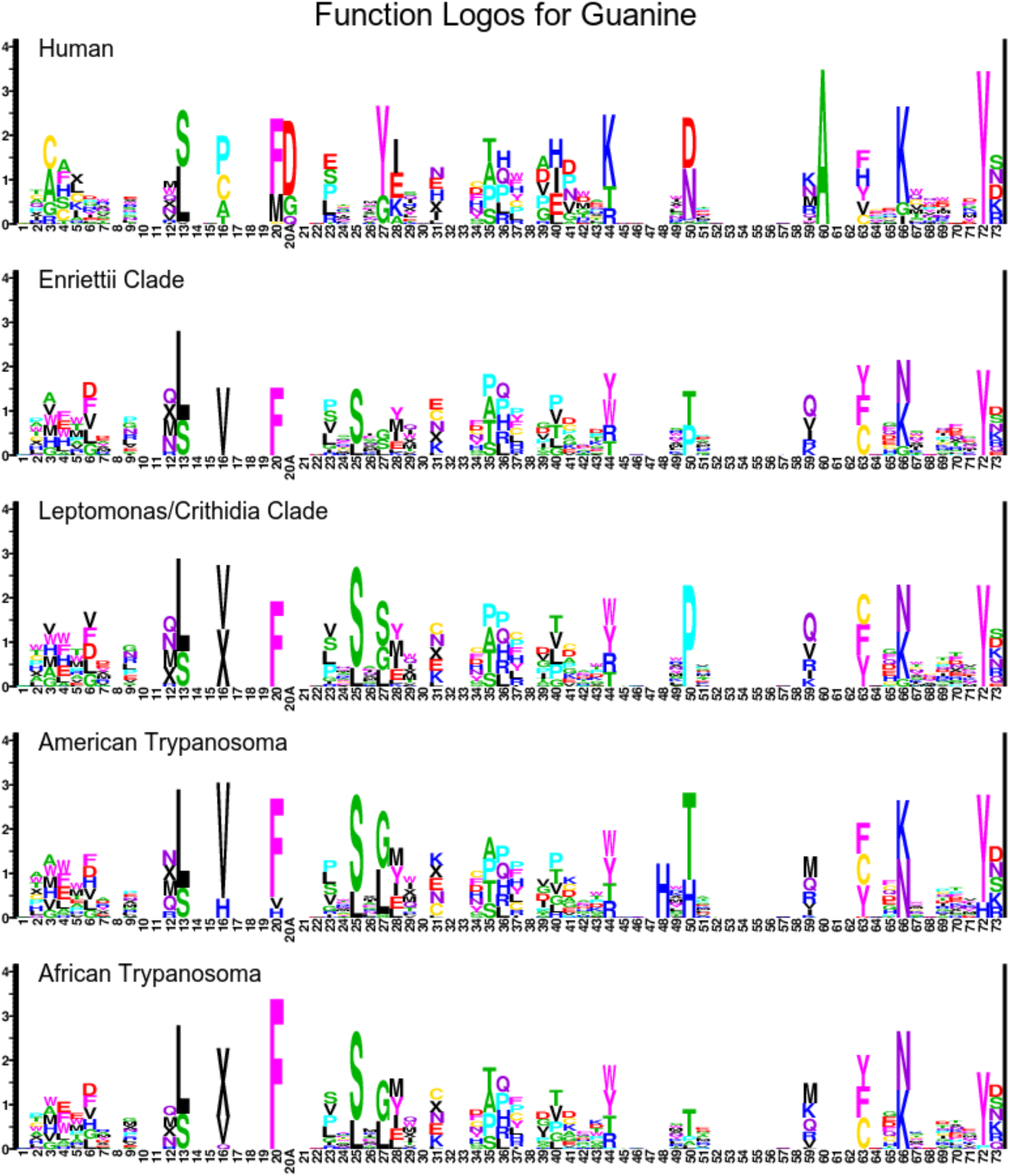
Guanine Function Logos, part II

**Supplementary Figure 28.**
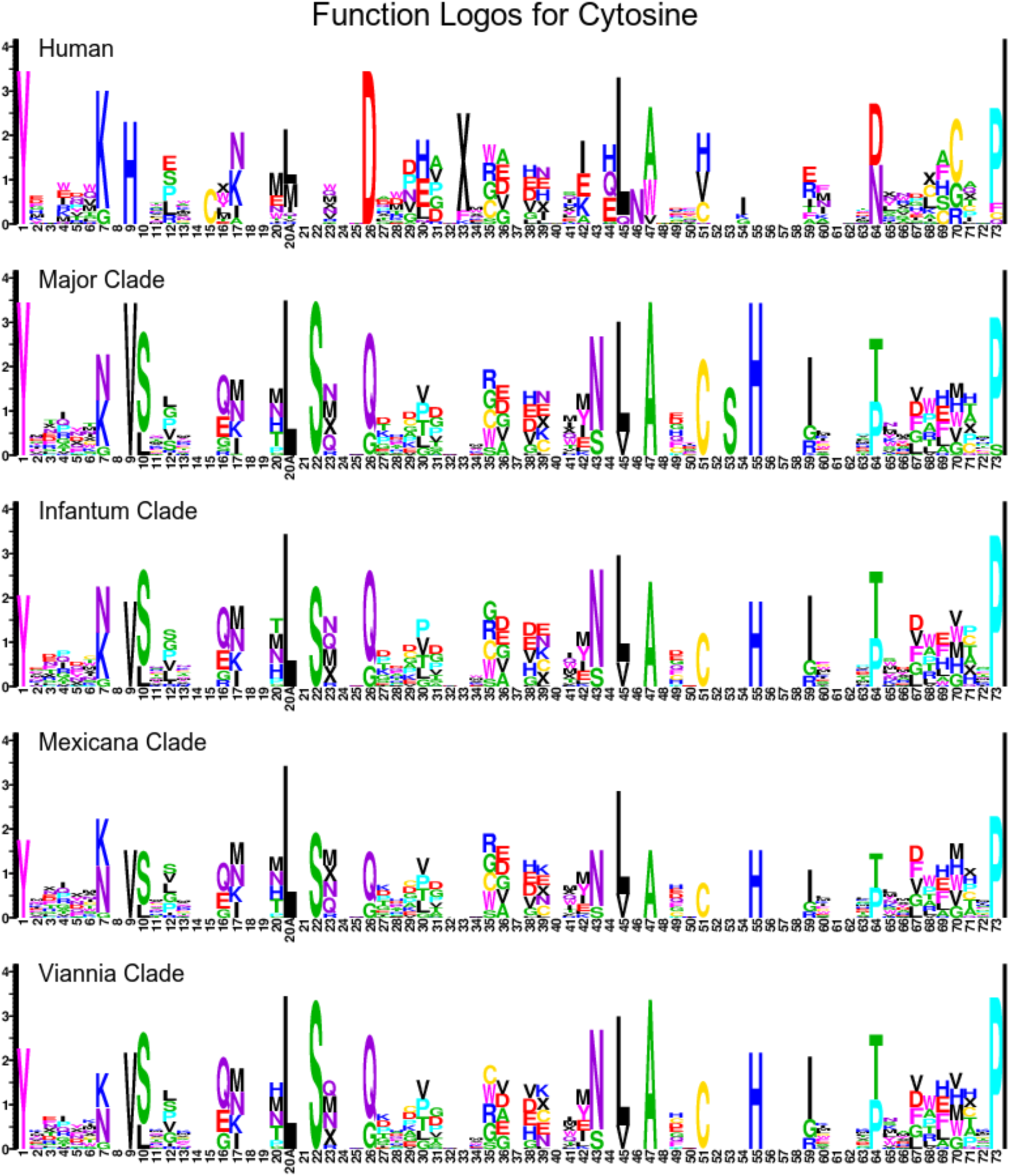
Cytosine Function Logos, part I

**Supplementary Figure 29.**
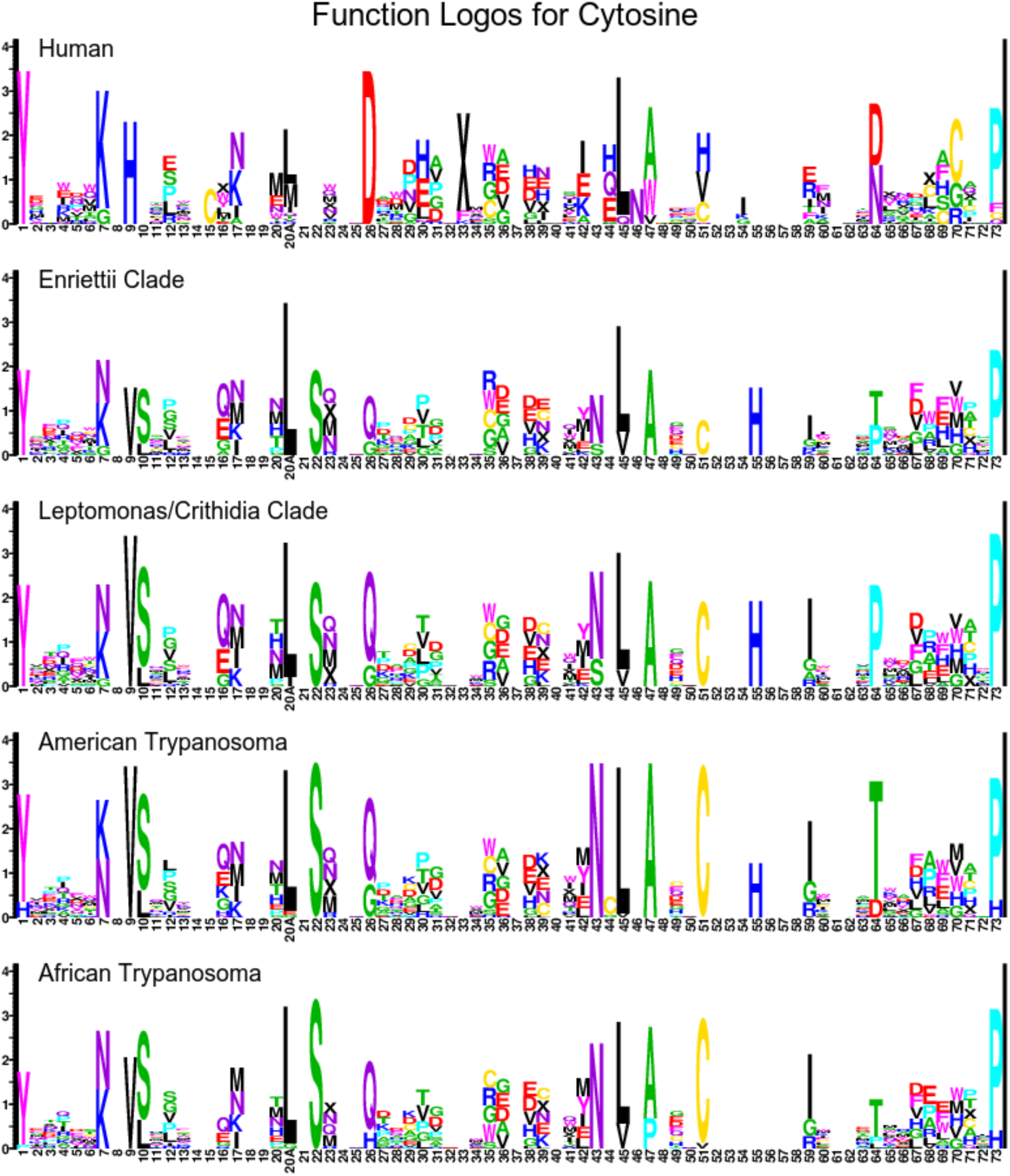
Cytosine Function Logos, part II

**Supplementary Figure 30.**
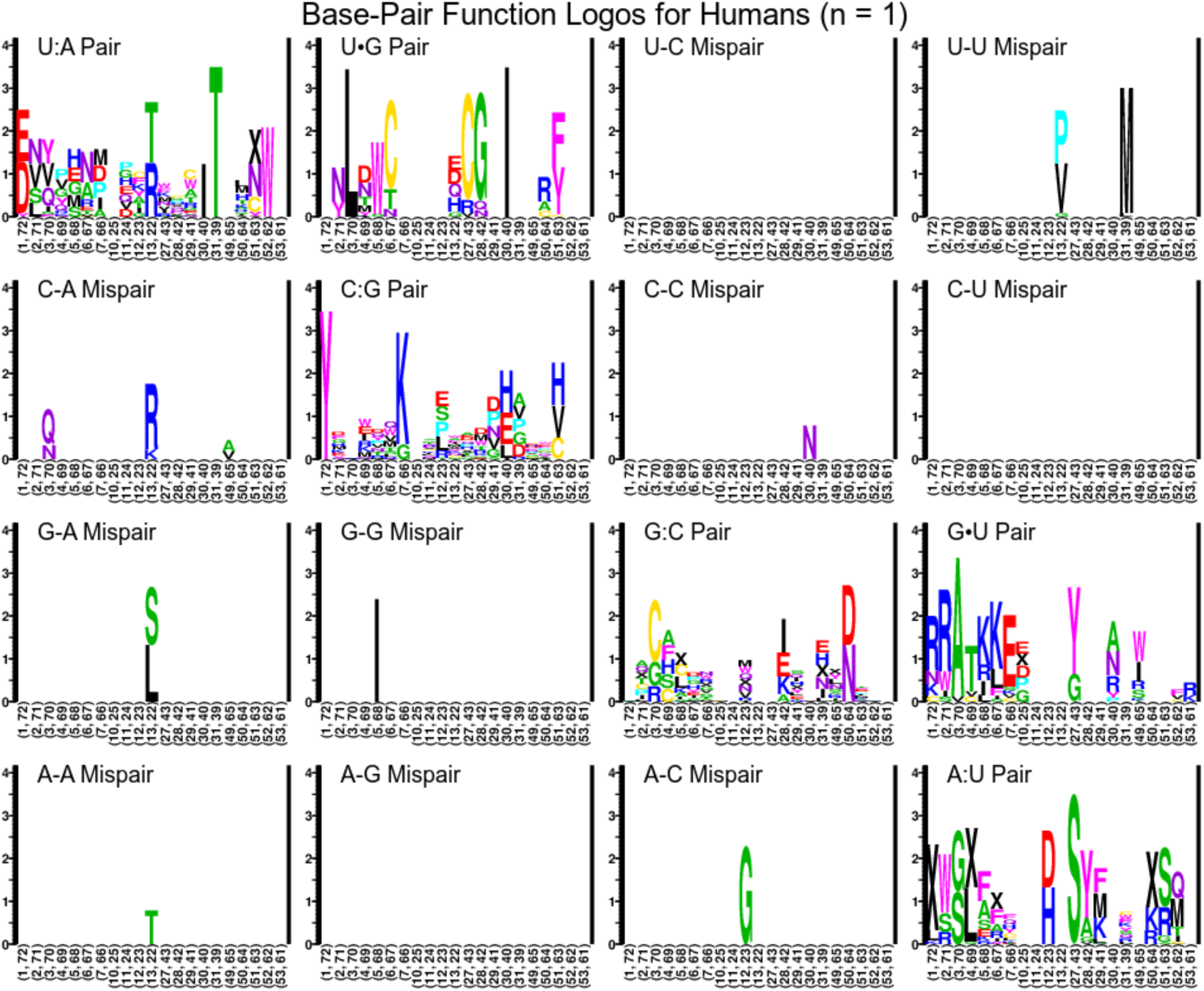
Paired-Site Function Logos for Humans

**Supplementary Figure 31.**
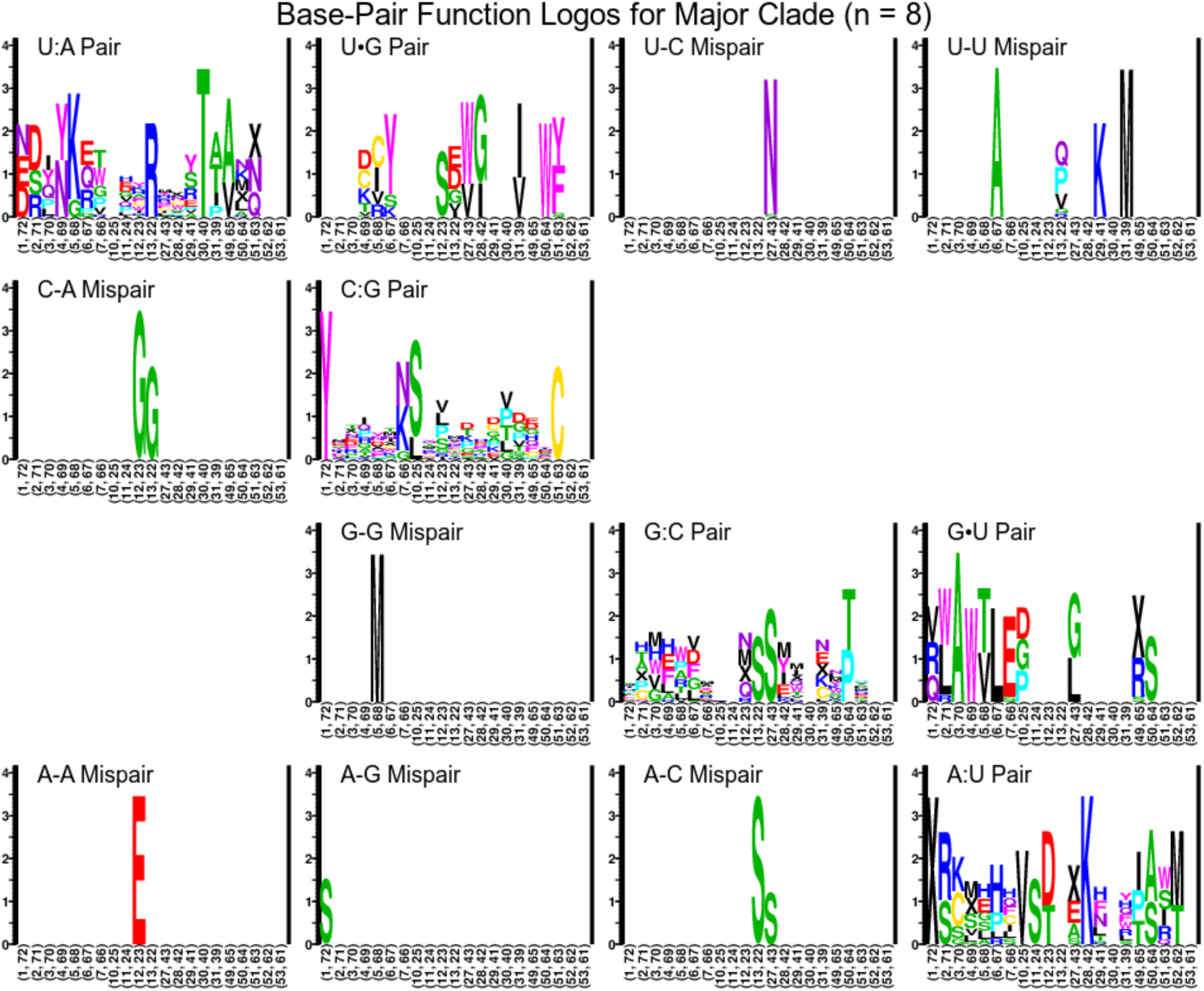
Paired-Site Function Logos for *L. Major* Clade

**Supplementary Figure 32.**
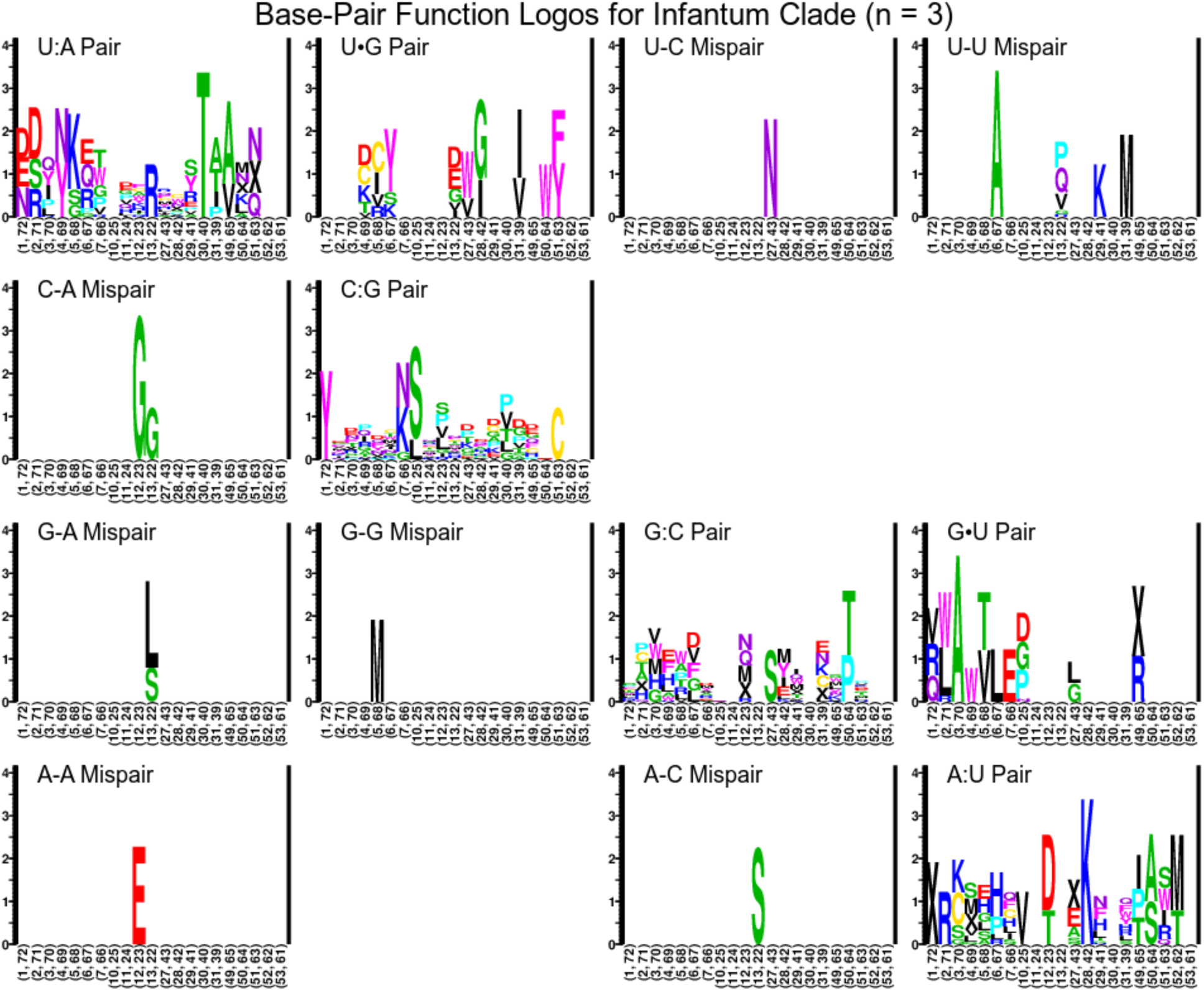
Paired-Site Function Logos for *L. Infantum* Clade

**Supplementary Figure 33.**
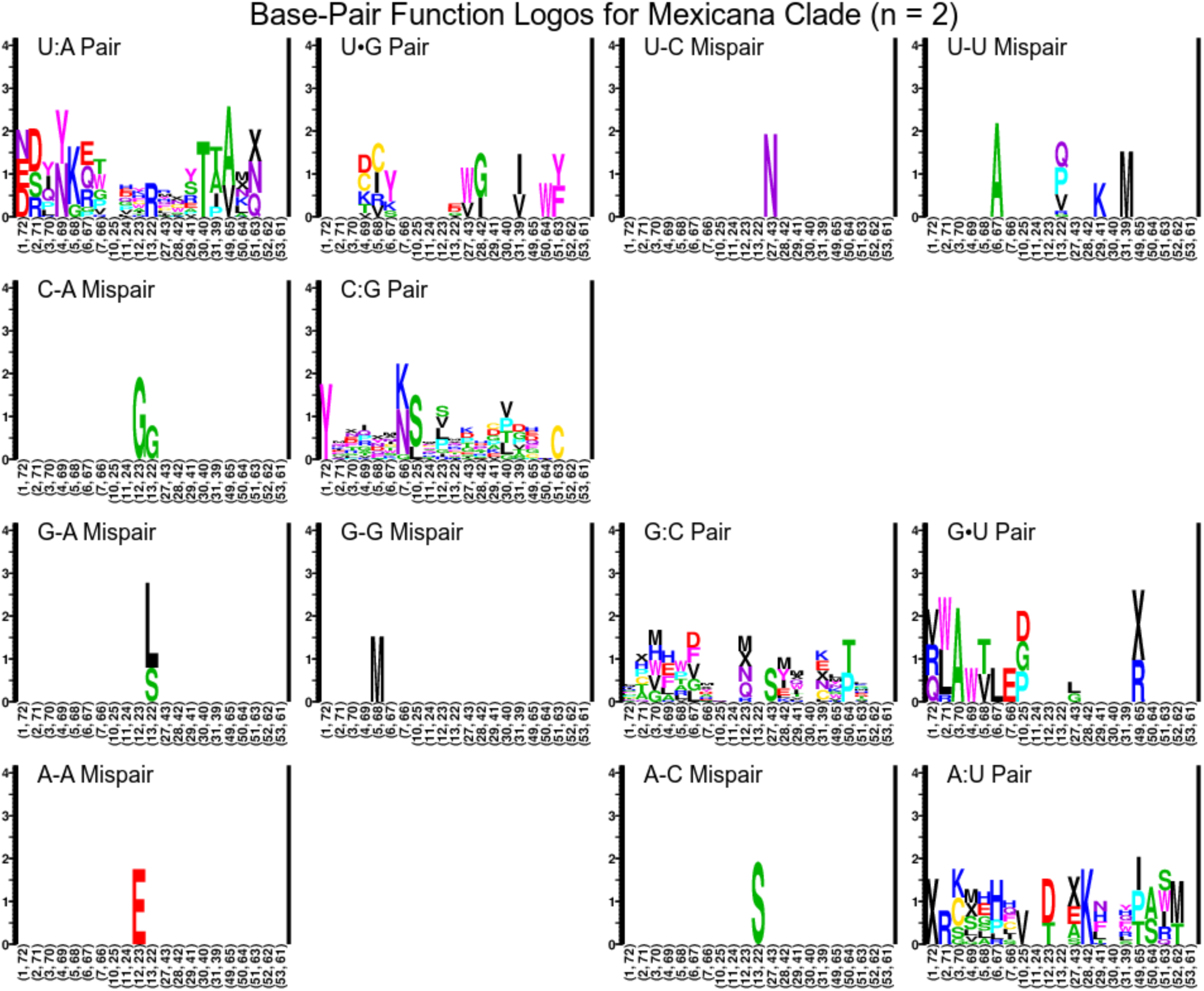
Paired-Site Function Logos for *L. Mexicana* Clade

**Supplementary Figure 34.**
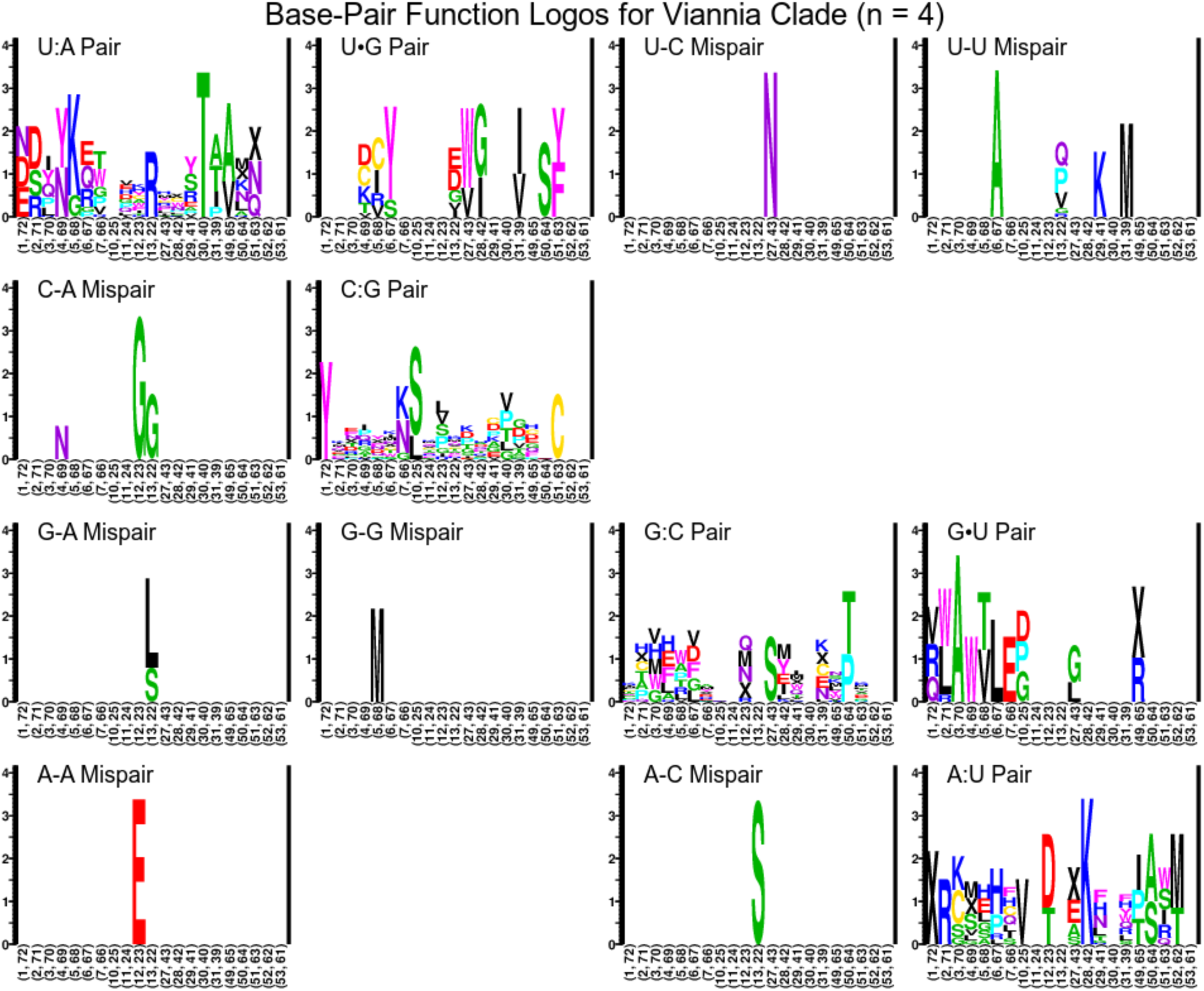
Paired-Site Function Logos for Viannia Subclade

**Supplementary Figure 35.**
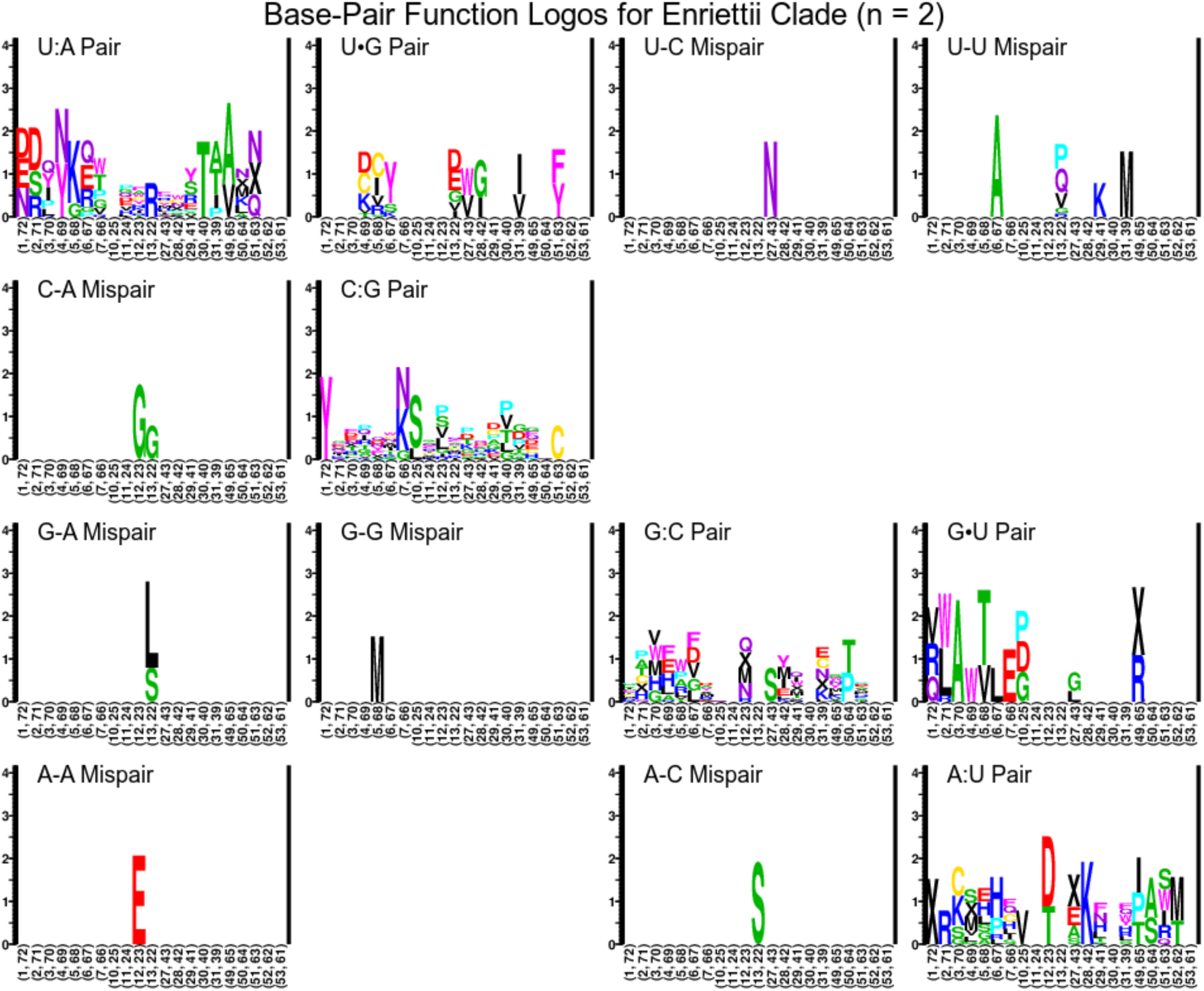
Paired-Site Function Logos for *L. Enriettii* Clade

**Supplementary Figure 36.**
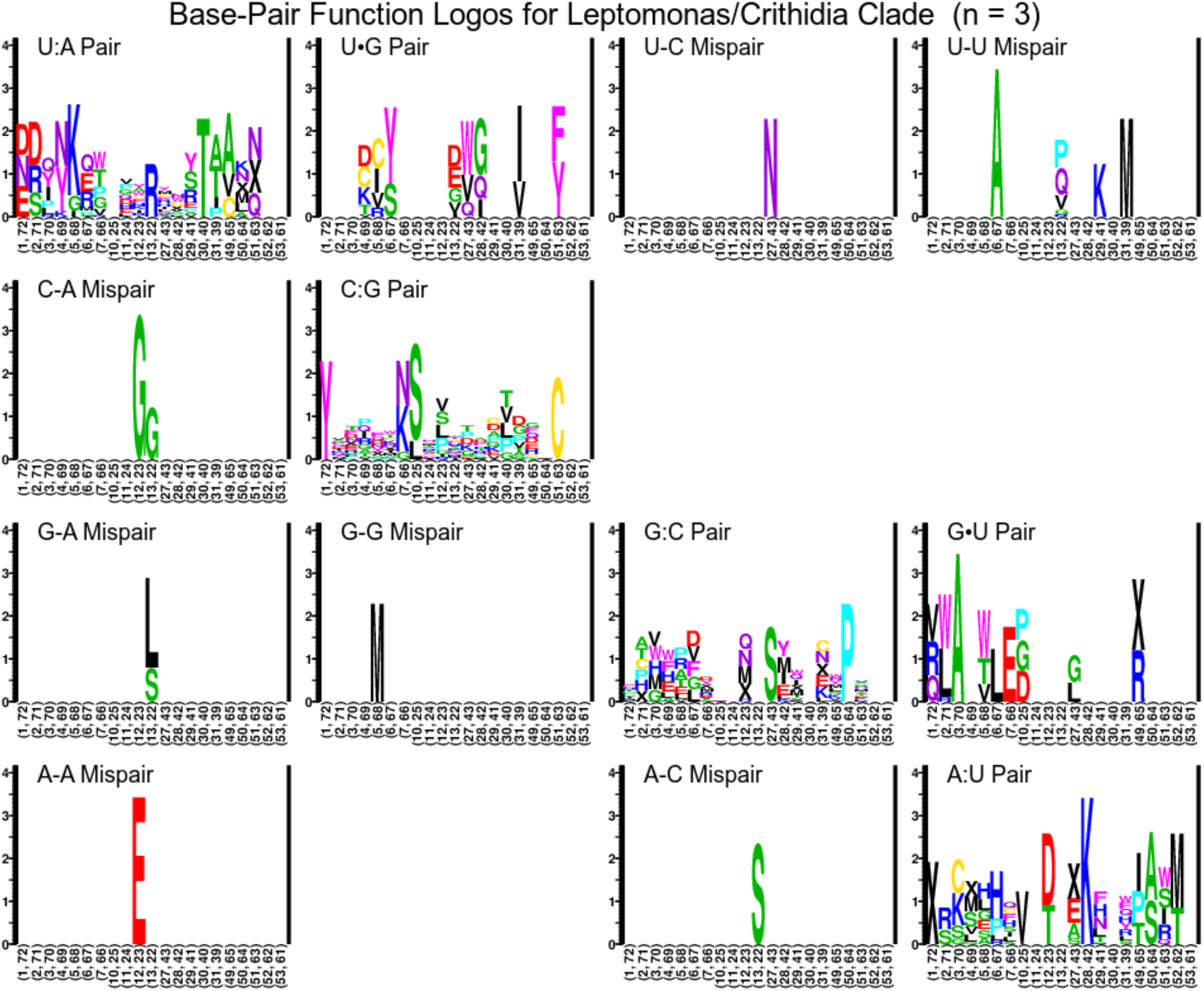
Paired-Site Function Logos for *Leptomonas*/*Crithidia* Clade

**Supplementary Figure 36.**
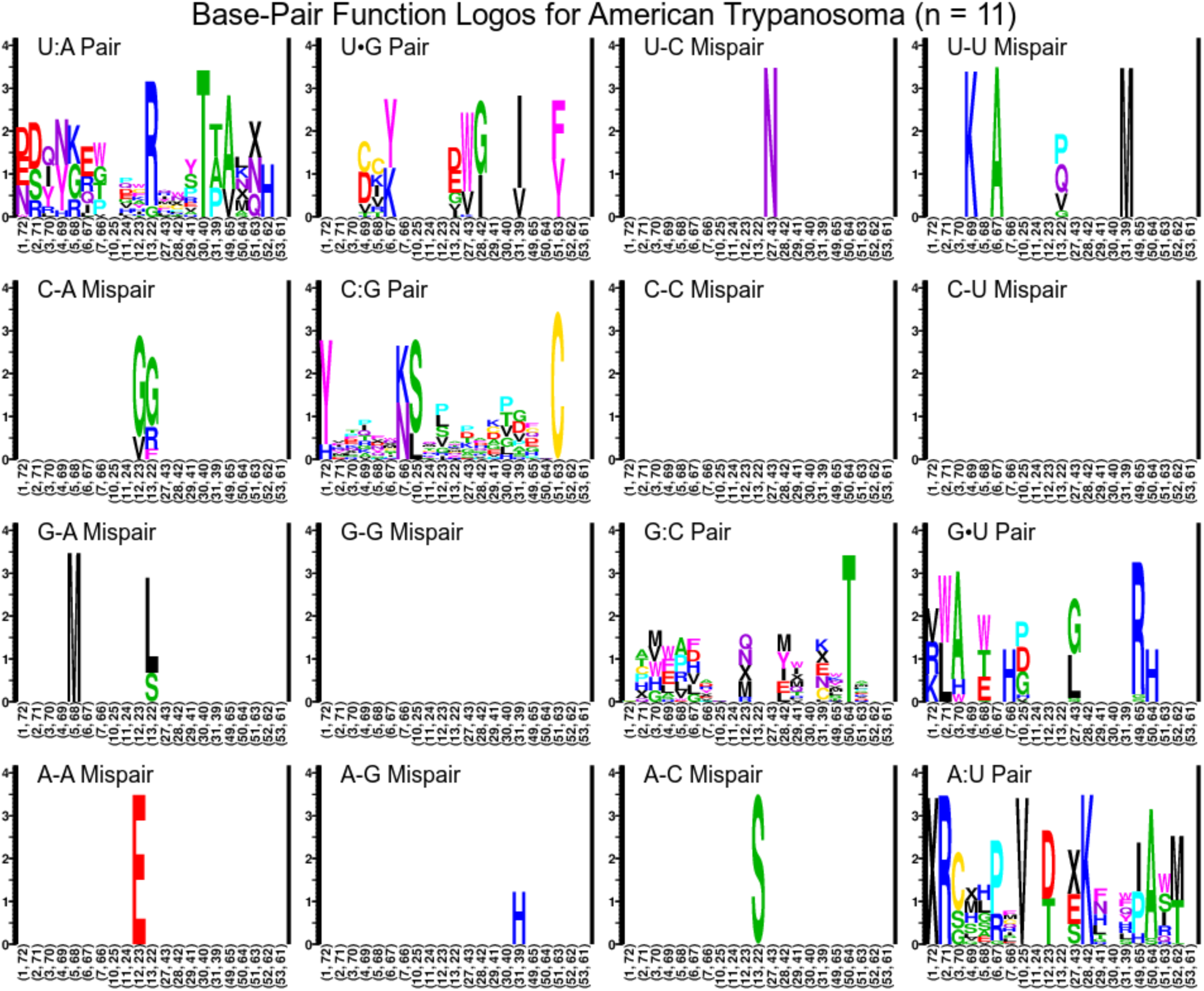
Paired-Site Function Logos for American *Trypanosoma* Clade

**Supplementary Figure 37.**
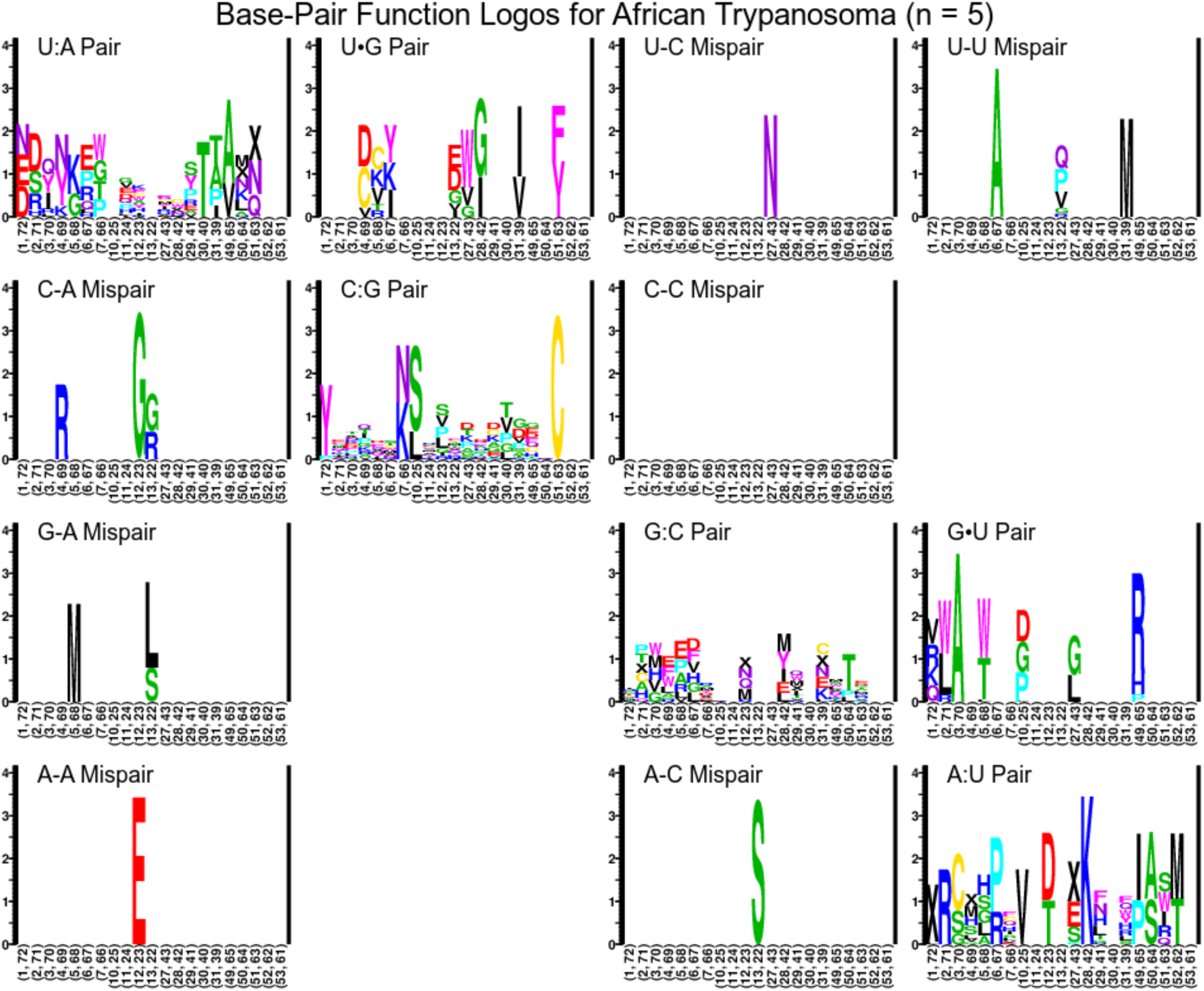
Paired-Site Function Logos for African *Trypanosoma* Clade

**Supplementary Figure 38.**
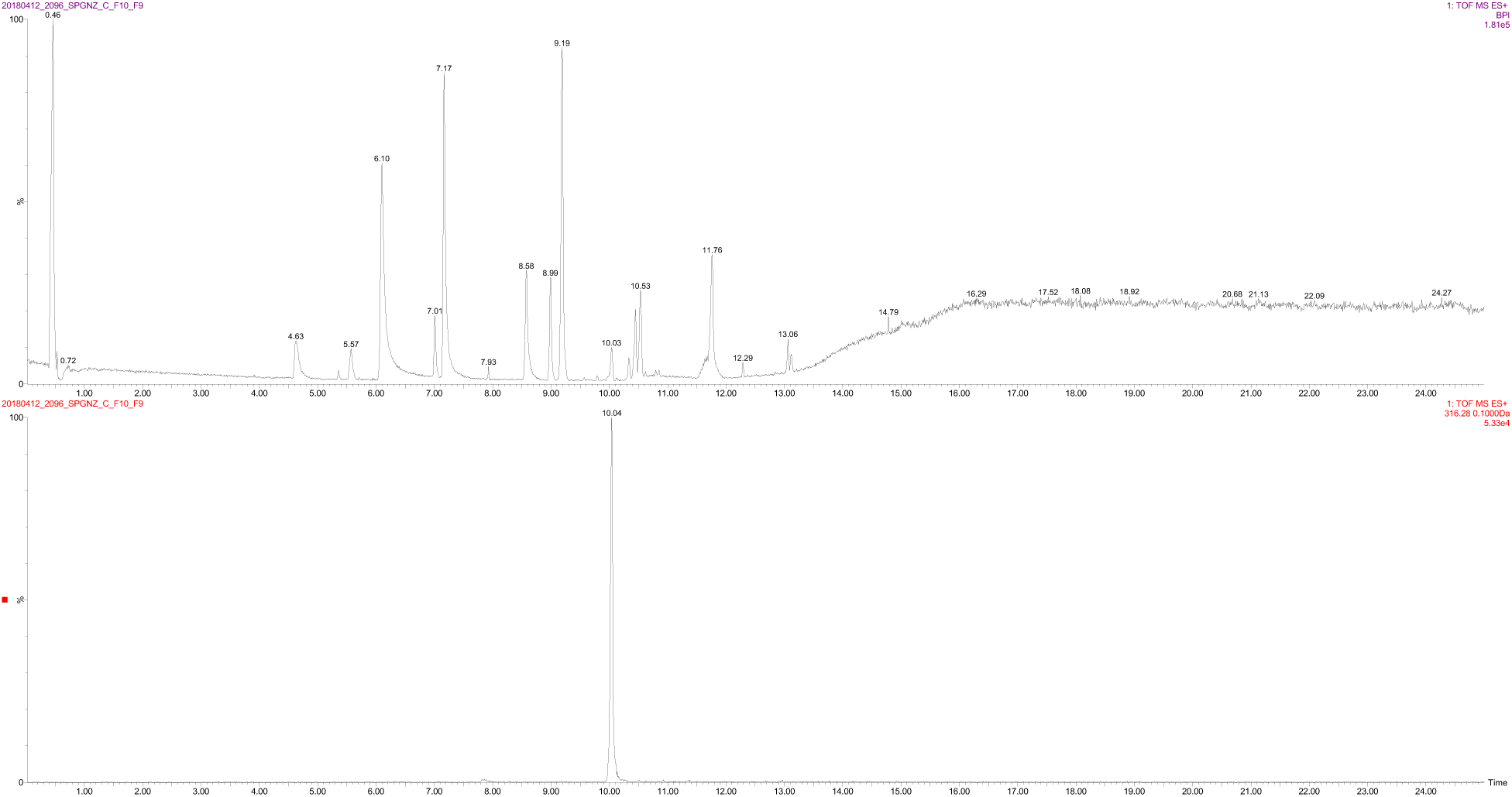
UPLC-qTOF base peak chromatogram of (A) RL12-182-HVF-D Sep-Pak Fraction C Subfraction 10-9 and (B) extracted ion chromatogram of peak 316.28 *m/z* in active region of trace.

**Supplementary Figure 38.**
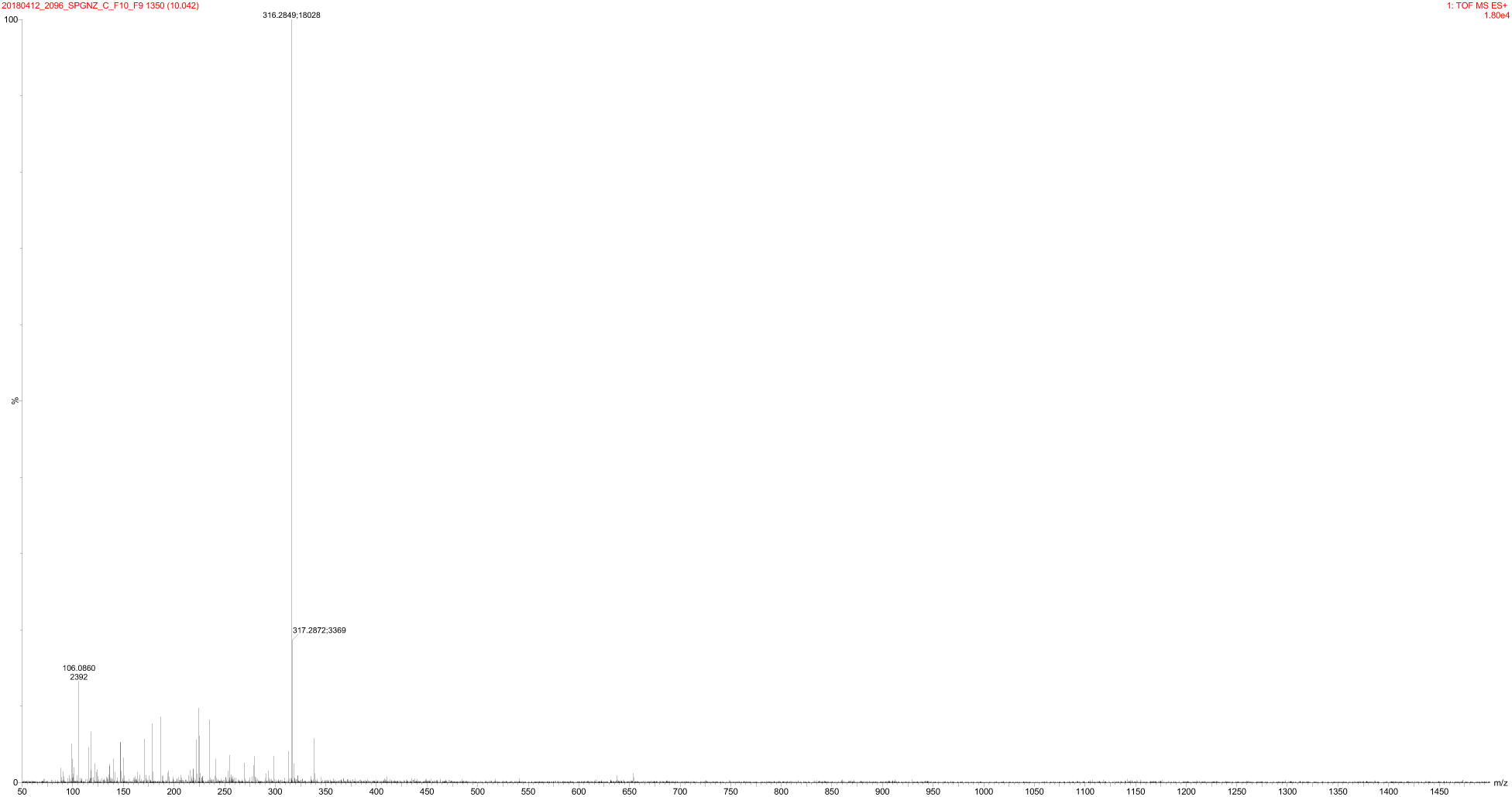
UPLC-qTOF mass spectrum for peak at 316.28 *m/z*.

**S40 Figure.**
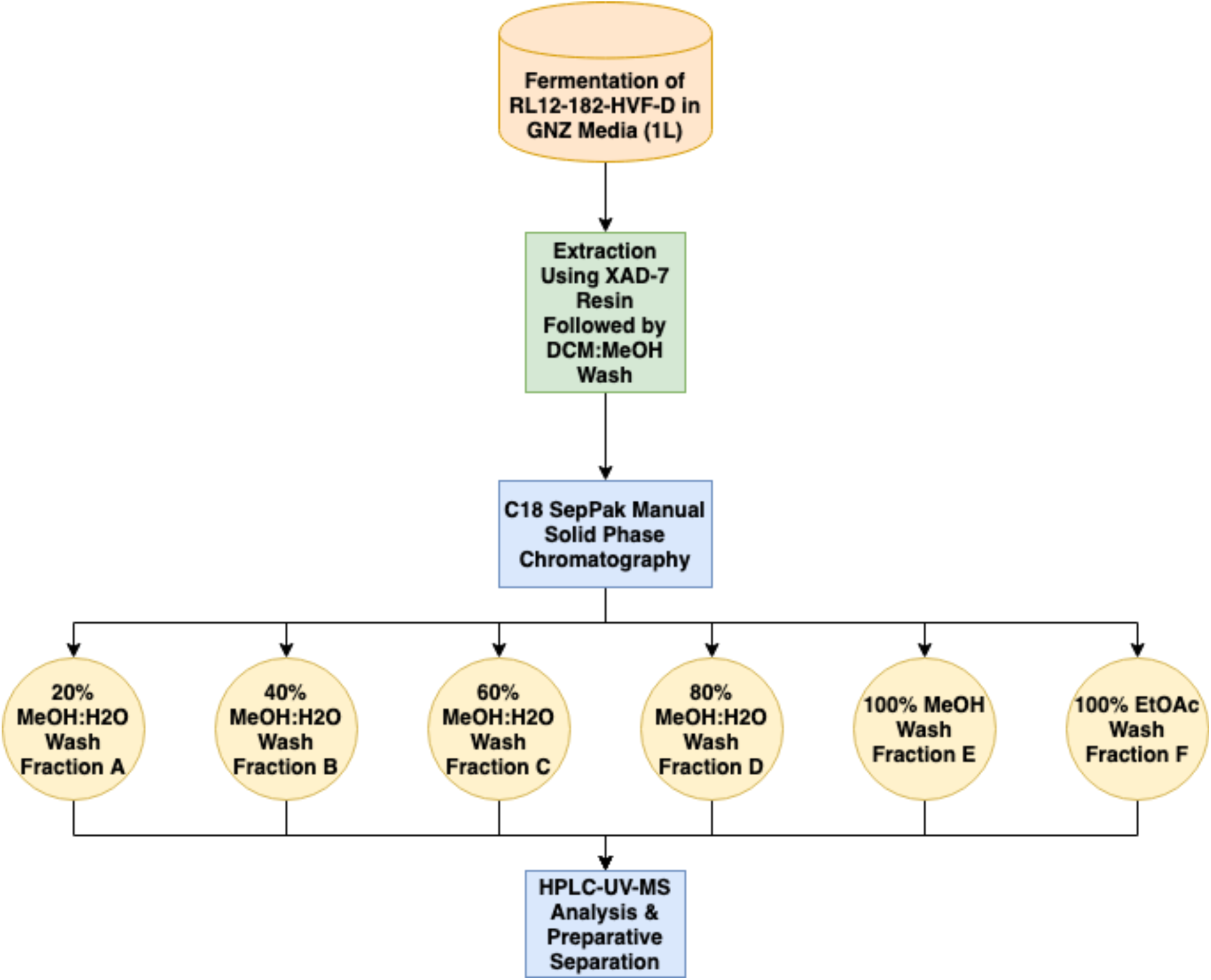
Schematic diagram of the large-scale fermentation and extraction of RL12-182-HVF-D.

**S41 Figure.**
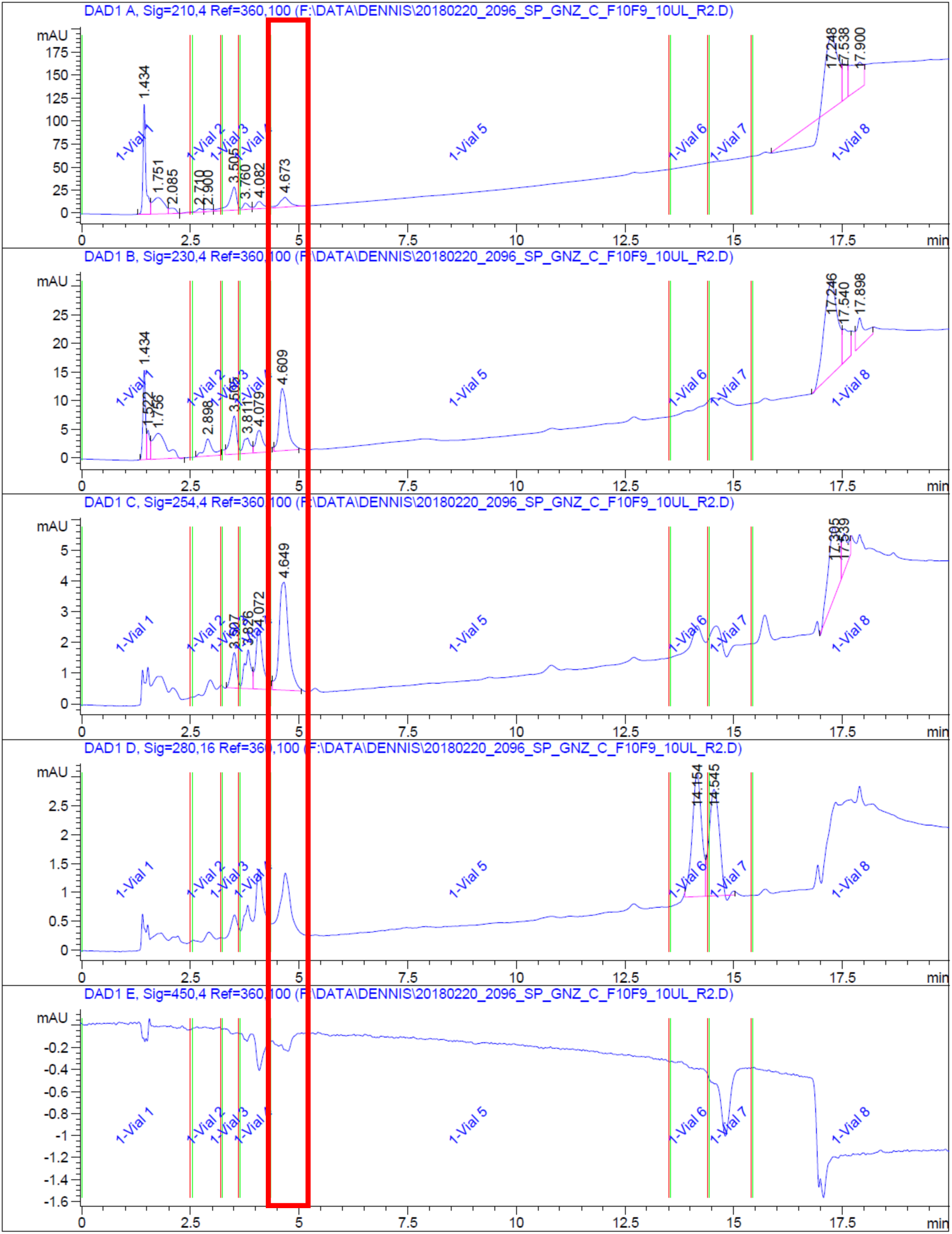
HPLC-UV trace of the final preparative isolation step for the RL12-182-HVF-D active compound. Red box denotes active peak.

**Supplementary Figure 42.**
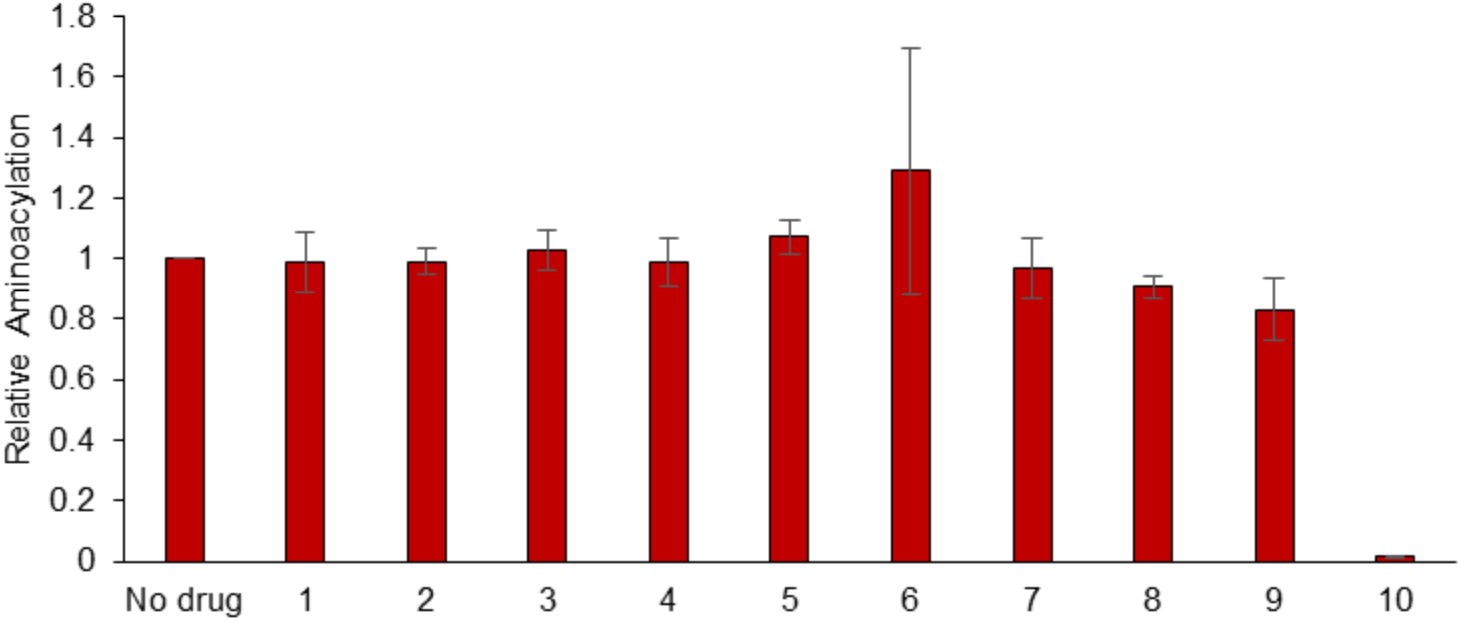
Marine natural product extract 2096D F10 inhibits *Leishmania major* AlaRS aminoacylation.

**Supplementary Table 1.**
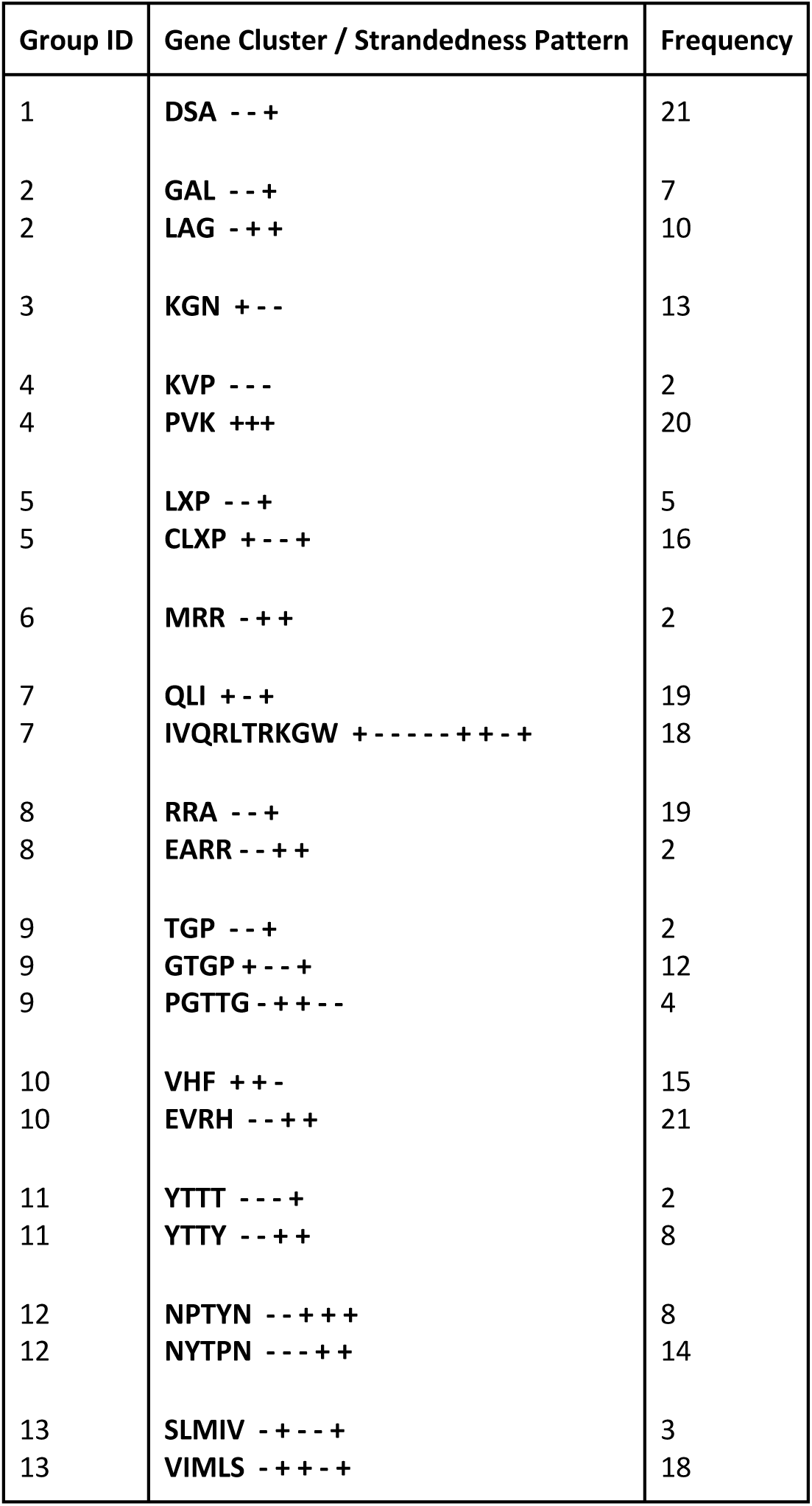
Putatively orthologous tRNA gene clusters of length at least 3 with frequencies in at least 2 *Leishmania* genomes. Clusters with the same group ID are considered similar.

**Supplementary Table 2.**
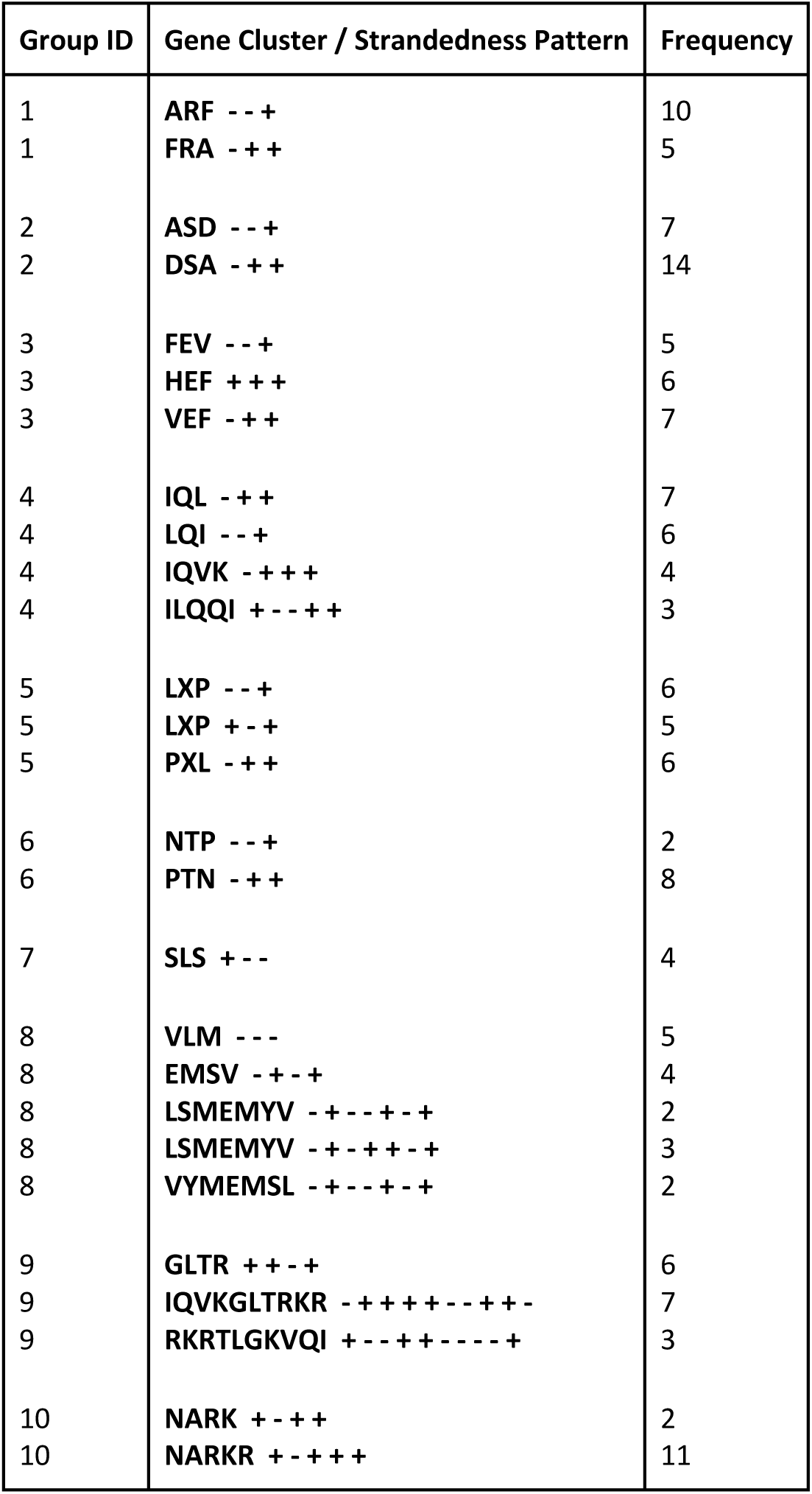
Putatively orthologous tRNA gene clusters of length at least 3 with frequencies in at least 2 *Trypanosoma* genomes. Clusters with the same group ID are considered similar.

**Supplementary Table 3.**
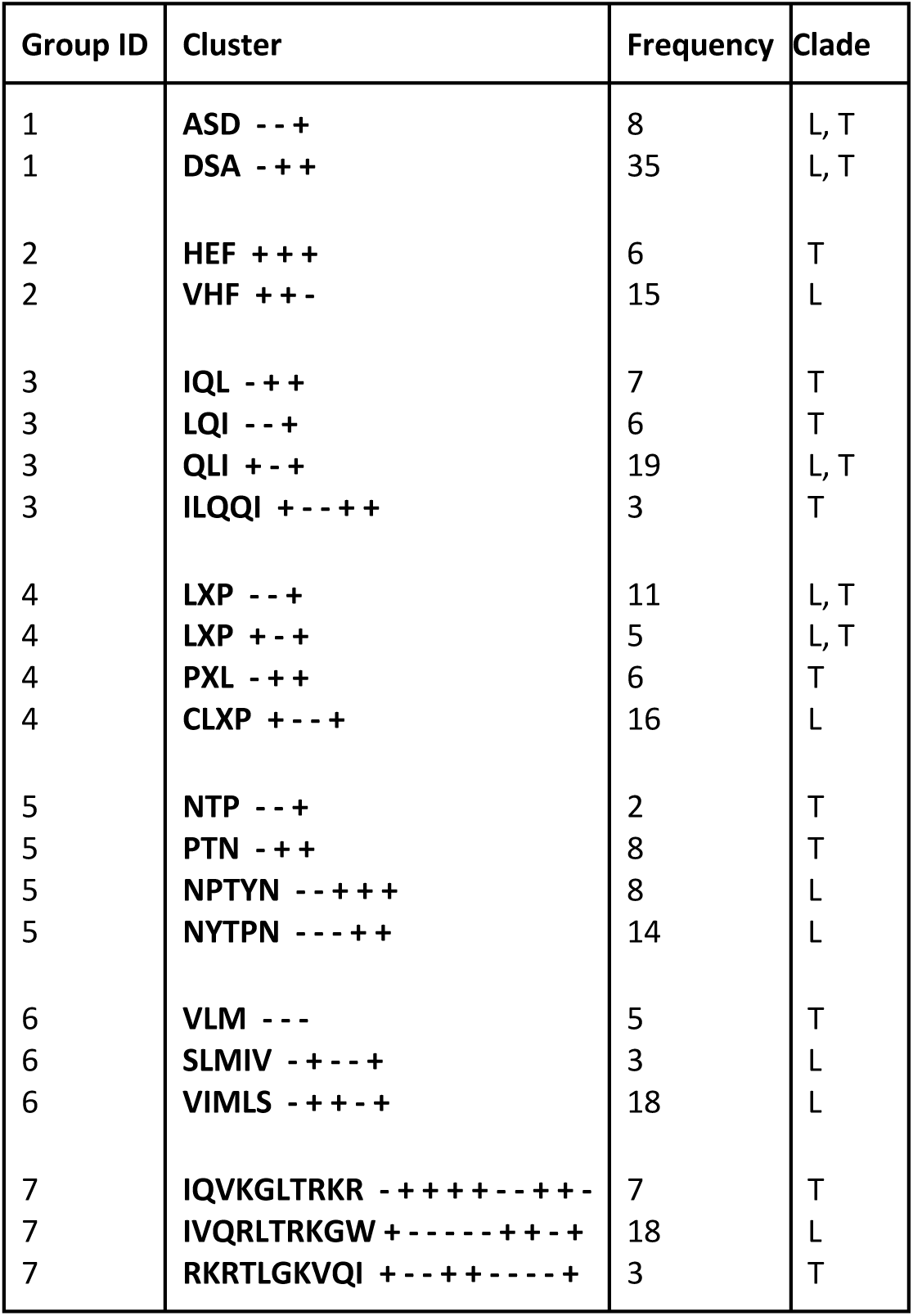
Putatively orthologous tRNA gene clusters of length at least 3 conserved across *Leishmania* (L) and *Trypanosoma* clade (T). Clusters ASD/DSA, LXP, abd QLI are the only clusters conserved in both clades. This table shows groups of similar clusters within both clades.

**Supplementary Table 4.** Statistics on Final Annotation Gene Set

|  | **Intersection Set** | **Aragorn-Only** | **Union Set** |
| --- | --- | --- | --- |
| **#tRNA genes** | 3579 | 36 | 3616 |
| **#genes/#genomes** | 74 | 98 | 75 |
| **Min. gene length** | 68 | 71 | 68 |
| **Max. gene length** | 89 | 206 | 206 |
| **%intron** | 2 | 28 | 3 |
| **%GC** | 58 | 59 | 58 |
| **#Ala tRNA genes** | 210 | 2 | 212 |
| **#Cys tRNA genes** | 64 | 1 | 65 |
| **#Asp tRNA genes** | 105 | 1 | 106 |
| **#Glu tRNA genes** | 160 | 1 | 161 |
| **#Phe tRNA genes** | 104 | 2 | 106 |
| **#Gly tRNA genes** | 228 | 3 | 231 |
| **#His tRNA genes** | 80 | 4 | 84 |
| **#Ile tRNA genes** | 171 | 1 | 172 |
| **#Lys tRNA genes** | 183 | 1 | 184 |
| **#Leu tRNA genes** | 335 | 6 | 341 |
| **#Met tRNA genes** | 97 | 0 | 97 |
| **#Asn tRNA genes** | 125 | 0 | 125 |
| **#Pro tRNA genes** | 200 | 0 | 200 |
| **#Gln tRNA genes** | 161 | 0 | 161 |
| **#Arg tRNA genes** | 348 | 2 | 350 |
| **#Ser tRNA genes** | 228 | 7 | 235 |
| **#Thr tRNA genes** | 218 | 4 | 222 |
| **#Val tRNA genes** | 236 | 0 | 236 |
| **#Trp tRNA genes** | 52 | 1 | 53 |
| **#iMet tRNA genes** | 76 | 0 | 76 |
| **#Tyr tRNA genes** | 88 | 0 | 88 |
| **#SeC tRNA genes** | 76 | 0 | 76 |
| **#Ambiguous tRNA genes** | 34 | 0 | 35 |

## Supplementary Text 1 — Supplementary methods for bacterial fermentation, natural product extraction, and active compound identification using HPLC-UV-MS and UPLC-ESI-qTOF-MS

### Bacterial Fermentation and Natural Product Extraction

Frozen stocks of the associated producing organism RL12-182-HVF-D was streaked onto fresh Marine Broth agar plates (Difco, USA) and incubated at room temperature (∼ 25 lllC) until discrete colonies became visible. Individual colonies were inoculated into 7 mL (small-scale) of modified saline SYP (mSYP) media (10 g starch, 4 g peptone, 2 g yeast extract and 31.2 g instant ocean in 1 L of distilled water) or GNZ media (20 g starch, 10 g glucose, 5 g NZ-amine, 1 g CaCO3, 5 g yeast extract in 1 L of distilled water). Bacterial fermentation was stepped up in stages by inoculating 3 mL of the 7 mL liquid mSYP or GNZ culture into 60 mL (medium-scale) of mSYP or GNZ, respectively. This was followed by inoculating 30 mL of the medium scale culture into 1 L (large-scale) in respective media with 20 g of pre-washed XAD-7 resin (CH2Cl2, MeOH and water). Small-scale cultures were incubated for four days, medium-scale cultures for four days and large-scale cultures for seven days, at ∼ 25 lllC and shaken at 200 RPM.

Large-scale cultures were extracted by first filtering the cellular/resin slurry under vacuum through two layers of Whatman filter paper. The cells, resin and filter paper were extracted twice with 500 mL of 1:1 CH2Cl2:MeOH and the suspension stirred for 1 hour. Combined organic extracts were filtered and concentrated to dryness in vacuo. Dried crude extracts for each media (mSYP and GNZ) were fractionated individually by manual solid phase extraction chromatography using Sep-Pak (SP) columns (5 g C18 cartridge, Supelco, USA). Chromatography proceeded using a stepwise MeOH/H2¬O gradient: 40 mL of 10% MeOH (wash), 20% (fraction A), 40% (fraction B), 60% (fraction C), 80% (fraction D), 100% (fraction E) then 100% EtOAc wash (fraction F). SP fractions A – F were concentrated to dryness in vacuo and prepared for HPLC-UV separation.

### HPLC-UV-MS and UPLC-ESI-qTOF-MS Analyses

All high-performance liquid chromatography analyses were performed on an Agilent 1200 series HPLC system equipped with both an Agilent photodiode array (PDA) detector and an Agilent 6130 single quadrupole mass spectrometer to acquire UV and MS data respectively. Samples were injected onto a C18 reverse-phase column (Synergi 10lll Fusion RP Column, Phenomenex, USA) using a H2O:MeOH (0.02% formic acid) elution profile: 0 – 3 mins, 5% MeOH; 3 – 25 min, linear gradient 5% to 100% MeOH; 25 – 30 min, isocratic at 100% MeOH; 30 – 35 min, isocratic at 5% MeOH, using a flow rate of 2 mL/min and positive ESI mode.

Accurate mass MS/MS data were acquired on an Acquity i-Class UPLC system (Waters Corporation) with SYNAPT G2-Si qTOF mass spectrometer (Waters Corporation) run in HRMS positive ESI mode. The instrument was operated using a 20 lllg/mL leucine enkephalin lockspray infusion injected every 10 seconds to control mass accuracy. Samples are injected onto a C18 reverse-phase column (HSS C18, 100 mm x 2.1 mm, 1.7lllm, Waters Corporation) using a H2O (0.1% formic acid):ACN (0.1% formic acid) elution profile: 0 – 0.3 min, 5% ACN; 0.3 – 4.7 min, linear gradient 5% to 90% ACN; 4.7 – 5.5 min, linear gradient 90 to 98% ACN; 5.5 – 5.8 min, isocratic at 98% ACN; 5.81 – 7.5 min, isocratic at 5% ACN, using a flow rate of 0.5 mL/min.

